# Fine mapping and genomic analyses reveal a tandem kinase-pseudokinase candidate underlying *Or_Deb2_*-mediated resistance to *Orobanche cumana* in sunflower

**DOI:** 10.64898/2026.02.12.705495

**Authors:** Belén Fernández-Melero, Lidia del Moral, Borja Rojas-Panadero, Alexandra Legendre, Sébastien Carrère, Jérôme Gouzy, Stéphane Muños, Leonardo Velasco, Begoña Pérez-Vich

## Abstract

Sunflower broomrape (*Orobanche cumana* Wallr.) is a holoparasitic weed that parasitizes sunflower (*Helianthus annuus* L.) and severely constrains yield. The *Or_Deb2_* gene confers resistance to broomrape race G in the sunflower line DEB2. It has been previously located within a 1.38-Mbp physical region in the *H. annuus* XRQ reference genome that contains a cluster of receptor-like proteins and receptor-like kinases (RLKs). In this study, an *Or_Deb2_* candidate has been proposed though high-resolution genetic and physical mapping, multi-allelic comparisons across resistant and susceptible lines, and exclusion of other potential candidates based on their function and genetic features. *Or_Deb2_* was delimited to a 219.4-kbp region in the DEB2 genome assembly that contained six annotated protein-coding genes belonging to two groups: three small heat shock proteins (HSP20) and three RLKs. Given that the only sunflower broomrape resistance gene cloned to date encodes an RLK, the key role of RLKs in resistance to biotic stresses in plants, and the similar genetic features between RLK-type resistance genes and *Or_Deb2_* as major dominant genes in gene-for-gene interactions, RLKs in the interval emerged as the most plausible candidates. Within them, the *RLK-2* gene encoding a root-expressed full-length kinase-pseudokinase protein belonging to the tandem kinase proteins (TKPs) family was prioritized, because the other two RLKs corresponded to a truncated TKP with a stop codon within the kinase domain. *RLK-2* also carried sequence variants unique to DEB2 relative to multiple susceptible genotypes. The results of this research reveal the potential role of TKP genes as new players in resistance to parasitic plants and provide a framework to dissect the genetic dialog between *O. cuma*na and sunflower in gene-for-gene interactions.

## Introduction

Sunflower broomrape (*Orobanche cumana* Wallr.) is a holoparasitic plant that lacks photosynthetic capacity and relies entirely on its host to complete its life cycle. It establishes vascular connections with sunflower (*Helianthus annuus* L.) roots, through which it withdraws water and nutrients, thereby ensuring its growth and reproduction. This parasitism causes severe yield losses in sunflower crops across many producing regions, from Southeastern Europe to Central Asia and China (Fernández-Martínez et al., 2015), and more recently in North Africa (Nabloussi et al., 2018). Furthermore, broomrape is expanding into the previously unaffected Americas, with a recent first report on sunflower crops in Bolivia (Barea et al., 2025), posing a growing threat to sunflower production worldwide.

Developing resistant sunflower cultivars is the main and most sustainable way to control broomrape parasitism (Cvejić et al., 2020). Generally, the genetic interaction between sunflower and broomrape follows the gene-for-gene model proposed by Flor (1971), in which a dominant allele for resistance in the host interacts with a dominant allele for avirulence in the parasite (Velasco et al., 2016; Lebedeva et al., 2023). This type of interaction leads to the emergence of different physiological races of the parasite, which can evolve in response to the host’s resistance mechanisms (Imerovski et al., 2016). As of now, several races of broomrape, categorized from A (least virulent) to H (most virulent) have been identified. For many of these races, a sunflower-dominant resistance gene has been identified and designated using the *Or_x_*or *HaOr_x_* nomenclature (Cvejić et al., 2020). Thus, it has been described the *Or_1_* to *Or_5_*genes conferring resistance to races A to E, respectively (Vrânceanu et al., 1980); the *Or_6_* (Pacureanu-Joita et al., 1998) and *HaOr_7_*(Duriez et al., 2019) controlling race F; and the *Or_SII_* (Martín-Sanz et al., 2020), *Or_Deb2_* (Velasco et al., 2012), and *Or_Anom1_* (Fernández-Melero et al., 2024) for race G. Nevertheless, in addition to these dominant genes, other types of inheritance of the broomrape resistance trait have also been described in sunflower, such as a single recessive gene (Imerovski et al., 2016), two recessive genes (Akhtouch et al., 2002, 2016) or multiple quantitative trait loci (QTL) (Imerovski et al., 2019; Louarn et al., 2016; Pérez-Vich et al., 2004).

Various genetic approaches have enabled the identification of genomic regions in sunflower that contain QTL and major genes for resistance to *O. cumana* (Cvejić et al., 2020). Concerning major genes, *Or_5_*has been mapped by several authors to the telomeric upper part of sunflower chromosome (Chr) 3 (Tang et al., 2003; Pérez-Vich et al., 2004), and is located in a genomic region of 193-kbp containing putative resistance genes encoding F-box/LRR (leucine-rich repeat)-repeat proteins or a TIR (Toll/interleukin-1 receptor)-NLR (nucleotide-binding leucine-rich repeat) protein (Pubert et al., 2024). The genes *Or_SII_* (Martín-Sanz et al., 2020), *Or_Deb2_* (Fernández-Aparicio et al., 2022), and *Or_Anom1_* (Fernández-Melero et al., 2024) have been mapped to a tight region of Chr 4, which houses a cluster of receptor-like kinase (RLK) and LRR-receptor-like protein (RLP) genes. These genes are characterised by lacking a cytoplasmic kinase domain (LRR-RLPs) or, alternatively, having one or two fused kinase domains and lacking an extracellular LRR region (RLKs) (Fernández-Aparicio et al., 2022). Finally, the *HaOr_7_* gene, mapped to Chr 7, is the only one cloned to date. It encodes an LRR-RLK, having an extracellular LRR domain and an intracellular kinase domain (Duriez et al., 2019).

Wild *Helianthus* species are a major reservoir of genes controlling resistance to the new virulent races of *O. cumana* (Seiler and Jan, 2014). The *Or_Deb2_* gene has been introgressed from a resistant accession of the wild *Helianthus debilis* subsp. *tardiflorus* (Velasco et al., 2012), and it is highly valuable for broomrape resistance breeding as it confers broad resistance to several *O. cumana* races. More specifically, this gene controls broomrape populations of races E, F and G (Martín-Sanz et al., 2016; Fernández-Aparicio et al., 2022), although populations surpassing *Or_Deb2_* resistance have been recently reported in central Spain and classified as race F^+^_CU_ (Fernández-Melero et al., 2023). In any case, *Or_Deb2_* is one of the most valuable resistance genes currently available for breeding for resistance to broomrape in sunflower, and its identification is key to understanding the molecular interactions between resistance and avirulence proteins in the gene-for-gene sunflower-*O. cumana* pathosystem. The objectives of this study were the genetic dissection and fine mapping of the *Or_Deb2_*locus. By combining high-resolution genetic and physical mapping, study of multiple alleles from resistant and susceptible lines, and exclusion of other potential candidates by their function and genetic features analyses, we aimed to propose an *Or_Deb2_* candidate as an RLK gene encoding a tandem kinase-pseudokinase protein.

## Materials and Methods

### Plant material and phenotypic evaluation

DEB2 is a sunflower line resistant to populations of race G of sunflower broomrape in which the resistance gene, named *Or_Deb2_*, was introgressed from a resistant accession of the wild species *H. debilis* subsp. *tardiflorus* (Velasco et al., 2012). HA821 is an inbred sunflower line released by the USDA-ARS and North Dakota State Agricultural Experimental Station in 1983 (Roath et al., 1986) that is susceptible to sunflower broomrape (Velasco et al., 2007). B117 is a sunflower confectionery landrace collected in Spain that is susceptible to sunflower broomrape (Martín-Sanz et al., 2016).

Fine mapping of *Or_Deb2_* was conducted by genotyping a population of 3696 F_2_ plants derived from crosses of DEB2 with B117 and HA821 (with 86% of the F_2_ plants from the B117 x DEB2 cross). The F_2_ plants were evaluated for resistance to broomrape race G in 2015 (HA821 x DEB2) and 2020 (B117 x DEB2) as described below. All F_2_ plants were self-pollinated to obtain F_3_ seed. F_2:3_ families (F_3_ plants) from selected recombinant F_2_ plants were evaluated for resistance to broomrape race G in the winter 2020/21 and 2023/24 to confirm susceptible phenotypes and to differentiate between homozygous and heterozygous F_2_ resistant plants. Since all recombinants identified came from the B117 x DEB2 cross, the susceptible line B117 was selected over HA821 for marker development and sequencing.

Evaluation of broomrape resistance was carried out in pots using a sunflower broomrape population of race G collected in 2000 in Çeşmekolu, Kirklareli Province, Turkey, (race G_TK_ according to Martín-Sanz et al. 2016 race nomenclature), named as GT. Sunflower seeds were germinated in moistened filter paper and sown in small pots (7 × 7 × 7 cm) in which 50 mg of GT broomrape was added to a mixture of sand and peat (1:1, by vol.). The plants were kept in a growth chamber for 20-25 days at 25 °C/20 °C (day/night) using a 16 h photoperiod and then transplanted to larger pots containing 6 L of fertilized soil mixture made of sand, silt, and peat (2:1:1, by vol.), and 8 g of NPK controlled release fertilizer Nutricote 15-9-10 (2MgO)+ME (Jcam Agri. Co., Ltd., Tokyo, Japan). Evaluation for broomrape resistance was done by the end of the sunflower flowering stage by counting emerged broomrape plants. Sunflowers were classified as resistant if they showed no emerged broomrape plants and susceptible if they showed emerged broomrape plants. In all cases, the susceptible parents, B117 and HA821, and the resistant parent, DEB2, were grown as controls. Also, the two sunflower lines XRQ and PSC8, from which the two sunflower reference genomes have been obtained, were tested for broomrape resistance to the GT population under open-air conditions in the spring of 2023. Phenotyping was carried out under greenhouse conditions for the winter sowings and under open-air conditions for the spring sowings, using the facilities available at the Institute for Sustainable Agriculture (IAS-CSIC, Córdoba, Spain).

### Tissue collection, DNA extraction, and plant genotyping

Two fully expanded young leaves were cut from each F_2_ or F_3_ plant, frozen at -80 °C, lyophilized, and ground to a fine powder in a laboratory mill. The DNA was extracted using the innuPREP Plant DNA kit (IST Innuscreen, Berlin, Germany) according to the manufacturer’s instructions.

Genotyping was performed in different consecutive rounds (rounds 1 to 5, summarized in Figure S1). (1) In the first round, a total of 309 susceptible F_2_ plants derived from the DEB2 × B117 and DEB2 × HA821 crosses were genotyped with 10 SNPs. These included three previously described Axiom SNP markers flanking *Or_Deb2_* (AX-105412178, AX-105399507, AX-105769027; Fernández-Aparicio et al., 2022) and seven newly developed SNP markers (Iasnip-15 to Iasnip-24). These seven novel SNPs were designed, as detailed in the following section, using cloned B117 and DEB2 fragments amplified with primers derived from annotated genes located within the 1.38-Mbp *Or_Deb2_* window described by Fernández-Aparicio et al. (2022). (2) From this first round, thirty-six informative recombinants that reduced the *Or_Deb2_* window were identified, used to develop 14 new SNP markers (Iasnip- 26 to 55) also based on cloned B117 and DEB2 fragments, and genotyped with former and the newly developed markers. (3) In the third round, larger F_2_ sets were analysed: 3a) 803 F_2_ plants from DEB2 × B117 and DEB2 × HA821 crosses, genotyped with 24 SNPs; and 3b) 2584 F_2_ plants from DEB2 × B117, genotyped with 10 selected SNPs. 4) From this last set, sixty-eight informative recombinants were further genotyped with two co-dominant PCR-based markers (FMDEB-57/58 and FMDEB-59/60; FLPs - Fragment Length Polymorphism markers), further narrowing the *Or_Deb2_* interval. (5) These different genotyping rounds allowed the final selection of five highly informative recombinants named R1 to R5.

This recombinant set, together with its derived F_2:3_ families (F_3_ plants), was then genotyped using both SNP and FLP markers described above, as well as additional PCR-based markers based on the DEB2 genome sequence, as explained in the next section. SNPs were genotyped by competitive allele-specific PCR assays based on KASP^TM^ technology (LGC genomics, Teddington, Middlesex, UK). PCR-based markers were genotyped as described below. Nucleotide sequences of the most informative markers used in the different genotyping rounds are provided in Table S1.

### Marker development based on the sequence of B117 and DEB2 cloned fragments, and on the DEB2 genome assembly

Two marker types were developed for fine mapping *Or_Deb2_*. The first type was SNP markers based on the sequence of cloned B117 and DEB2 fragments (markers Iasnip- in the interval from 15 to 55) (Table S1). For that, PCR primers were designed primarily in coding regions of the reference sunflower genome HanXRQr2.0 (https://www.heliagene.org/HanXRQr2.0-SUNRISE) and used to amplify the parental lines B117 and DEB2 in order to obtain cloned sequences and identify SNPs. PCR reactions were performed in 50 µl using 1.25 units of MyTaq DNA polymerase (Bioline, Meridian Life Science, Memphis, USA), 1x MyTaq reaction buffer (containing 5 mM dNTPs and 15 mM MgCl2), 0.4 µM of each primer, and 100 ng of genomic DNA. The amplification was performed in a GeneAmp PCR system 9700 (Applied Biosystem, Foster City, CA, USA) with an initial denaturation step of 3 min at 95 °C followed by 34 cycles of 30 s at 95 °C, 30 s at annealing temperature (48-58 °C) and 1 min/kb at 72 °C, followed by an extension step of 20 min at 72 °C to create a poli-A tail. PCR products were analysed by electrophoresis in an agarose gel. Non-polymorphic products were purified using the MEGAquick-spin Plus Total Fragment DNA Purification kit (Intron Biotechnology, Inc, Korea) and cloned into the pSpark TA Done DNA cloning vector as described by the manufacturer (Canvax Biotech SL, Córdoba, Spain). Sequences were obtained by Sanger sequencing (Stab Vida, Lisbon, Portugal) and analysed to identify SNPs between the two lines. Sequences and their identities were confirmed using BLAST. Sequence analysis was conducted using the Lasergene SeqMan Ultra and MegAlign Pro software within the DNASTAR, Inc. package.

The second marker type was PCR-based. These included: (i) FLPs of primary PCR fragments (co-dominant FMDEB-57/58 and FMDEB-59/60 markers) (Table S1), designed inside the fine-mapped region in coding regions from the reference sunflower assembly HanXRQr2.0, and (ii) PCR-based DEB2-specific dominant markers (FMDEB-274/275, FMDEB-282/283, FMDEB-284/285, FMDEB-462/463, FMDEB-476/477, FMDEB-504/505, FMDEB-517/518) (Table S1) developed once the DEB2 genome sequence was obtained, as described below. For genotyping the FLPs and the DEB2-specific dominant markers, PCR reactions were performed in 30 µl using 0.75 units of MyTaq DNA polymerase (Bioline, Meridian Life Science, Memphis, USA), 1x MyTaq reaction buffer (containing 5mM dNTPs and 15 mM MgCl2), 0.4 µM of each primer, and 50 ng of genomic DNA. The amplification was performed in a GeneAmp PCR system 9700 (Applied Biosystem, Foster City, CA, USA) with an initial denaturation step of 3 min at 95 °C followed by 34 cycles of 30 s at 95 °C, 30 s at annealing temperature (50-60 °C), and 1 min/kb at 72 °C, followed by an extension step of 7 min at 72 °C. PCR products were analysed by electrophoresis in a 1% to 1.5% agarose gel. The FLPs markers FMDEB-57/58 and FMDEB-59/60 showed duplicated positions in the genome assemblies, but only one locus was genotyped, the one where the distance between forward and reverse primers matched the amplicon length (the forward and reverse primers were separated by 900 bp at the locus actually genotyped, but they were separated by more than 7000 bp at the other locus). In addition, their PCR products were cloned and sequenced as described above to further confirm the locus being genotyped.

### Sequencing of new genomes

Different new genome assemblies developed in this study were used. They were obtained from the DEB2 genotype (assembly referred to as HannDeb2.20230803) and the accession PI 468691 of *H. debilis* subsp. *tardiflorus* (assembly referred to as HdebPI468691r0). It is worth noting that *H. debilis* PI 468691 is the same accession used in the interspecific crosses to develop the DEB2 line (Velasco et al., 2012). The *H. debilis* PI 468691 population used for sequencing is maintained at the LIPMe (Laboratoire des Interactions Plantes-Microrganismes; INRAe, Toulouse, France) and is resistant to the broomrape population GT (Chabaud et al., 2022). Additionally, the genome sequence of the susceptible parental sunflower landrace B117 was also obtained.

### a) *H. debilis* subsp. *tardiflorus* PI 468691 genome sequencing and annotation

*Helianthus debilis* accession PI 468691 was sequenced at the Gentyane INRAe platform (Clermont-Ferrand, France) using the PacBio Sequel II system. Canu v2.1 (Nurk et al. 2020) was used to assemble 6.92 million PacBio HiFi reads, representing 138.8 Gbp. Spurious short contigs were then removed by identifying hits with a larger contig that spanned 80% of the shorter contig’s length using Minimap2 (Li, 2018). This process resulted in a final genome assembly of 1,825 contigs, totalling 4.8 Gbp, with an N50 of 15.3 Mbp. Contigs were then scaffolded using Bionano Solve software (release 3.6.1; https://bionanogenomics.com/downloads/bionano-solve/) and an optical map spanning 4.6 Gbp with an N50 of 89 Mbp. Gene models were predicted using the EuGene pipeline version 1.6.6 (Sallet et al., 2019), a method previously described in Badouin et al. (2017). This gene model prediction was informed by 4.78 million PacBio IsoSeq transcript sequences (N50: 2,483 nucleotides; total: 10.6 Gbp) extracted from leaf, bud, and stem samples.

Protein-coding genes were annotated using a custom pipeline (Carrere and Gouzy, 2023) that integrated several sources of information. A tiered approach was used to prioritize results based on the expected accuracy of each source. Priority was given to the following sources in descending order: (1) Reciprocal Best BLASTp Hits: Genes were first annotated by identifying reciprocal best hits against a curated set of Asterids proteins from the Uniprot (downloaded on 2021-07-20) database that were tagged as "reviewed". A minimum sequence span of 90% and an identity of 80% were required for a hit to be considered; (2) Enzyme Commission (EC) Numbers: Putative enzymes were assigned EC numbers using the E2P2 tool (version 3.1). (3) Transcription Factors and Kinases: Transcription factors and kinases were identified using ITAK (release 1.7); (4) Transcription Factors: Additional transcription factors were identified using PlantTFCat; and (5) Functional Descriptions and Gene Ontology (GO) Terms: Finally, functional descriptions and GO terms were generated using the Blast2GO annotation service. The input for this service included data from InterPro (release 85.0) and BLASTp hits from the NCBI NR database (downloaded on 2021-07-20), both restricted to Viridiplantae proteins.

### b) DEB2 genome sequencing and annotation

The DEB2 genome was sequenced by CNRGV (Centre National de Ressources Génomiques Végétales) INRAe platform (Toulouse, France) using a PacBio sequencing platform. HiFiasm (Cheng et al., 2021) was used to assemble 4 million PacBio HiFi reads, representing 74.3 Gbp, thus, an estimated 21x genome coverage of corrected reads was achieved. 833 contigs, totalling 3.2 Gbp with an N50 of 35 Mbp were obtained. Assembly completeness was evaluated using BUSCO v4 (Seppey et al., 2019) with the viridiplantae_odb10 lineage dataset (downloaded on 2019-11-20). A high percentage of complete orthologs indicated strong completeness, particularly in genic regions. Optical mapping was performed using the Bionano Genomics Saphyr system. Sixty-nine optical maps were assembled for the DEB2 genotype. Hybrid scaffolding was conducted by combining the optical maps with the HiFiasm assembly. Finally, the genome of the DEB2 line was assembled at the chromosome level (17 chromosomes and 11 additional unanchored contigs), totalling 3.5 Gbp.

The EuGene pipeline version 2 (Sallet et al., 2019) was used for gene prediction, combining transcriptomic evidence and protein homology to identify gene models. Homology-based support was provided by multiple proteome references: *Brachypodium distachyon* (refProteome), *Arabidopsis thaliana* (TAIR10), and UniProt databases (Swiss-Prot and TrEMBL). Annotation completeness was evaluated with BUSCO using embryophyta_odb10 lineage dataset (created on 2020-09-10), obtaining a high percentage of expected orthologs (98.6% complete BUSCOs for the genome and 96.4% for the proteome). Transposable elements were identified and masked using Dfam consensus sequences (https://dfam.org) and the TransposonPSI database (http://transposonpsi.sourceforge.net). Within the EuGene pipeline, the handling of repeated regions was done in two steps: (i) Masking of all repeated elements identified with the software Red repeat masking and LTRharvest, and (ii) partial unmasking of specific regions in which Eugene identified locations where there are some hits of biological evidence, such as transcripts or protein or ncRNA, and unmask them before integrating all the results. These regions were also annotated, at the cost of unmasking small portions of adjacent repeated elements.

To obtain the functional annotation, protein sequences derived from the annotated coding sequence were first aligned to the *H. debilis* (HdebPI468691r0) and then to the reference sunflower (HanXRQv2) proteomes. Proteins showing more than 80% identity and coverage with a functionally annotated protein were assigned the corresponding functional annotation. The remaining proteins were further annotated by comparison against the UniProt/Swiss-Prot (SPROT) database downloaded on 2025-02-13. Genes whose encoded protein showed no significant similarity to any known proteins were classified as hypothetical proteins.

Finally, within the *Or_Deb2_* fine-mapped region, protein sequences were queried against the UniProt and the NCBI-ClusteredNR/Reference protein databases, and a domain structure analysis was performed using ScanProsite (de Castro et al., 2006) and InterproScan (Jones et al., 2014) to confirm putative functions. Additionally, since the Eugene pipeline used might annotate and unmask small portions of LTR (Long Terminal Repeat) elements, genes located within LTR retrotransposons were retained only if they did not contain domains associated typically with retroelements (gag, pol, reverse transcriptase, and integrase).

### c) B117 genome sequencing and annotation

The B117 genome was sequenced using a PacBio HiFi sequencing platform. A total of 3.55 million HiFi reads (62.97 Gbp) were assembled with HiFiasm (Cheng et al., 2021), producing 1,213 contigs totalling 3.1 Gbp and an N50 of 38.3 Mb. Assembly completeness was evaluated using BUSCO v4 (Seppey et al., 2019) with the embryophyta_odb10 lineage dataset (created on 2024-01-08). A high percentage (99%) of complete orthologs indicated strong completeness, particularly in genic regions. Contigs were scaffolded using RagTag (Alonge et al., 2022), resulting in a chromosome-level assembly comprising 17 chromosomes and 371 unanchored contigs of 0.5-1.5 Mbp, for a total genome size of 3.13 Gbp. Helixer (Stiehler et al., 2020) was used to identify the structural annotation of this genome. Retrotransposons were identified using LTRHarvest (Ellinghaus et al., 2008). Annotation completeness was evaluated with BUSCO using the embryophyta_odb10 lineage dataset (downloaded on 2024-01-08) and the annotated proteome contained 95% of complete BUSCOs. Functional annotation was performed using BLAST. First, predicted proteins were aligned against the HanXRQv2 protein set. Then, the corresponding coding sequences (CDS) were aligned against the HanXRQv2 CDS set. Proteins without annotation from these comparisons were subsequently searched against the UniProt/Swiss-Prot (SPROT) database downloaded on 2025-02-13. In all cases, only hits showing more than 80% identity and 80% coverage were retained. Genes whose encoded protein showed no significant similarity to any known proteins were classified as hypothetical proteins. Markers within the *Or_Deb2_*region were aligned to this genome. All of them mapped to Chr 4. *Or_Deb2_*flanking markers AX-105525205 and AX-105399507 were used to retrieve the corresponding *Or_Deb2_*-Chr 4 genomic interval in B117 (referred to as B117_*Or_Deb2_*_AX interval; supplementary File S1) for comparison with DEB2 genome assembly.

### Identification of the *H. debilis* region introgressed into DEB2

The DEB2 genotype was developed after crossing the sunflower (*H. annuus*) susceptible cultivar HA89 with *H. debilis* subsp. *tardiflorus* accession PI 468691 and two rounds of backcrossing to HA89 (Velasco et al., 2012). In order to further characterize the *Or_Deb2_* region, an analysis to determine the *H. debilis* genomic region introgressed into DEB2 and associated to *Or_Deb2_*was carried out. For that, markers used for the *Or_Deb2_* fine mapping were aligned to the *H. debilis* assembly (HdebPI468691r0) by BLAST, all of them being positioned in one scaffold (HdebPI468691s0002, 213-Mbp long). Then, sequences of Chr 4 from the HA89 genome assembly HanHA89r1 (Huang et al., 2023) and the scaffold HdebPI468691s0002 from the *H. debilis* genome assembly were aligned to Chr 4 of the DEB2 genome assembly (HannDeb2.20230803) using D-genies (Cabenettes and Klopp, 2018).

### Comparative analysis of the *Or_Deb2_* locus in other genomes

The region containing the larger *Or_Deb2_* locus defined by Fernández-Aparicio et al. (2022) and delimited by markers AX-105525205 and AX-105399507 was extracted in all the sunflower genomes available to date, obtained from the sunflower (*H. annuus*) cultivated genotypes XRQ, PSC8, RHA438, IR, HA89, LR1, OQP8, HA300 (Badouin et al., 2017; Huang et al., 2023), and B117 (this study; B117_*Or_Deb2_*_AX interval), and the wild relatives *H. annuus* PI659440 (Huang et al., 2023), *H. anomalus* PI 468642 (Fernández-Melero et al., 2024) and *H. debilis* PI 468691 (this study), and compared to that in DEB2. Conservation of the order of the markers and protein-coding genes among the different genomes was studied. Synteny plots were generated using NGenomeSyn v1.41 (He et al., 2023). Pairwise BLASTp searches were performed between predicted protein sequences, and only hits showing ≥90% identity were considered. In duplicated regions where multiple proteins from one genome matched multiple proteins from other genome, only the best-ordered one-to-one correspondence was retained to preserve collinearity and accurately represent synteny between genomes. Protein multiple sequence alignments were performed using Clustal Omega in MegAlign Pro software within the DNASTAR, Inc. package with default parameters.

### cDNA *Or_Deb2_* candidate gene cloning and expression analyses

Selected candidate genes for *Or_Deb2_*, as explained in the results section, were cloned and sequenced from cDNA from the resistant parental genotype DEB2 and the susceptible B117. For this, two pools of root cDNA from three of DEB2 and B117 replicates inoculated with seeds of the GT broomrape population were prepared. Mini-rhizotron assays to obtain samples of sunflower roots parasitized by *O. cumana* and RNA extraction for cDNA synthesis were carried out as described in the following section. PCR primers used to get the cDNA sequences from selected candidate genes are detailed in Table S2. cDNA pools were amplified using 30 μl of reaction mixture containing 1x buffer, 1mM of dNTPs, 3mM of MgCl2, 0.4 µM of each primer, 0.5 U of Taq polymerase (Velocity ™ Bioline, Meridian Life Science, Memphis, USA), and 30 ng of cDNA. The amplification was performed in a GeneAmp PCR System 9700 (Applied Biosystems, Foster City, CA, USA) with an initial denaturation step at 98 °C for 5 min, followed by 34 cycles of 30 s at 98 °C, 45 s at the annealing temperature for each primer, and 2.5 min at 72 °C, and ending with an extension period of 7 min at 72 °C. PCR products were separated on a 1.5% agarose gel and purified using the MEGAquick-spin TM Plus Total Fragment DNA Purification Kit (iNtRON Biotechnology, Inc., Korea) for cloning. The purified fragments were ligated into the pSPARK Done DNA cloning vector system (Canvax Biotech, Córdoba, Spain) as described by the manufacturer and Sanger sequenced (StabVida, Lisbon, Portugal).

All the obtained cDNA sequences were aligned through BLAST to the DEB2 and B117 genome assemblies to confirm the different loci cloned, and compared to orthologous genes in all the sunflower genomes available to date. Sequence analysis was conducted using the Lasergene SeqMan Ultra and MegAlign Pro software within the DNASTAR, Inc. package. Domain structure analysis was performed as indicated above.

The *Or_Deb2_* candidate gene expression analysis was conducted with a chip-based digital PCR (dPCR) system (QuantStudioTM Absolute QTM Digital PCR, ThermoFisher Scientific). The assay was carried out with two technical replicates conducted on different days using cDNA from: (i) three biological replicates of the lines DEB2 and B117 infected with broomrape race GT and sampled at 14- and 21-days post inoculation (dpi), and (ii) three biological replicates of non-infected plants of the same both genotypes at both 14- and 21-dpi stages. Mini-rhizotron assays to obtain the cDNA samples are described in the next section. Primers and probes for the dPCR (Table S2) were designed based on the DEB2 sequence assembly using the IDT RealTime qPCR Design Tool (https://eu.idtdna.com/scitools/Applications/RealTimePCR/). Their specificity was verified by comparison with the B117 genome assembly. Sequences were compared and checked for primer dimer formation using the Multiple Primer Analyzer available on the ThermoFisher webpage. Two sets of primers and probes were designed: (i) HANNDEB2CHR04g005720-exp, designed to amplify only the *Or_Deb2_* best candidate *HannDeb2Chr04g005720* based on a deletion found in this gene, and (ii) ACT-exp, developed to amplify the actin gene, used as reference (GenBank Accession number FJ487620) (Ochogavía et al., 2017). To avoid contaminant gDNA amplification, each set of primers and probes was designed to amplify the junction between two different exons. Each 10 µl of reaction mixture consisted of 900 nM of the forward and reverse primers, 250 nM of the probes, 1x of the Master Mix provided by the manufacturer, and 50 ng of cDNA. Molecular-grade water was used for non-template control reactions. According to the manufactureŕs instructions, 9 µl of reaction mixture was loaded into each well, followed by 15 µl of isolation buffer. The PCR protocol consisted of 10 min at 96 °C, followed by 40 cycles of 5 s at 96 °C and 15 s at 58 °C. After checking the viability of each set of primers and probes separately, all the primers and probes were multiplexed. The absolute quantity of cDNA per sample (copies/µl) was determined using the software provided by the manufacturer. The threshold value was manually set for each probe by discriminating between positive and negative droplets in all the samples in each experiment. Actin was used as a reference gene for data normalization. Data were analysed through ANOVA with sunflower genotype (B117 and DEB2), inoculation/non-inoculation, and time of sampling (14-dpi and 21-dpi) as fixed factors. Significance was established at *p* < 0.05.

### Mini-rhizotron assays and cDNA preparation for cloning and expression analyses

Mini-rhizotron assays to obtain sunflower root tissue for cDNA synthesis were performed as described by Le Ru et al. (2021). Three independent mini-rhizotron assays were assembled with 10 DEB2 plants inoculated with seeds of GT broomrape population, 5 B117 plants inoculated with seeds of GT, 5 DEB2 plants non-inoculated, and 5 B117 plants non-inoculated. Samples were taken at 14- and 21-dpi as follows. Two cm sections of sunflower roots, with a broomrape attachment in the middle in the case of inoculated plants, were cut, placed directly in liquid nitrogen, and frozen at -80 °C. For each of the three assays, 140 sections of DEB2 and B117 inoculated and non-inoculated were taken at 14-dpi, and 60 sections of each genotype at 21-dpi. Mini-rhizotrons used to take samples at 14-dpi were discarded after the sampling to avoid gene expression bias in samples at 21-dpi. RNA extraction was performed using the kit RNeasy Plant Mini Kit (Qiagen, Valencia, CA, USA). RNA concentration was quantified in a NanoDrop™ Lite Spectrophotometer (Thermo Fisher Scientific, Wilmington, DE, USA), and RNA integrity was quantified with the FemtoPulse system (Agilent 2100 Bioanalyzer, Agilent Technologies Inc., Santa Clara, CA, USA). 1 µg of RNA was then retrotranscribed into cDNA using the protocol of Transcriptor First Strand cDNA Synthesis Kit (Roche, Switzerland).

## Results

### Fine mapping *Or_Deb2_*

#### a) Initial rounds for *Or_Deb2_* fine mapping: F_2_ phenotyping and genotyping with Iasnip-and Axiom- SNP markers

DEB2 exhibited resistance to broomrape infection while B117 was susceptible, which was in concordance with previous studies (Martín-Sanz et al., 2016; Fernández-Aparicio et al., 2022). HA821, XRQ, and PSC8 were also susceptible, showing an average number of broomrape stalks per plant from 8.5 to 61. In the F_2_ fine-mapping population, there were 2921 resistant plants and 748 susceptible plants (27 plants died and could not be evaluated), corroborating the dominance of the gene responsible for the resistance. The observed segregation ratio (2921:748) was significantly different from the expected segregation ratio 3:1 based on the χ2 test (*p* < 0.001). There was a deficiency in the number of susceptible F_2_ plants, and a corresponding excess of resistant F_2_ plants. Considering genotyping data obtained later, this segregation distortion was caused by some susceptible F_2_ plants (a total of 169) that escaped broomrape infection. Escapes have been observed in broomrape evaluations due to the influence of environmental conditions in the phenotypic assays (Fernández-Martínez et al., 2015). In genetic studies, escapes in the F_2_ generation are commonly detected by evaluating the F_3_ generation of the complete F_2_ population, which is impractical when the population size is large, as in the present study. They were instead detected using molecular markers tightly linked to the *Or_Deb2_* gene. Accordingly, the 169 F_2_ genotypes identified as escapes were excluded from the recombinant analysis.

In order to fine-map *Or_Deb2_*, the F_2_ fine-mapping population was genotyped with a set of newly developed SNPs (prefix Iasnip-) designed along annotated genes located in the 1.38-Mbp interval between AX-105525205 to AX-105399507 in the HanXRQr2.0 assembly. Some Axiom markers (prefix AX-) formerly used to map *Or_Deb2_* were also included, although the flanking markers AX-105525205 and AX-105399507 were not polymorphic in this population. Genotyping the fine mapping F_2_ population yielded, on the left side of the *Or_Deb2_* target interval, 65 recombinants between Iasnip-41 and FMDEB-57/58_59/60 (Table 1). The nomenclature FMDEB-57/58_59/60 denotes two markers, FMDEB-57/58 and FMDEB-59/60, derived from the same gene (Table S1) and co-segregating, which is also used in the manuscript for other co-segregating marker pairs. Subsequently, 1 recombinant (R1) was identified between FMDEB57/58_59/60 and Iasnip-30 (Table 1). On the right side, 47 recombinants were found between Iasnip-49 and Iasnip-15_16 (Table 1). Then, five recombinants were found between Iasnip-50_55 and Iasnip-49 (Table 1). Finally, a more precise delineation of the right side was achieved by identifying four recombinants (named as R2 to R5) between markers Iasnip-30 and Iasnip-50_55. All these data narrowed the *Or_Deb2_* target interval between FMDEB57/58_59/60 and Iasnip-50_55. Blast of these markers into the HanXRQr2.0 assembly showed that markers FMDEB-57/58_59/60 were located at duplicated positions at 8.25 and 8.28 Mbp, the second inside a gene annotated as heat shock protein HSP20 (*HanXRQr2Chr04g0142331*). Iasnip-50 and Iasnip-55 were located at duplicated positions 8.31 and 8.99 Mbp inside the acyl-CoA desaturase loci *HanXRQr2Chr04g0142341* and *HanXRQr2Chr04g0142551*. In the HanPSC8 assembly, unique positions for all markers were identified (Table 1), delineating a 151-kbp a window for the *Or_Deb2_* locus.

**Table 1.**
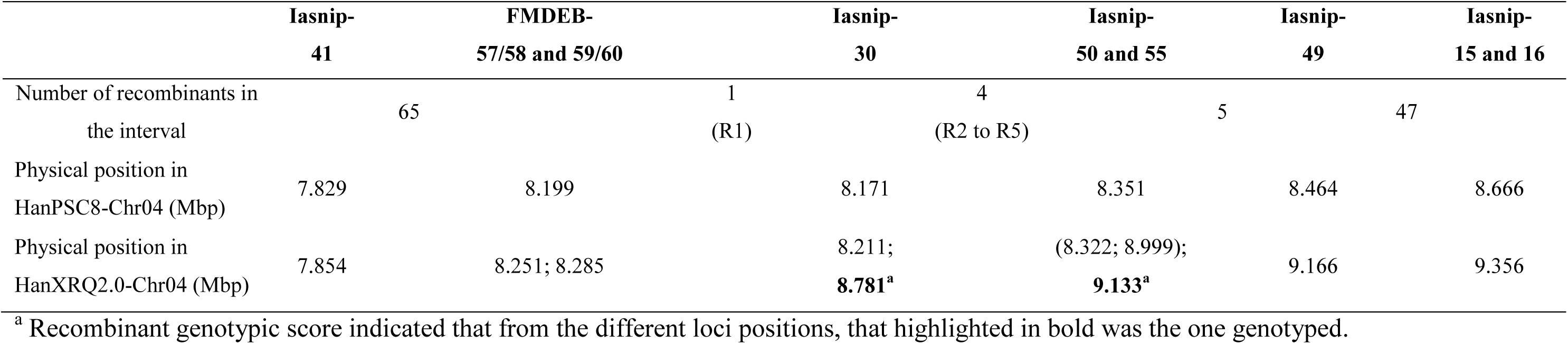
Number of F_2_ plants showing a recombination event inside the AX-105525205 to AX-105399507 1.38-Mbp *Or_Deb2_* target interval and physical position of the markers based on the two sunflower reference genomes assemblies HanPSC8 and HanXRQr2.0.

### b) Saturation of the *Or_Deb2_* region with additional DEB2-specific markers based on the DEB2 genome assembly

In HannDeb2.20230803, FMDEB-59/60 was located at 8,660,219 bp, while Iasnip-50 was at 8,997,981 bp, delineating a window of 338 kbp. Iasnip-30, located inside this interval, was positioned at 8,756,009, 8,838,222, and 8,913,233 bp, which corresponded to three of four repeated regions, as will be explained below. Further refinement of the FMDEB-57/58_59/60 to the Iasnip-50_55 region was achieved using new PCR-based markers based on the DEB2 genome sequence. Seven primer pairs present in DEB2-HannDeb2.20230803 and absent in B117 were designed within the 338-kbp interval. These markers, which show a dominant polymorphism between the parental lines DEB2 and B117, were used to genotype the recombinant plants found between FMDEB-57/58_59/60 and Iasnip-50_55 (R1 to R5) (Table 1, Figure 1A, Table S3). No polymorphic markers were found between FMDEB-57/58_59/60 and Iasnip-30, so it was not possible to further refine the left side of the *Or_Deb2_* interval, which was delimited by the recombinant R1.

**Figure 1.**
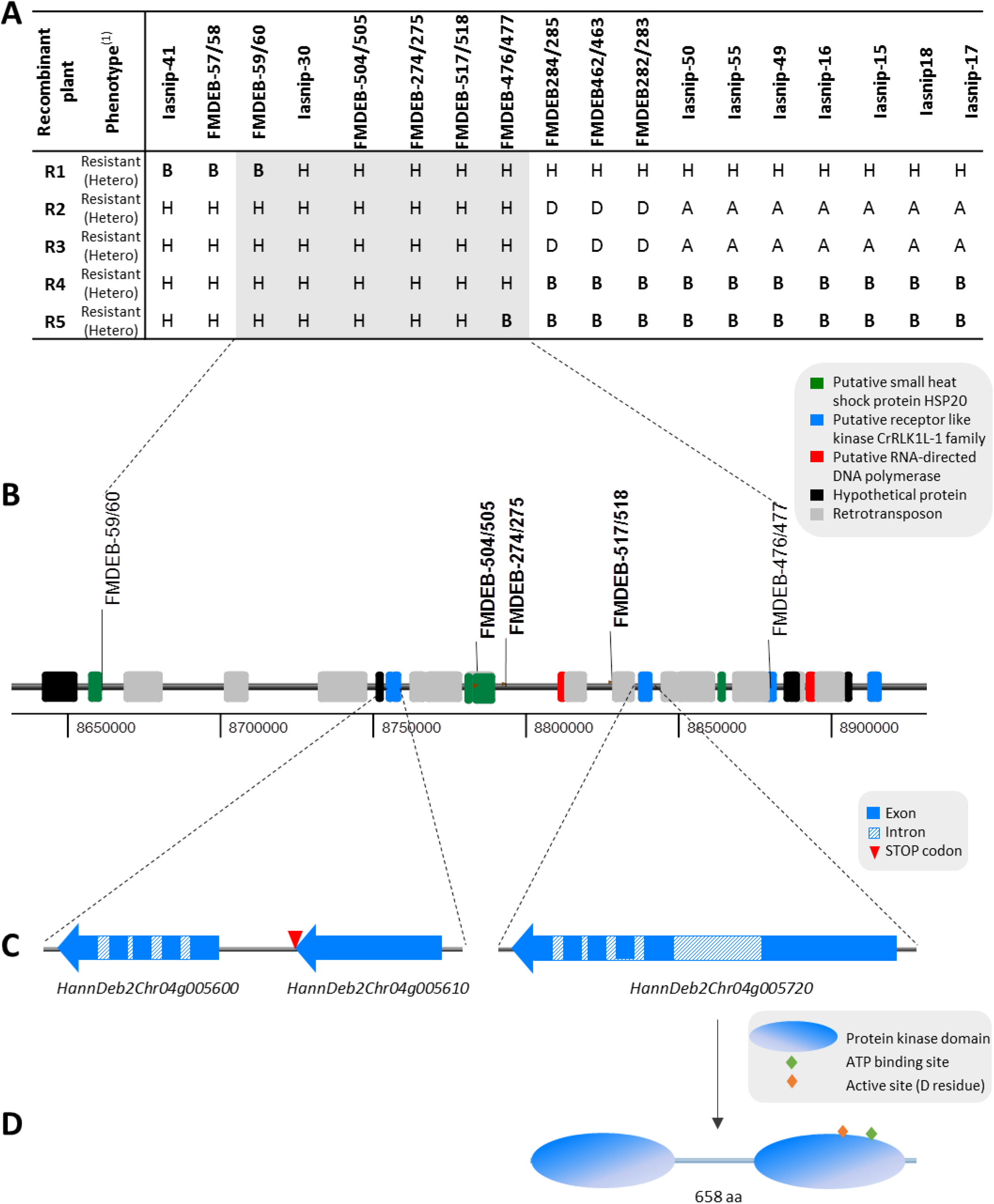
Fine mapping and gene structure at the *Or_Deb2_* resistance locus. **A)** Genotype and phenotype (for race G broomrape resistance) for the five FMDEB-57/58_59/60 to Iasnip-50/55 recombinant F_2_ plants (R1 to R5). The Iasnip- prefix indicates SNP markers and the FMDEB- prefix indicates PCR-based markers. A= DEB2 (resistant) allele; B= B117 (susceptible) allele, H= Hetero, D= A or H. F_2_ scores for DEB2 dominant markers were based on the genotyping of the corresponding F_3_ families, as detailed in Table S3, in such a way that the F_2_ was scored as B when the F_3_ was fully homozygous B (no amplification at any individual); H when F_3_ segregated and D when the whole F_3_ was A or H (all individuals showed amplification so no distinction can be made between homozygous A or Hetero). *Or_Deb2_* fine-mapped region between markers FMDEB-59/60 and FMDEB-476/477 is marked in grey; **B)** Detailed annotated structure of the *Or_Deb2_* fine-mapped region; **C)** Detailed structure of the RLKs found in the *Or_Deb2_* fine-mapped region; **D)** Protein configuration of the *HannDeb2Chr04g005720* gene based on cDNA cloning. ^(1)^The phenotype column shows the F_2_ plant phenotype and in brackets the phenotype based on the corresponding F_3_ segregation – Due to the dominant nature of *Or_Deb2_* resistance, F_2_ plants R1 to R5 were resistant and based on phenotypic segregation of their corresponding F_3_ families for both resistant and susceptible plants (Table S3), they were classified as heterozygous.

DEB2 dominant markers had the disadvantage of not allowing distinction between heterozygous and homozygous genotypes. Thus, only the right-side recombinant plants R4 and R5, which were resistant but showed a susceptible genotype on the right of the *Or_Deb2_* locus, were informative for fine mapping the *Or_Deb2_* gene. The right side was first refined by the recombinant R4, which showed a recombination event between FMDEB-476/477 and FMDEB-462/463, and then by the recombinant R5, which showed a recombination event between FMDEB-517/518 and FMDEB-476/477 (Figure 1A, Table S3). This data allowed us to fine-map the *Or_Deb2_* gene to a 219.4-kbp interval between FMDEB-57/58_59/60 and FMDEB-476/477 on Chr 4 of the DEB2 genome assembly. Markers Iasnip-30, FMDEB-504/505, FMDEB-274/275 and FMDEB-517/518 were identified as putatively co-segregating with the gene. The R1 to R5 corresponding F_2:3_ families (F_3_ plants) were also genotyped with this set of markers (Table S3). Their genotyping enabled us to distinguish between heterozygous (H) and DEB2 homozygous (A) genotypes in dominant F_2_ scores (D = A or H) (Table S3) and corroborated the *OrDeb2* fine-mapped interval.

### Analysis and dissection of the *Or_Deb2_* locus

#### a) Study of the *H. debilis* fragment introgressed into DEB2

The *H. debilis* introgression into DEB2 was analysed through the comparison among the genome assemblies of: (i) HA89, the *H. annuus* cultivar used to develop DEB2 (genome assembly HanHA89r1.0; Huang et al., 2023), (ii) DEB2 (HannDeb2.20230803; this study) and (iii) *H. debilis* subsp. *tardiflorus* (HdebPI468691r0; this study). The analysis showed that a 17-Mbp segment on the upper part of Chr 4 from the DEB2 line was introgressed from *H. debilis*, with the rest of the chromosome identified as coming from HA89 (Figure S2).

### b) Analysis of the AX-105525205 to AX-105399507 larger *Or_Deb2_* region in DEB2 and *H. debilis* PI468691 newly developed genome assemblies

Markers AX-105525205 and AX-105399507 used by Fernández-Aparicio et al. (2022) to map *Or_Deb2_* were aligned to the DEB2 HannDeb2.20230803 assembly. They delineated a window in Chr 4 of 1.5 Mbp, from positions 7,811,184 to 9,345,095, which was slightly bigger than the 1.38 Mbp in the HanXRQr2.0 assembly. Marker order was overall conserved in both assemblies, but genomic sequences were highly divergent (Figure S3A). The analysis of the DEB2 1.5 Mbp region revealed four duplications spanning between 8.74 and 8.93 Mbp (marked with a square in Figure S3B) with more than 99% of identity among them, and a total of 42 annotated protein-coding genes (Table S4A).

Using the same approach, the genomic region corresponding to the larger *Or_Deb2_* locus of Fernández-Aparicio et al. (2022) was analysed in the *H. debilis* assembly (HdebPI468691r0). Flanking markers AX-105525205 and AX-105399507 mapped onto the *H. debilis* genome delimited a 1.39-Mbp region on scaffold s0002 between positions 7,360,261 and 8,754,638. A total of 31 protein-coding genes were annotated in this interval (Table S4B). The four duplications identified in this region in DEB2 were not observed in the *H. debilis* genome.

### c) Analysis of the AX-105525205 to AX-105399507 larger *Or_Deb2_*region in ten other sunflower genome assemblies previously published

The region delimited by markers AX-105525205 and AX-105399507 was extracted from the genome assemblies of the *H. annuus* cultivated lines XRQ, PSC8, RHA438, IR, HA89, LR1, OQP8, HA300, and the wild relatives *H. annuus* PI659440 and *H. anomalus* PI 468642, and compared to that in DEB2, *H. debilis* PI468691, and B117. The structure and annotation of this region in each different genome were analysed and are shown in Figures 2 and 3. The order of the markers, as well as the gene order, was generally conserved across all assemblies. Three of the cultivated sunflower genomes, HA89, RHA438, and LR1, as well as that of the parental B117, shared the same structure as XRQ with two duplicated subregions, according to gene order and amino acid sequence identity of the gene products, as explained by Fernández-Aparicio et al. (2022), and they will be referred to as XRQ-type. Contrarily, the cultivated sunflower accessions HA300, OQP8, PSC8, and IR, and the wild species *H. debilis* PI 468691, and *H. anomalus* PI 468642 showed no such duplicated subregions in this larger interval, and they will be referred to as PSC8-type (Figure 2 and Figure 3). The wild *H. annuus* PI659440 exhibited a PSC8-type configuration, but in this case, it was duplicated in another part of the genome, 8-Mbp away. This configuration differed from the two-subregion arrangement, as they were not contiguous, and the flanking markers were detected twice in the *H. annuus* PI659440 genome.

**Figure 2.**
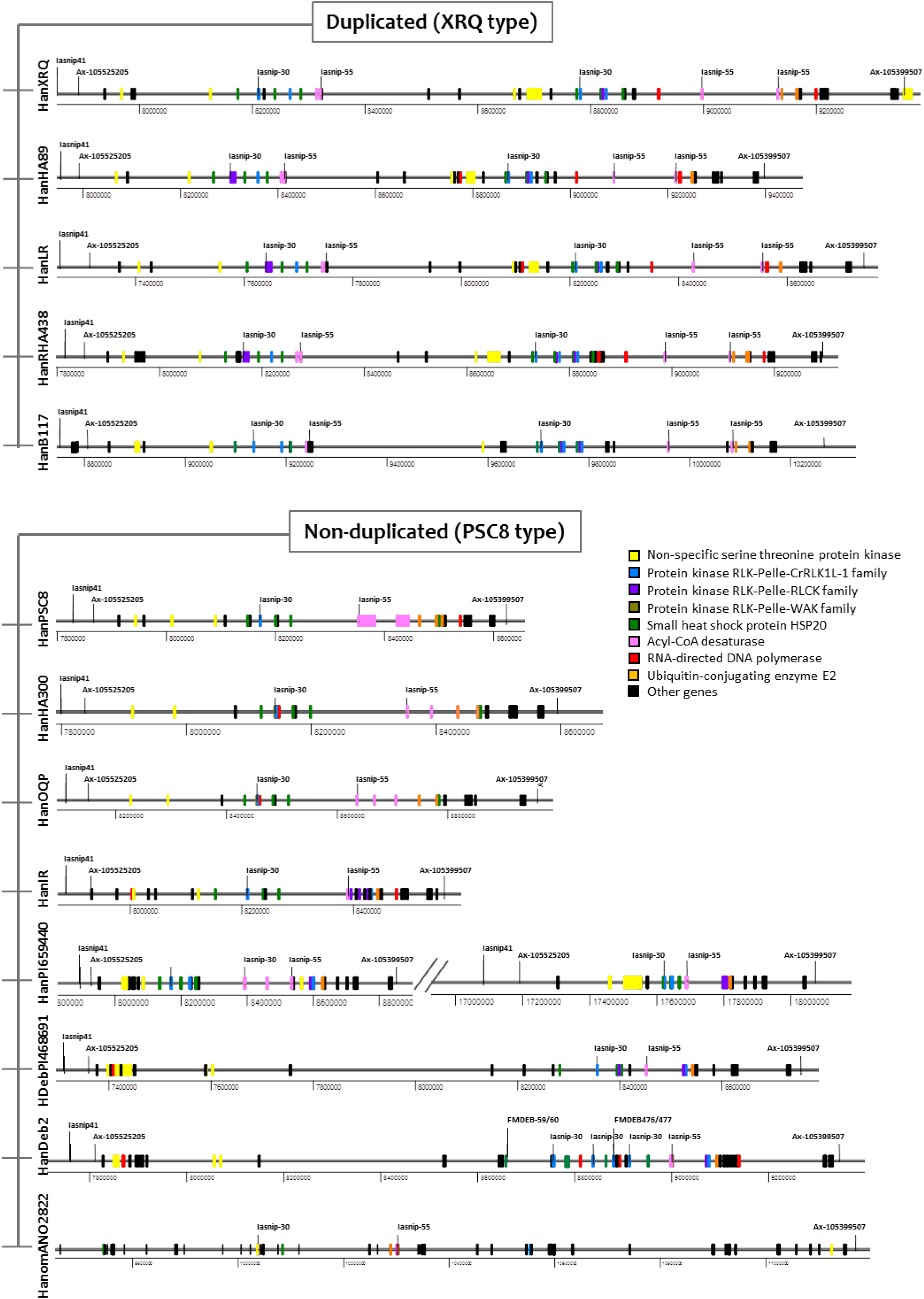
Gene structure and nature in the region delimited by AX-markers 105525205 and AX-105399507 containing the larger *Or_Deb2_* locus defined by Fernández-Aparicio et al. (2022) within all the sunflower genomes available to date, obtained from the cultivated lines XRQ (assembly HanXRQr2.0), PSC8 (HanPSC8r1.0), RHA438 (HanRHA438r1.0), IR (HanIRr1.0), HA89 (HanHA89r1.0), LR1 (HanLR1r0.9), OQP8 (HanOQP8r0.9), HA300 (HanHA300r0.9) (Badouin et al., 2017; Huang et al., 2023), and B117 (HanB117; this study), and the wild relatives *H. annuus* PI659440 (HanPI659440r1.0; Huang et al., 2023), *H. anomalus* PI 468642 (HanomANO2822; Fernández-Melero et al., 2024) and *H.debilis* PI 468691 (HdebPI468691r0; this study), and compared to that in DEB2 (HannDeb2.20230803).

**Figure 3.**
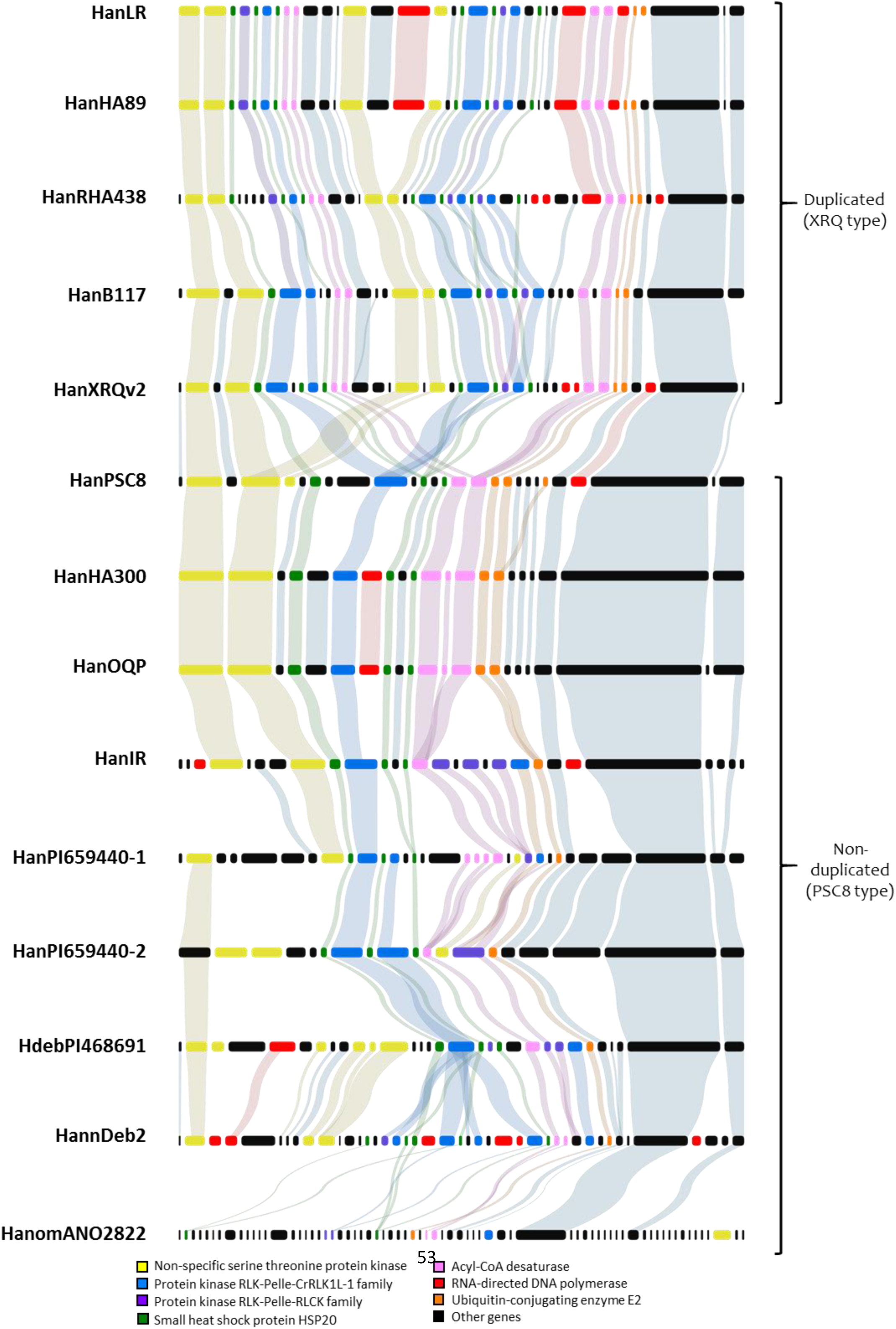
Synteny of the genomic region delimited by markers AX-105525205 and AX-105399507, containing the *Or_Deb2_* locus defined by Fernández-Aparicio et al. (2022), across all currently available sunflower genomes. The analysis includes cultivated lines XRQ (assembly HanXRQr2.0), PSC8 (HanPSC8r1.0), RHA438 (HanRHA438r1.0), IR (HanIRr1.0), HA89 (HanHA89r1.0), LR1 (HanLR1r0.9), OQP8 (HanOQP8r0.9), HA300 (HanHA300r0.9) (Badouin et al., 2017; Huang et al., 2023), and B117 (HanB117, this study), as well as wild relatives *H. annuus* PI659440 (HanPI659440r1.0; Huang et al., 2023), *H. anomalus* PI468642 (HanomANO2822; Fernández-Melero et al., 2024), and *H. debilis* PI468691 (HdebPI468691r0; this study), and is compared to the corresponding region in DEB2 (HannDeb2.20230803). Protein-coding genes are represented as colored boxes, with conserved orthologs connected by lines. The figure highlights conserved genomic organization as well as structural variations among sunflower assemblies.

The DEB2 configuration also corresponded to the PSC8 type; however, it displayed a specific feature: although the region was not duplicated as a whole, four local mini-duplications, spanning approximately 10 kbp, were detected, as described above (Figure S3B). DEB2 proteins sharing >90% identity with their orthologues in other genomes were identified, such as small heat shock proteins HSP20, acyl-CoA desaturases, RNA directed DNA polymerases, ubiquitin conjugating enzymes E2, transcription factors of the CCHC(Zn) family, RLPs (with a large LRR ectodomain, a single transmembrane helix and no kinase domains) and RLKs (with one or two fused kinase domains) (Figure 3).

### d) Analysis of the *Or_Deb2_* fine-mapped region in DEB2 and *H. debilis* PI468691 genome assemblies

In DEB2, a total of eight protein-coding genes were annotated within the 219.4-kbp *Or_Deb2_* fine-mapped region delimited by the recombinant analyses (Fig. 1B and Table S4A). Among them, one gene (*HannDeb2Chr04g005590*) was annotated as a hypothetical protein, a prediction with no supporting BLAST similarities, so it was considered an annotation artefact. Three other genes (*HannDeb2Chr04g005600*, *HannDeb2Chr04g005610*, and *HannDeb2Chr04g005720*) were annotated as putative RLKs (RLK-Pelle), whose detailed analysis is provided in the following sections. Another gene (*HannDeb2Chr04g005700*) was identified as a putative RNA-directed DNA polymerase. Because such genes are typically derived from transposable elements, it was likely to represent an unmasked domain of the adjacent retrotransposon (shown in Figure 1B). Finally, three genes encoding putative small heat shock proteins (HSP20; *HannDeb2Chr04g005670*, *HannDeb2Chr04g005690*, and *HannDeb2Chr04g005770*) were also annotated, with *HannDeb2Chr04g005670* and *HannDeb2Chr04g005770* being identical. *HannDeb2Chr04g005670*/g005770 and *HannDeb2Chr04g005690* encoded 118-aa and 214-aa proteins, respectively, that belonged to the protein family “Heat shock protein 21-like” (IPR044587) and presented a small heat shock protein (sHSP) domain profile (PS01031). Robust lines of evidence that will be discussed in the Discussion section supported the RLKs identified in the 219.4-kbp fine-mapped region as the most plausible candidates underlying *Or_Deb2_*, and, accordingly, they were analysed in detail.

It is worth mentioning that the RLKs in the DEB2 219.4-kbp fine-mapped interval were located at the first two of the four repeated genomic intervals found in this region (Figure S3B). Notably, other RLKs were found in the other two repeated regions, starting at position 8.8 Mbp and at 8.9 Mbp (Table S4A). Figure 4 represents the structure and homology analysis of the four repeated genomic regions containing these RLKs, which, for simplicity, will be hereafter referred to as *RLK-1* to *RLK-4* (located, respectively, at the first through fourth genomic repeats, with *RLK-1* and *RLK-2* inside the 219.4-kbp fine-mapped region; Figure 4, Table S4A). *RLK-2* (*HannDeb2Chr04g005720*) and *RLK-4* (*HannDeb2Chr04g005880*) corresponded to full-length kinases, whose domain analysis and classification are provided in the next section. In contrast, *RLK-1* in the first duplication contained two adjacent RLK annotated genes (*HannDeb2Chr04g005600* and *HannDeb2Chr04g005610*) that, after being aligned with the full-length kinases, were revealed as being a truncated version of the full-length RLKs showing a premature stop codon (Figure 4). A similar situation was observed in the third duplication, where the full-length kinase was truncated due to the insertion of transposable elements. In this case, *HannDeb2Chr04g005800* and *HannDeb2Chr04g005820* together represent the truncated RLK *RLK-3*

**Figure 4.**
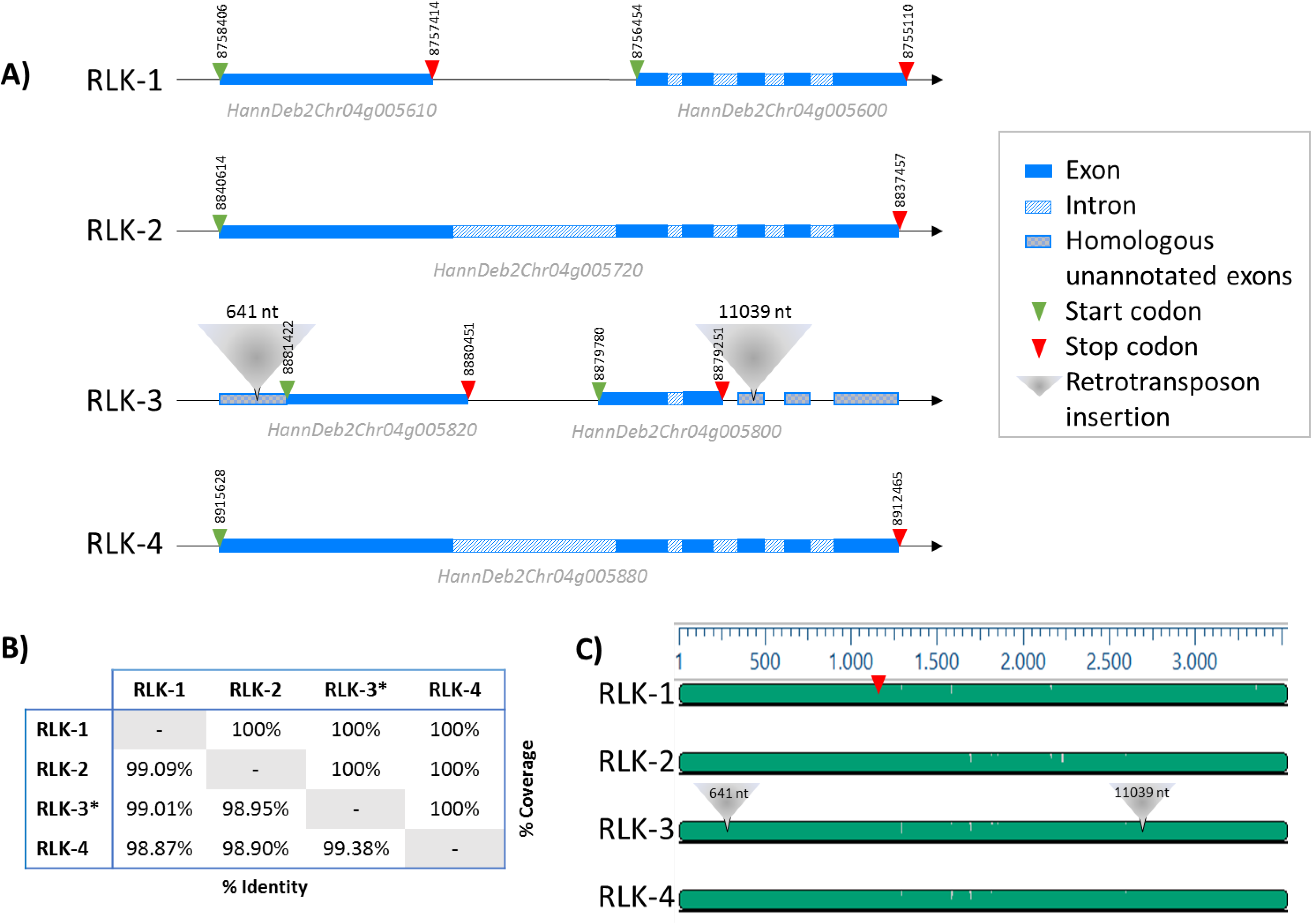
Genomic comparison of RLK gene regions across four repeated loci. **A)** Genomic organization of four regions containing receptor-like kinase (RLK) annotations. Although multiple RLK genes are annotated in some loci, sequence alignment shows they represent fragments or truncated versions of the same gene. Full-length kinase genes are found in repeats 2 (*RLK-2*) and 4 (*RLK-4*), whereas repeats 1 (*RLK-*1) and 3 (*RLK-3*) contain incomplete versions due to a premature stop codon and retrotransposon insertions, respectively. The position of the start and stop codons are indicated. **B)** Pairwise nucleotide identity and coverage percentages between the genomic regions (*retrotransposons removed), supporting the high sequence similarity among them. **C)** Nucleotide alignment of the four regions, showing conserved sequences and structural disruptions consistent with the observations in (A).

In *H. debilis*, the *Or_Deb2_*-corresponding fine-mapped region was studied by mapping the markers defining this region in the HdebPI468691r0 assembly and analysing annotated genes. Marker FMDEB-57/58 was located at position 8,284,807 on HdebPI468691r0 scaffold s0002, whereas marker FMDEB-476/477 was not found (Table S1). However, markers located downstream of FMDEB-476/477 were found from position 8,400,535 onward; therefore, this position was considered as the fine-mapping interval right border. FMDEB-517/518 and Iasnip-30, which co-segregated with the resistant locus, were located at positions 8,344,097 and 8,355,055, respectively. Consequently, the *Or_Deb2_* fine-mapped region corresponded to a 116-kbp interval in the *H. debilis* PI468691 genome. Three protein-coding genes were annotated within this region: a putative protein kinase RLK-Pelle-CrRLK1L-1 (*Hdeb2414_s0002g00045311*), a putative small heat shock protein HSP20 (*Hdeb2414_s0002g00045321*), and a putative protein kinase RLK-Pelle-RLCK-VIIa2 (*Hdeb2414_s0002g00045341*) (Table S4B). A detailed synteny analysis of the proteins within the *Or_Deb2_* fine-mapped region between DEB2 and *H. debilis* PI468691 is presented in Figure S4, revealing Hdeb2414_s0002g00045311 and Hdeb2414_s0002g00045341 as orthologues, respectively, to DEB2 full-length RLK-2, and to truncated DEB2 RLKs.

### Detailed analysis of the DEB2 *Or_Deb2_* candidates: cDNA cloning, sequence analysis, expression, and comparison with their orthologues

#### a) cDNA cloning and detailed sequence analysis

The RLKs located within the 219.4-kbp interval (*RLK-1* and *RLK-2*) were further analysed. The DEB2 genomic sequences of these RLKs extracted from the DEB2 genome assembly were 3526-bp for *RLK-1*, and 3520-bp for *RLK-2*, respectively, and identity among them was ≥ 99%. There were a number of SNPs among these genes, and a premature stop codon was present in *RLK-1*, as mentioned above.

The coding sequence of *RLK-2* was obtained from DEB2 cDNA extracted from infected plants at 14- and 21-dpi. This gene was expressed in both developmental stages, and a cDNA sequence was obtained from 7 out of 18 total cloned sequences. The cloned cDNA was identical to the coding sequence predicted from its corresponding gDNA annotation. The *RLK-2* gene contained 6 exons (Figure S5) and encoded a 658-aa protein (Figure S6). Domain structure analysis revealed a protein with two tandem fused kinase domains (Figure 1D, Figure S6A). The first domain was a kinase domain that carried an essential conserved aspartate residue as a proton acceptor in the active site and also conserved residues at an activation loop (A-loop), an ATP binding site, and a polypeptide substrate binding site, all within the catalytic domain of serine/threonine kinases (InterPro cd14066). The second one was a pseudokinase domain that lacked the above-mentioned conserved residues (Figure S6B). This structure was consistent with the kinase-pseudokinase protein structure, which belongs to the tandem kinase proteins (TKPs) family (Klymiuk et al., 2021) within the kinase fusion proteins (KFPs) class (Fahima and Coaker, 2023). RLK-2 exhibited the same structure kinase-pseudokinase as some TKPs that have been experimentally validated as resistance genes in the literature (*WTK1*, *WTK3*, *RWT4*, *WTK5*, *WTK6-vWA* and *WTK6b-vWA*; Figure S7), all of them from monocots in the *Triticeae* tribe (Reveguk et al., 2025). Amino acid sequence similarities found between RLK-2 and the other sequences ranged from 53% for WTK1 (32% identity) to 48% for WTK6b-vWA (29% identity). In all cases, the active site of the kinase domain was a residue of aspartate. Based on the classification by Lehti-Shiu and Shiu (2012), the kinase domain from RLK-2 was classified as RLK-Pelle CrRLKL1L-1, while its pseudokinase domain was classified as RLK-Pelle RLCK-VIIa-2.

Additionally, due to the high sequence similarity, the DEB2 cDNA of the full-length *RLK-4*, outside the fine-mapped interval, was also obtained. The gene was also expressed in the infected roots at both 14- and 21-dpi and 11 sequences out of 18 total cloned cDNAs corresponded to this gene. The cloned cDNA sequence was identical to that predicted from its corresponding gene annotation and encoded a 662-aa protein that shared the same tandem kinase structure as RLK-2, but contained 11 misassembled residues (Figure S5). None of the cloned DEB2 cDNAs corresponded to *RLK-1*. The *RLK-1* gene encodes a truncated protein due to a premature stop codon within the conserved kinase domain (Figure 4). As a result, the two kinase and pseudokinase domains were annotated as the independent loci *HannDeb2Chr04g005600* and *HannDeb2Chr04g005610* (Figure 4), which encoded, respectively, a predicted protein of 282-aa with the pseudokinase domain and another of 330-aa with the kinase domain.

The genomic structure in the *Or_Deb2_* interval of the susceptible parental line B117, (B117_*Or_Deb2_*_AX region) revealed two orthologous full-length RLKs (*B117Chr04_g003667* and *B117Chr04_g003660*). Their cDNA sequences were obtained at infected roots at 21-dpi (one clone for each one) and matched the corresponding predicted coding nucleotide sequence at 100% (Figure S5). Comparison of genomic and cDNA sequences identified a six-exon gene (Figure S5) encoding 662-aa proteins with a kinase-pseudokinase domain architecture similar to that of RLK-2 (Figure S6C).

### b) Expression analyses

#### b1) Expression of DEB2 HannDeb2Chr04g005720 (RLK-2)

A dPCR assay was designed to determine *RLK-2* expression in relation to the infection with *O. cumana* at different developmental stages (dpi). The ANOVA revealed a significant difference for the genotype (DEB2-resistant vs B117-susceptible) (F [1, 35] = 62.895, p < 0.001). In contrast, no significant main effects were observed for inoculation (F [1, 35] = 2.11, p = 0.155) and time of sampling (F [1, 35] = 0.132, p = 0.719) (Figure 5). The *RLK-2* gene was expressed at 14-dpi in inoculated (normalised expression ratio = 2.85 ± 0.8) and non-inoculated (1.69 ± 0.61) plants as well as at 21-dpi in inoculated (2.75 ± 0.43) and non-inoculated (2.14 ± 0.13) plants. As expected, no expression of the DEB2 *RLK-2* specific allele was found in the susceptible B117 in any of the conditions. Raw data are presented in Figure S8.

**Figure 5.**
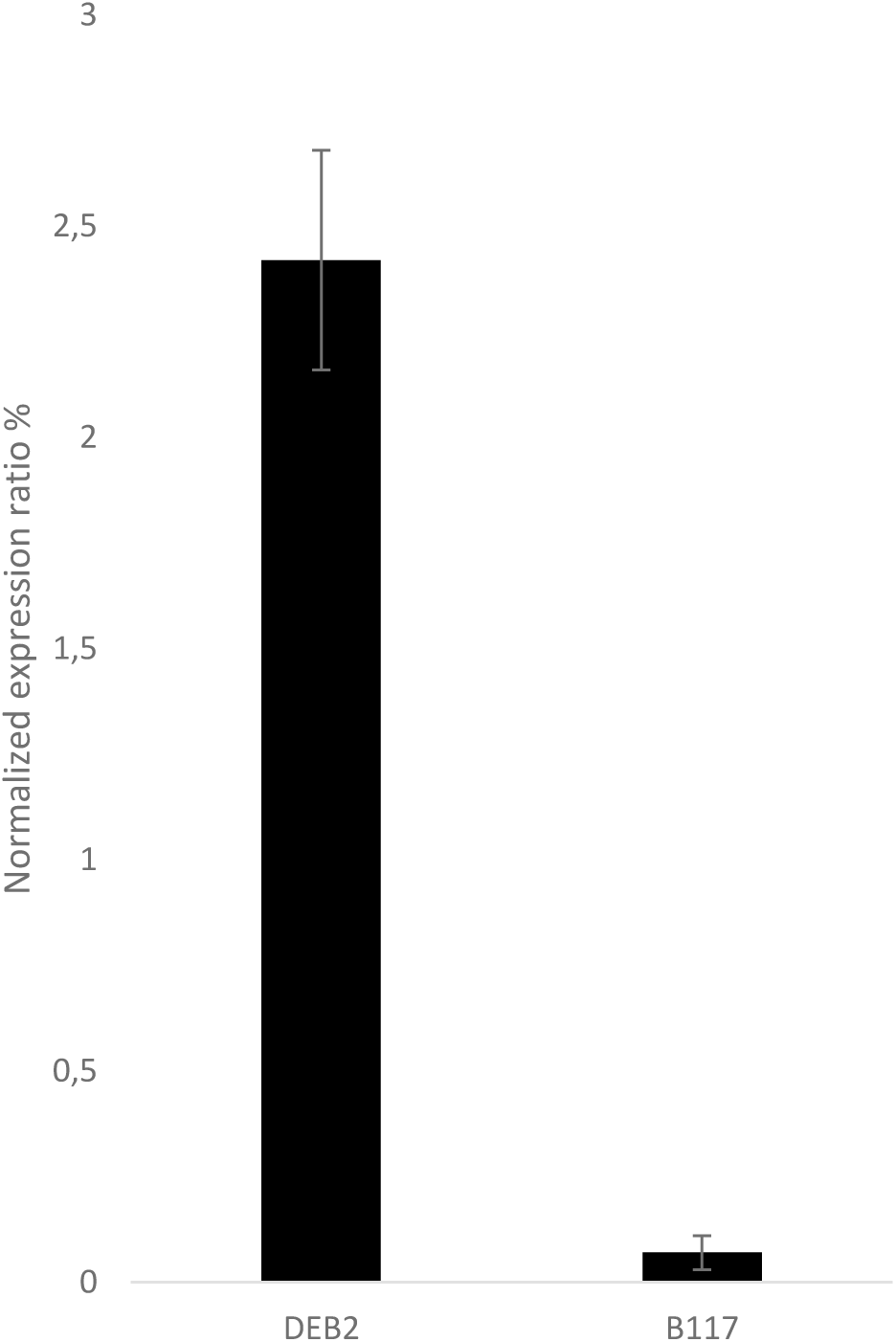
Expression profile of the *HannDeb2Chr04g005720* (*RLK-2*) gene in the two sunflower genotypes DEB2 and B117. All the conditions for each sunflower genotype (broomrape inoculation and date of sampling) are presented together, since no significant differences were found for these treatments. The data correspond to average plus minus standard error from 3 independent biological replicates of each condition from two different technical replicates.

### b2) Expression of *Helianthus debilis* subsp. *tardiflorus* PI 468691 genes in the *Or_Deb2_*-corresponding fine-mapped region

Expression of the three *H. debilis* subsp. *tardiflorus* PI 468691 genes annotated in the *Or_Deb2_*-corresponding fine-mapped region (Table S4B) was assessed by BLAST against the *H. debilis* transcripts using the IsoSeq dataset generated in this study (Table S5). *Hdeb2414_s0002g00045311* (orthologue to full-length *RLK-2*) was detected in transcripts from all three tissues analysed, with higher expression in the stem compared to buds and leaves. Regarding *Hdeb2414_s0002g00045321* (orthologue to *HSP20*), it was detected only in bud transcripts at a low level, with no hits in stem or leaf transcripts. Finally, transcripts of *Hdeb2414_s0002g00045341* (orthologue to truncated RLK) were not detected in any tissue.

### c) Comparative analysis of *Or_Deb2_* candidate sequence with its orthologues in several other sunflower genomes

For this analysis, the DEB2 full-length *RLK-2* in the fine-mapped region was compared to orthologous kinase genes in i) the B117 susceptible genotype used as parental line, ii) the lines XRQ, PSC8, HA89 and LR1 evaluated against broomrape and verified as also being susceptible to race G of *O. cumana* (Calderón-González et al., 2023 and this study), and iii) other sunflower lines and wild accessions whose genomes are available but not reported to be evaluated against race G of *O. cumana*.

All sunflower genomes had a variable number of RLKs, both from the Pelle-CrRLK1L-1 and Pelle-RLCK-VIIa-2 families (Figure 2). These genes were less represented in the PSC8-type genomes (with 1 or 2 copies for CrRLK1L-1 or 0 to 3 for RLCK-VIIa-2) than in the XRQ-type genomes (with 3 to 5 copies for CrRLK1L-1 or 1 to 3 for RLCK-VIIa-2). *RLK-2* was compared though BlastP to the kinases present in the AX-105525205 to AX-105399507 interval (*Or_Deb2_* large locus region) of the genomes of sunflower lines and wild species. Orthologous protein groups with high identity were found and aligned (Figure 6). The first one was a full-length orthogroup (OrtG) (named as OrtG 1) with 100% coverage and >90% identity, which included two and one RLKs of the reference genomes HanXRQ2.0 and HanPSC8, respectively, two of the parental genotype B117, one from the sunflower lines HA89, IR, LR1, RHA438, and one or three, respectively, from the accessions of wild species *H. debilis* PI 468691, and *H. annuus* PI659440. All the genes in OrtG 1 were annotated as RLK-Pelle CrRLK1-L1 protein kinases and shared a tandem kinase-pseudokinase domain structure.

**Figure 6.**
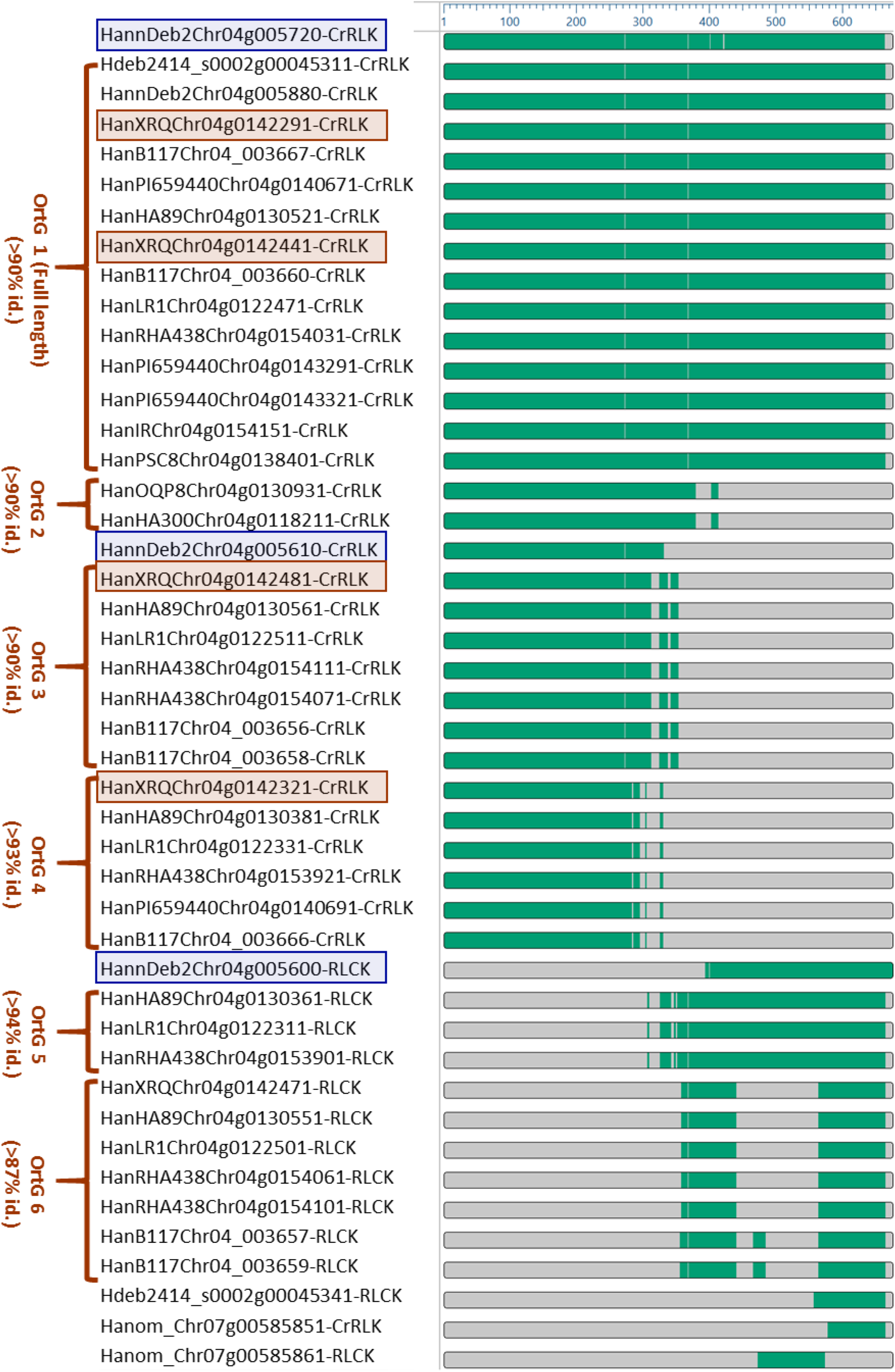
RLK conservation across sunflower genomes. Clustal Omega alignment of DEB2 RLKs within the *Or_Deb2_* fine-mapped interval [including the full-length kinase RLK-2 (*HannDeb2Chr04g005720*) and the truncated kinase RLK-1 (*HannDeb2Chr04g005600* and *HannDeb2Chr04g005610*)] with orthologues from cultivated lines (XRQ, PSC8, RHA438, IR, HA89, LR1, OQP8, HA300, B117) and wild relatives (*H. annuus* PI659440, *H. anomalus* PI468642, *H. debilis* PI468691) is shown, highlighting sequence conservation and structural divergence among full-length and truncated RLKs.

In addition to these full-length aligned genes, other orthologues to *RLK-2* (identity >87% to >94%) were found, but coding for truncated proteins with a reduced aligned fragment, most of them having less than 400 aa (Figure 6). Proteins in OrtG 2, 3, and 4 had a complete left-side kinase domain with an active site. Those in OrtG 5 and 6 had the right-side kinase domain, complete or interrupted, with no active site (pseudokinase domain). The two annotated genes corresponding to the truncated *RLK-1*, *HannDeb2Chr04g005610* and *HannDeb2Chr04g005600* were similar, respectively, to OrtG 2, 3, 4 and to OrtG 5.

Detailed amino acid sequence alignment analysis showed that RLK-2 presented unique variations characterised by two amino acid substitutions and two deletions of 2 and 1 amino acids, respectively (red arrows in Figure S9). These variations were not found in any of the genotypes susceptible to race G of *O. cumana* analysed, which included the parental genotype B117 and the lines XRQ (this study), PSC8 (this study), HA89 (Calderón-González et al., 2023) and LR1 (Calderón-González et al., 2023), and also the DEB2 RLK found outside the fine-mapped region (*RLK-4*).

LTR-retrotransposons were found to interrupt RLK genes, leading also to truncated proteins (Figure 7). This was the case, among others, for truncated genes in OrtG 3 and 6, represented by the HanXRQ genes *HanXRQr2Chr04g0142481* and *HanXRQr2Chr04g0142471*, respectively, which constitute two parts of an RLK gene interrupted by a retrotransposon (Figure 7). In DEB2, the gene *RLK-3*, contiguous to the *Or_Deb2_* fine-map interval, also had a retrotransposon inside (Figure 7) leading to a truncated protein, as explained also in previous sections (Figure 4).

**Figure 7.**
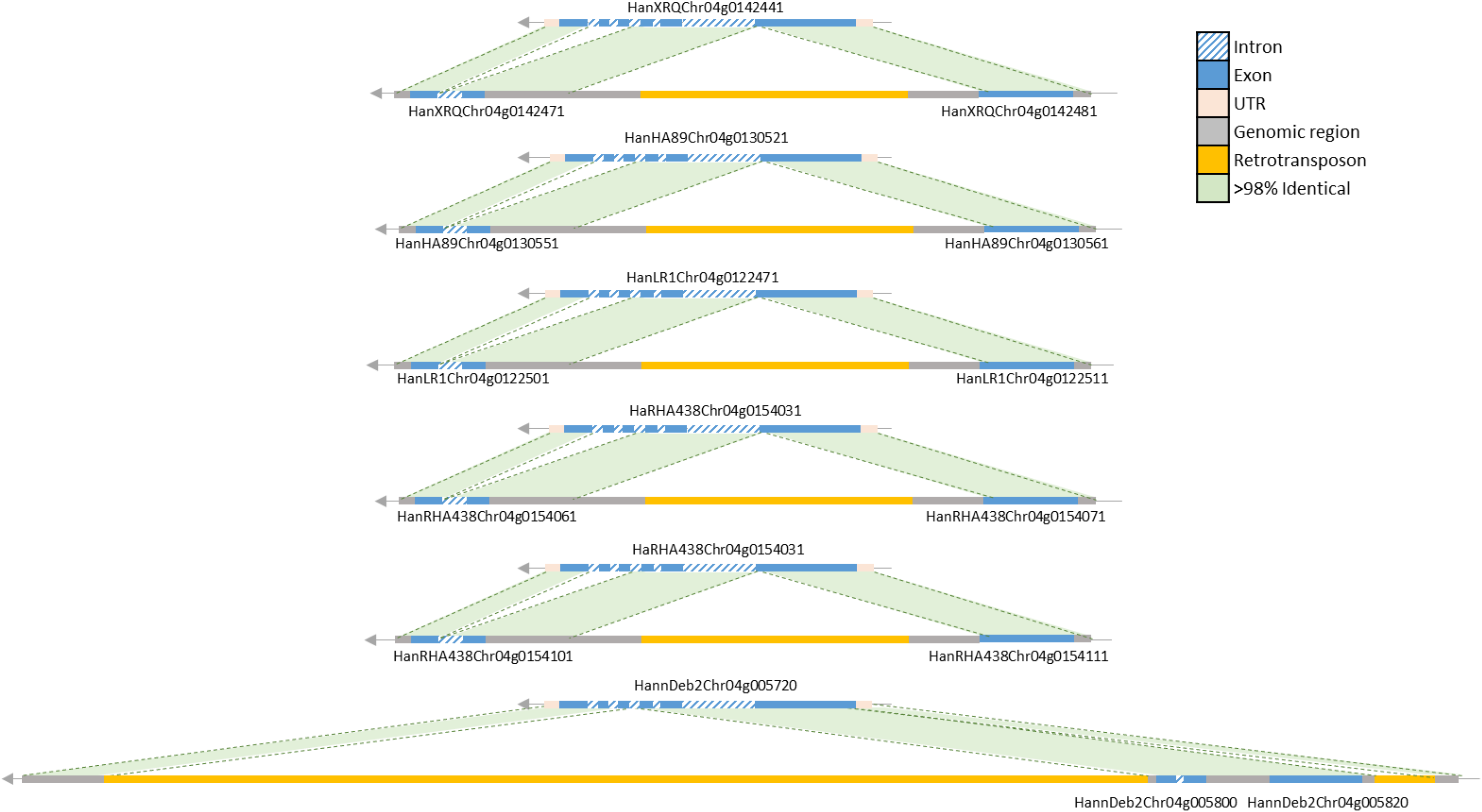
Transposon-mediated truncation of RLKs. Comparison of the full-length kinase *HanXRQChr04g0142441* with the truncated copies *HanXRQChr04g0142471* and *HanXRQChr04g0142481*, separated by a transposon, in the sunflower reference genome assembly HanXRQr2.0. The region highlighted in green indicates >98% nucleotide identity between the sequences. A similar structure, with a full-length kinase disrupted into two truncated copies by a transposon, is also present in other sunflower genomes (HanHA89, HanLR1, and HanRHA438, where it occurs twice) and in the DEB2 assembly (HannDeb2.20230803).

## Discussion

The DEB2 genotype carrying *Or_Deb2_* originated from a cross between a resistant accession of the wild sunflower species *H. debilis* subsp. *tardiflorus* (PI468691) and the cultivated susceptible *H. annuus* cultivar HA89 (Velasco et al., 2012). Comparison of the Chr 4 genomic sequences of DEB2, HA89 and *H. debilis* PI468691 revealed that a 17-Mbp segment at the top of Chr 4 in DEB2 was introgressed from *H. debilis*, the remainder of the chromosome identified as originating from HA89. This result is consistent with the *Or_Deb2_*gene introgressed from *H. debilis* and its position in the upper part of Chr 4, as reported by Fernández-Aparicio et al. (2022). Since the content of an introgression is not necessarily predictable from a reference sequence (Hübner et al. 2019), the genomic interval of the *Or_Deb2_* locus delimited by the SNP markers AX-105525205 and AX-105399507 defined in the initial studies of Fernández-Aparicio et al. (2022) has been analysed in this research in detail in DEB2, and compared to all sunflower genomes published to date and the *H. debilis* and B117 genome assemblies developed in this research. Marker order as well as orthologous genes such as small heat shock proteins HSP20, acyl-CoA desaturases, RNA directed DNA polymerases, ubiquitin conjugating enzyme E2, RLPs and RLKs were overall conserved across the different genome assemblies. Within the DEB2 genome assembly, the AX-105525205 and AX-105399507 markers delimited a 1.5-Mbp interval with 42 annotated genes, among which RLPs and RLKs of the RLK-Pelle family were found. The region containing the RLKs was quadruplicated in this genome, which led to four RLKs sharing high homology: *RLK-1* (truncated), *RLK-2* (full-length), *RLK-*3 (truncated) and *RLK-4* (full-length).

Through analysis of recombination points in the fine-mapping population, the *Or_Deb2_* gene has been fine-mapped to a 219.4-kbp region of the Chr 4-DEB2 genome assembly, which contains six protein-coding genes. They belong to two protein families: heat shock proteins (HSP20; 3 annotated genes) and RLK-Pelle proteins (3 annotated genes). HSP20 proteins act as molecular chaperones, preventing undesirable protein-protein interactions and assisting in the refolding of denatured proteins (Waters and Vierling, 2020). As such, they have been reported to influence defence responses to invading pathogens by protecting other proteins from stress-induced damage (Liu et al., 2018; Maimbo et al., 2007; van Ooijen et al., 2010). However, this relationship is not associated with gene-for-gene interactions, as is the case with the *O. cumana*-sunflower system, but rather with a broader innate immune mechanism (Lopes-Caitar et al., 2016). To our knowledge, no cloned major-effect resistance gene has been identified in plants as a heat shock protein (Chen et al., 2024). By contrast, the only gene cloned to date in sunflower that confers resistance to *O. cumana* is an RLK (Duriez et al., 2019). This fact, together with the central role of RLK genes in plant resistance acting as major genes that perceive pathogen and damage signals to trigger defence responses (Dievart et al., 2020; Klymiuk et al., 2021; Sung et al., 2025) make the RLK-Pelle genes in the fine map interval the best candidates for *Or_Deb2_*. In this interval, two RLKs were found: the full-length gene *RLK-2* (annotated as *HannDeb2Chr04g005720*), and the truncated gene *RLK-1* (annotated as *HannDeb2Chr04g005600* + *HannDeb2Chr04g005610*). Multiple lines of evidence, as discussed below, point to the *RLK-2* gene as the most plausible candidate for *Or_Deb2_*.

*RLK-2* encoded a full-length and fully expressed TKP. The RLK-2 protein was made up of two fused kinase domains: kinase and pseudokinase, classified, respectively, as RLK-Pelle CrRLKL1L-1 and RLK-Pelle RLCK-VIIa-2 types. Reveguk et al. (2025), in a survey of TKPs across the plant kingdom, found that CrRLKL1L-1_RLCK-VIIa-2 domain structure was only present in the species from the Eudicotyledoneae clade. Interestingly, among the 68 dicot species analysed in that study, *Lactuca sativa*, representative of the sunflower order Asterales, had the second-highest number of TKPs (96 in total). Additionally, BLASTp survey of DEB2 RLK-2 identified direct homologs (score > 500) across 23 Asteraceae species, showing a wide distribution of TKPs across the sunflower family.

Recently, KFPs have emerged as new key players in plant immunity (Klymiuk et al., 2021). In *Triticeae* species, these represent about 10% of cloned major resistance genes, most of which encode tandem kinases (Reveguk et al., 2025). They included for example the tandem kinase genes *Rpg1* (RPG1 protein) conferring resistance to many pathotypes of the stem rust fungus *Puccinia graminis* f. sp. *tritici* in barley (Brueggeman et al., 2002), and the wheat genes *Yr15* (Wheat Tandem Kinase 1-WTK1) (Klymiuk et al., 2018), *Sr60* (WTK2) (Chen et al., 2020), *Pm24* (WTK3) (Lu et al., 2020) or *WTK4* (WTK4) (Gaurav et al., 2022) conferring resistance to, respectively, yellow rust (*Puccinia striiformis* f. sp. *tritici*), stem rust (*Puccinia graminis* Pers. f. sp*. tritici*), and powdery mildew (*Blumeria graminis* f. sp. *tritici*; *Pm24* and *WTK4*). *RLK-2* shares the same tandem kinase structure as these genes, with *Yr15* (WTK1) sharing the highest similarity. Although phylogenetically distant, both RLK-2 and WTK1 have kinase and pseudokinase domains of the RLK-Pelle family and show a conserved active site at the kinase domain and no transmembrane or membrane-targeting motifs. The lack of these last motifs is consistent with the experimentally validated cytoplasmic localisation of WTK1 (Klymiuk et al., 2018). It is important to note that in WTK1 both kinase and pseudokinase domains are required for resistance, as mutations in either domain render plants susceptible (Klymiuk et al., 2018).

Another piece of evidence supporting a tandem kinase gene as the best candidate for *Or_Deb2_* is that functionally validated tandem-kinase genes that confer resistance to fungal diseases in cereals share important genetic features with *Or_Deb2_*. First, they represent genes with a dominant, monogenic inheritance pattern, e.g., *Yr15* (Gerechter-Amitai et al. 1989), *Pm24* (Huang et al. 2000) and *Rwt4* (Hau et al. 2007). Second, the pathogens to which they confer resistance are characterised by the presence of physiological races, i.e., populations with defined virulence patterns against specific host genotypes, as it is also characteristic of the parasitic weed *O. cumana* (Molinero-Ruiz et al., 2015). This is, for example, the case for wheat yellow rust (Ali et al., 2017), wheat stem rust (Singh et al., 2015) or wheat powdery mildew (Kang et al., 2020). Finally, in both cereal/fungal pathogen and sunflower/*O. cumana* interactions, genetic evidence supports the key role of this class of genes in gene-for-gene relationships, which involve direct or indirect recognition of a pathogen avirulence (Avr) gene product by the corresponding plant resistance (R) gene product. This is supported by genetic studies in *O. cumana* (Rodríguez-Ojeda et al., 2013; Calderón-González et al., 2024) and has been experimentally validated for the wheat resistance gene RWT4 (Sung et al., 2025).

Analysis of candidate gene allele sequences in susceptible and resistant sunflower genotypes provided additional evidence supporting *RLK-2* as the gene underlying *Or_Deb2_*. The *RLK-2* gene was compared with its full-length tandem kinase orthologous genes in the B117 susceptible parental line and in other genomes of genotypes susceptible to race G of *O. cumana*, namely XRQ, PSC8, HA89, and LR1. It was also compared to the DEB2 full-length RLK *RLK-4*, located outside the fine-mapped interval. The DEB2 allele of *RLK-2* showed a distinctive protein sequence characterised by two unique amino acid substitutions and two deletions, absent in all susceptible genotypes and in DEB2-*RLK-4*. It is interesting to note that all the susceptible genotypes have the full-length RLK orthologues, which indeed are confirmed to be expressed in B117 (this study) and in XRQ (XRQ transcriptome; Badouin et al., 2017), so it seems that the susceptible allele might not be a defective gene coding for a truncated protein, as was the case for *HaOr7-RLK* (Duriez et al., 2019).

The role of *RLK-2* as the candidate gene for resistance to broomrape in DEB2 is further supported by the fact that the only cloned major gene that confers resistance to *O. cumana* in sunflower (*HaOr7*) is also an RLK (Duriez et al., 2019). *HaOr7* encodes a membrane-bound receptor protein having an extracellular LRR domain and an intracellular kinase domain (LRR-RLK protein). By contrast, RLK-2 lacks an LRR domain and has two fused kinase domains. To date, there are very few cloned genes conferring resistance to parasitic plants. Two of them are cell-surface localised receptors, including the above mentioned *HaOr7* LRR-RLK, and the *CuRe1* gene, which confers resistance to *Cuscuta reflexa* in tomato (*Solanum lycopersicum*) and encodes a LRR-RLP with an extracellular LRR domain and no kinase intracellular domain (Hegenauer et al., 2016). A third one is an intracellular nucleotide-binding domain leucine-rich repeat (NLR) immune receptor (RSG3-301), that confers resistance to *Striga gesnerioides* in cowpea (*Vigna unguiculata*) (Li and Timko, 2009). *Or_Deb2_*-*RLK-2* represents a new likely intracellular receptor involved in plant immune response against a parasitic weed. Recent scientific findings have highlighted the significance of tandem kinase proteins as intracellular receptors, with RWT4 reported to directly bind a fungal effector (Sung et al., 2025), and WTK3/*Rwt4* or Sr62/*Sr62* described to establish cooperative intracellular “sensor-executor” mechanisms with NLRs in mediating fungal resistance pathways (Chen et al., 2025; Lu et al., 2025).

Other sunflower genes conferring resistance to race G of *O. cumana* have been mapped to Chr4, tightly linked to *Or_Deb2_*. These include the *Or_SII_*(Martín-Sanz et al., 2020) and *Or_Anom1_* (Fernández-Melero et al., 2024) genes from PHSC1102 and ANOM1 resistant sunflower genotypes, respectively. These resistance genes differed from *Or_Deb2_* in their associated resistance mechanism, with *Or_SII_* and *Or_Anom1_*showing a late post-attachment response that impeded broomrape development after vascular connections with the host are established (Martín-Sanz et al., 2020; Fernández-Melero et al., 2024), while that in *Or_Deb2_* was characterised by an early post-attachment system that blocks parasite intrusion at the root cortex level (Fernández-Aparicio et al., 2022). In the case of ANOM1, the resistance gene is known to be introgressed from the wild sunflower relative *H. anomalus* accession PI 468642. Particularly, *Or_Anom1_* has been mapped between SNP markers Iasnip-41 and Iasnip-17 (Fernández-Melero et al., 2024), which also delimitate the *OrDeb2* larger locus (Figure 1). Although the *Or_Anom1_*has not been fine-mapped or the ANOM1 genome sequenced, the study of this region in the sequenced *H. anomalus* accession ANO-2822 also identified two RLK-Pelle genes as putative *Or_Anom1_* candidates. The characterization of the ANOM1 genome sequence, which is underway, would contribute to determine if *Or_Anom1_* might represent a different allele of the DEB2-*RLK-2* gene and the role of these genes in broomrape resistance.

Defective tandem kinase gene structures coding for truncated proteins have been found in DEB2 and in all the studied genomes in the present study. In DEB-2, the other RLK annotated within the fine-mapped region (*RLK-1*), highly homologous to *RLK-2* (> 99% of nucleotide homology), showed a full-length genomic sequence that, when translated, contained a stop codon close to the end of the first exon, resulting in a truncated protein. This has also been observed for the susceptible allele of the sunflower RLK gene *HaOr7*, which contains a stop codon that results in a truncated protein lacking the intracellular kinase domain (Duriez et al., 2019). In addition to stop codons, defective conformations of the RLK genes were also found to be created by retrotransposon insertions in DEB2, as well as in other sunflower genomes such as HanXRQ, HanHA89, HanLR1 and HanRHA438, resulting also in truncated kinases. These interruptions cause gene mutations that most often lead to truncated, non-functional proteins, as documented in sunflower, for example, for downy mildew resistance genes (Franchel et al., 2013) and for genes associated with tocopherol biosynthesis (Tang et al., 2006).

In addition to transposon insertion, localized gene duplications appeared to be a mechanism underlying the expansion and evolution of the *Or_Deb2_*-associated RLKs. The *Or_Deb2_* region containing these candidates was found to be quadruplicated explicitly in the DEB2 genome, which led to three RLKs sharing high homology (*RLK-1*, *RLK-2* and *RLK-4*) and a fourth RLK also with high homology but truncated by a transposon (*RLK-3*). Tightly linked full-length and truncated RLKs were also found across all the genomes studied. This kind of gene rearrangements generating split and shorter or, contrarily, fused and longer RLKs associated with duplications and and/or transposon element insertions, have been described as one of the primary genetic events causing the expansion of the RLK-Pelle gene family (Shiu and Bleecker, 2003; Shiu et al., 2004; Zhang et al., 2005). In addition, tandem kinase-pseudokinase genes, as characterised by having two fused kinase domains with high sequence similarity, have also been proposed to have originated by duplication events (Klymiuk et al., 2021).

Although DEB2 was developed from the introgression of the broomrape resistance trait from *H. debilis* subsp. *tardiflorus* PI468691, differences at the genomic level between these two genotypes within the *Or_Deb2_*-containing region have been detected. Among them, a significant quadruplicated region, mentioned above, was observed in DEB2 but not in *H. debilis* PI468691. Also, the full-length orthologue TKP found in the *H. debilis* genome within the *Or_Deb2_*-corresponding fine-mapped interval (*HdebPI468691s0002g00045311*) presented slight differences at the amino acid sequence level in relation to DEB2 full-length TKP RLK-2 (Figure S9). Since the *H. debilis* accession PI468691 used for genome sequencing is resistant to sunflower broomrape (Chabaud et al., 2022), this may suggest that the two genome assemblies might share the same genomic configuration and the same resistant allele. The observed genomic differences at the *Or_Deb2_* locus may be associated with noticeable intra-accession diversity present in wild sunflower species, which are obligate outcrossers due to self-incompatibility systems that lead to divergence and heterogeneity (Seiler et al., 2017), particularly considering that the wild *H. debilis* accession PI468691 did not co-evolve with sunflower broomrape. Whereas *O. cumana* is naturally distributed through south-western Asia and south-eastern Europe (Pujadas-Salvà and Velasco, 2000), the genus *Helianthus* is native to North America (Seiler et al., 2017). In particular, *H. debilis* subsp. *tardiflorus* was collected in Florida (https://npgsweb.ars-grin.gov/gringlobal/accessiondetail?id=1363627), whereas *O. cumana* was never detected in the American Continent (Parker, 2013) until very recently, with a first report in Bolivia in 2023 (Barea et al., 2025). Thus, the resistance gene from North American *H. debilis* PI 468691 had no interaction with *O. cumana* in the wild, escaping long-term co-evolutionary dynamics in natural settings, and as a consequence the *Or_Deb2_*locus configuration has not been influenced by selection pressure posed by the presence of the parasite.

## Conclusion

The *Or_Deb2_* gene has been located within a genomic section of 219.4 kbp from Chr 4 of the DEB2 genome assembly, as determined by high-resolution genetic and physical mapping. Within this region, a full-length and root-expressed tandem kinase-pseudokinase gene (*RLK-2*) has been proposed as the most plausible *Or_Deb2_* candidate and will be named *HaOr_Deb2_-TKP-2*. This study provides the first evidence suggesting a role of TKPs in disease resistance within dicot plants and in gene-for-gene interactions between crops and parasitic weeds. The results also provide an opportunity to gain a deeper understanding of the genomic organization, function, and evolution of these gene types in the sunflower genome. Finally, they will support sunflower breeding programs aimed at marker-assisted introgression of *O. cumana* resistance genes into elite sunflower germplasm, and identify of new allelic variants conferring resistance, contributing to building more durable and sustainable breeding strategies based on genetic resistance and in-depth knowledge of the sunflower-*O. cumana* parasitic system.

## Acknowledgements

We thank Plácida Nieto and Alberto Merino (IAS-CSIC, Córdoba, Spain) for technical support, and INRAe GENTYANE platform of Clermont-Ferrand INRAE Center (https://gentyane.clermont.inrae.fr/) and CNRGV (Centre National de Ressources Génomiques Végétales) INRAe platform (Toulouse, France) (https://cnrgv.toulouse.inrae.fr/) for providing assistance in NGS sequencing.

## Competing interests

We declare no conflicts of interest in regard to this manuscript.

## Author contributions

BFM developed SNP and FLP markers, fine-mapped the resistance gene, analysed the genomic structure of the *Or_Deb2_* locus in several sunflower genotypes, and performed rhizotron experiments, cloning of RLK genes, expression analyses, and phenotypic assays. BPV developed the Iasnip SNP markers ranging from 16 to 55 and contributed to initial cloning DEB2 and B117 fragments for marker design. LdM contributed to sample collection, DNA extraction, genotyping, and cloning. LV coordinated all phenotyping assays. SM contributed to genotyping and coordinated DEB2 genome sequencing. AL prepared sunflower DNA for sequencing different sunflower genomes. JG and SC generated the *H. debilis* genome data and performed genome assembly and annotation. BFM produced the B117 genome assembly and annotation. BFM and BRP contributed to the DEB2 genome annotation and conducted the introgression study. BPV and LV designed the experiments. BPV coordinated the overall study and contributed to phenotypic assays and overall data analyses. BPV, LV, and BFM wrote the manuscript. All authors revised the manuscript and approved the final version.

## Data availability

PCR primer and probe sequences, SNP context sequences, amino acid alignments among RLKs from different sunflower genomes, the genotyping dataset from informative recombinants, and raw expression analysis data are provided as Supplementary Data. The *H. debilis* PI468691 whole-genome shotgun project has been deposited at DDBJ/ENA/GenBank under the accession number JBPZNU000000000. The version described in this paper is JBPZNU010000000. The DEB2 whole-genome shotgun project has been deposited at DDBJ/ENA/GenBank under the accession JBPDJB000000000. The version described in this paper is version JBPDJB010000000. The B117 Chr 4 genomic interval delimited by SNP markers AX-105525205 and AX-105399507 (referred to as B117_OrDeb2_AX) is available as Supplementary Data.

## Funding

The study was funded by research project PID2020-117286RB-I00 of the Spanish Ministry of Economy and Innovation (co-funded with EU FEDER Funds) and was supported by the grant to Belén Fernández-Melero PRE2018-084486 funded by MCIN/AEI/10.13039/501100011033 and ESF “ESF investing in your future” and Junta de Andalucía (Spain) Qualifica Project QUAL21_023 IAS.

## Supporting Information

### Supplementary files

**File S1.** Fasta file for the B117 genomic interval between markers AX-105525205 and AX-105399507 containing the *Or_Deb2_* locus (B117_*Or_Deb2_*_AX interval).

### Supplementary tables

**Table S1.** Sequences and their position in the HannDeb2.20230803-Chr04, HanXRQ2.0-Chr04, HanPSC8-Chr04 and HdebPI468691r0-s0002 genome assemblies of SNP and PCR-based markers used for fine mapping *Or_Deb2_*.

**Table S2.** Sequence of PCR primers used for cDNA amplification for cloning the *Or_Deb2_* candidate gene *HannDeb2Chr044g005720* and sequence of primers and probes for dPCR assay for the expression analysis.

**Table S3.** F_2_ and F_3_ genotyping score of the five FMDEB-57/58_59/60 to Iasnip-50/55 recombinant plants (R1 to R5). A= DEB2 (resistant) allele; B= B117 (susceptible) allele, H= Heterozygous, D= A or H. F_2_ score for DEB2 dominant markers was based on the genotyping of the corresponding derived F_3_ families, in such a way that the F_2_ was scored as B when the F_3_ was fully homozygous B (no amplification at any individual); H when F_3_ segregated and D when the whole F_3_ was A or H (all individuals showed amplification so no distinction can be made between homozygous A or heterozygous). The phenotype was set according to the number of broomrapes emerged per plant: R= 0 broomrapes; S = 1 or more broomrapes. Lost data are represented with a hypen (-).

**Table S4. (A)** DEB2 protein-coding genes annotated in the HannDeb2.20230803 assembly between markers AX-105525205 and AX-105399507. Genes located within the interval FMDEB-59/60 and FMDEB-476/477 are marked as *Or_Deb2_*fine-mapped region. Protein length and domain structure of the genes annotated as CrRLK are specified. InterPro domains related to retroelements are annotated. Grey marks indicate retrotransposon annotation by LTRharvest. Annotated genes with clear evidence of being retrotransposons domains and non-protein-coding genes were excluded from the analysis and are marked in light grey. **(B)** Gene annotation of the *Helianthus debilis* subsp. *tardiflorus* PI468691 assembly (HdebPI468691r0) between markers AX-105525205 and AX-105399507. Genes located within *Or_Deb2_*-corresponding-fine-mapped region are highlighted. Annotated genes with clear evidence of being retrotransposons domains and non-protein-coding genes are marked in light grey.

**Table S5.** Iso-Seq analysis of *H. debilis* genes within the *Or_Deb2_* fine-mapped region. BLAST results of reference mRNA *H. debilis* sequences against Iso-Seq transcripts obtained from bud, stem, and leaf tissues is shown. Hits with ≥80% coverage of both query and subject sequences are highlighted in green.

### Supplementary figures

**Figure S1.** Schematic representation of the different genotyping rounds of the fine-mapping population with SNP and PCR-based markers. White flowers represent susceptible plants. Black flowers represent resistant plants.

**Figure S2**. Dot plots of HannDeb2.20230803 assembly aligned with **(A)** HanHA89-Chr04 and **(B)** HdebPI468691-s0002 showing the introgressed 17 Mbp region.

**Figure S3.** Dot plots comparing the genomic region within markers Iasnip-41 and Iasnip-16 **(A)** between the *H. annuus* genome assembly HanXRQr2.0-SUNRISE and the DEB2 genome assembly HannDeb2.20230803, and **(B)** the HannDeb2.20230803 assembly against itself. Quadruplicated region of HannDeb2.20230803 is indicated with a square and each duplicated segment is indicated with a number.

**Figure S4.** Comparative synteny analysis of the protein-coding genes from the *Or_Deb2_* fine mapped region in DEB2 and their putative orthologs in *H. debilis*. Only orthologous protein pairs with ≥88% sequence identity are shown. Conserved gene order, structural relationships, and gene duplications are illustrated by connecting lines between orthologous proteins.

**Figure S5.** Sequence alignments of the cloned RLK cDNAs from B117 and DEB2. **(A)** Alignment of the reference sequences with their corresponding cloned cDNA sequences, illustrating overall sequence conservation and exon–intron structure. **(B)** Nucleotide alignments at higher resolution, highlighting the correspondence between the nucleotides of each sequence.

**Figure S6.** Protein architecture of the full-length kinases HannDeb2Chr04g0005720 (RLK-2) from DEB2 and B117Chr04_003660 from B117. **(A)** Schematic representation of the HannDeb2Chr04g0005720 (RLK-2) protein domains. KD = Kinase domain; PKD = Pseudokinase domain. Orange lines indicate ATP-binding motifs, and the red line marks the predicted active site. **(B)** HannDeb2Chr04g0005720 (RLK-2) InterProScan domain annotation. The protein contains two kinase-related domains: a canonical kinase domain (KD) and a putative pseudokinase domain (PKD). The domain organization includes conserved domains, predicted intrinsically disordered regions, conserved residues, and active site, and binding site predictions. Functional classification is supported by Pfam, SMART, and other domain databases. **(C)** B117Chr04_003660 InterProScan domain annotation.

**Figure S7.** Comparison of *HannDeb2Chr04g005720* with the other validated TKPs with the same domain structure known to date. **(A)** Protein domain structure using ScanProsite is shown. Red diamond indicates active site (D residue). Grey diamond with a green line indicates the conserved ATP binding domain (K residue). **(B)** Amino acid sequence alignment between HannDeb2Chr04g005720 and WTK1. Asterisks (*) indicate identical residues; colons (:) indicate strongly similar residues; periods (.) indicate weakly similar residues; and the absence of a symbol indicates no similarity.

**Figure S8.** dPCR analysis for level of expression of *HannDeb2Chr04g005720* (*RLK-2*). **(A)** The table summarizes the raw data from the full experiment, including all biological replicates, the two technical replicas and the two probes: HANNDEB2CHR04g005720-exp (blue) and ACT-exp (green). **(B)** The graphs display representative fluorescence amplitude plots from a single replicate from 1) DEB2 and 2) B117. In each plot, the blue line indicates the fixed threshold used across probes; droplets above the threshold are classified as positive, and those below as negative. Each row corresponds to a different sunflower line. NTC refers to the non-template control.

**Figure S9.** Amino acid alignment of the full-length RLK-2 (HannDeb2Chr04g005720) and the truncated RLK-1 kinase (HannDeb2Chr04g005600 + HannDeb2Chr04g005610) (highlighted with a blue square) and their orthologues from different sequenced sunflower genome assemblies. Unique variations in RLK-2 (HannDeb2Chr04g005720) are indicated with red arrows.

## Notes

### Competing Interest Statement

The authors have declared no competing interest.

