## Supplemental Figure S1 for "Fine mapping and genomic analyses reveal a tandem kinase-pseudokinase candidate underlying *Or_Deb2_*-mediated resistance to *Orobanche cumana* in sunflower"

### Slide 1
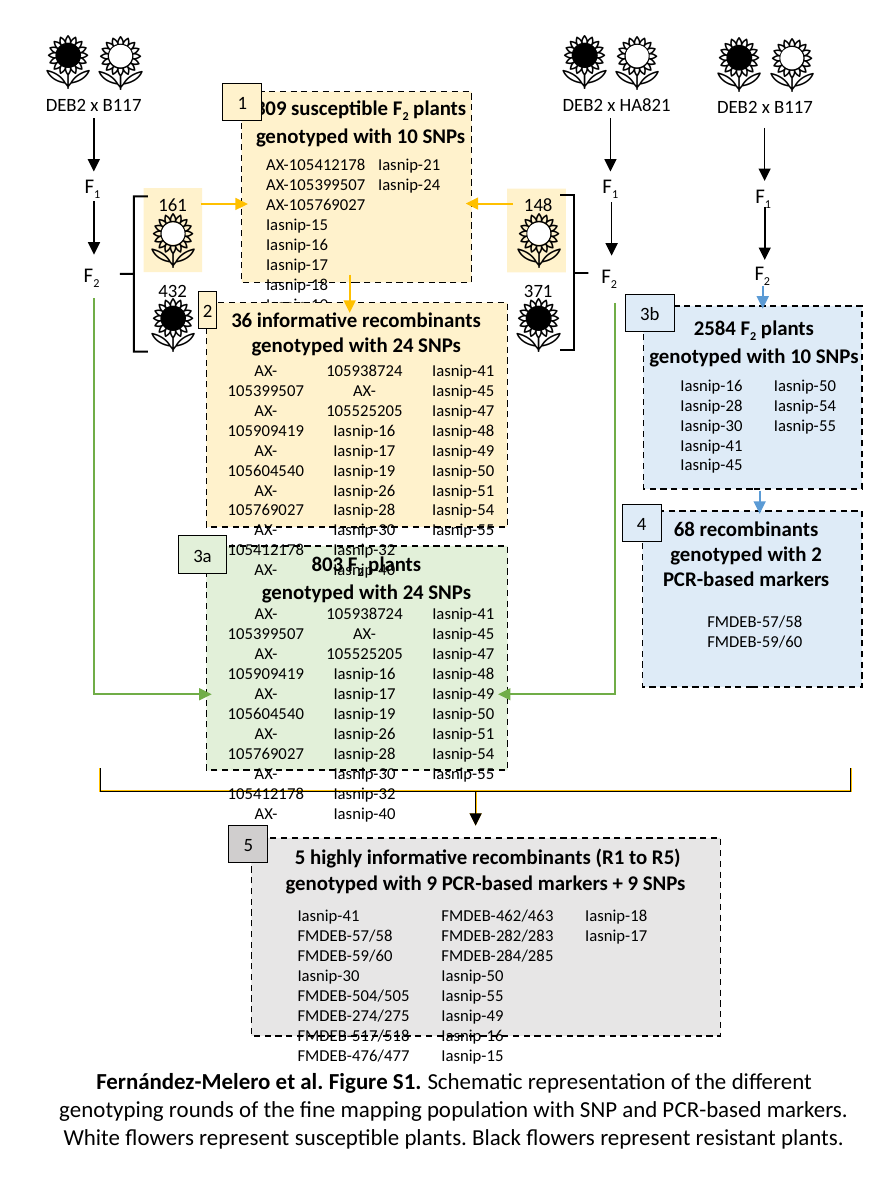

1
DEB2 x B117
DEB2 x HA821
DEB2 x B117
309 susceptible F_2_ plants
genotyped with 10 SNPs
AX-105412178
AX-105399507
AX-105769027
Iasnip-15
Iasnip-16
Iasnip-17
Iasnip-18
Iasnip-19
Iasnip-21
Iasnip-24
F_1_
F_1_
F_1_
161
148
F_2_
F_2_
F_2_
432
371
2
3b
36 informative recombinants
genotyped with 24 SNPs
2584 F_2_ plants
genotyped with 10 SNPs
AX-105399507
AX-105909419
AX-105604540
AX-105769027
AX-105412178
AX-105938724
AX-105525205
Iasnip-16
Iasnip-17
Iasnip-19
Iasnip-26
Iasnip-28
Iasnip-30
Iasnip-32
Iasnip-40
Iasnip-41
Iasnip-45
Iasnip-47
Iasnip-48
Iasnip-49
Iasnip-50
Iasnip-51
Iasnip-54
Iasnip-55
Iasnip-16
Iasnip-28
Iasnip-30
Iasnip-41
Iasnip-45
Iasnip-47
Iasnip-49
Iasnip-50
Iasnip-54
Iasnip-55
4
68 recombinants
genotyped with 2 PCR-based markers
3a
803 F_2_ plants
genotyped with 24 SNPs
AX-105399507
AX-105909419
AX-105604540
AX-105769027
AX-105412178
AX-105938724
AX-105525205
Iasnip-16
Iasnip-17
Iasnip-19
Iasnip-26
Iasnip-28
Iasnip-30
Iasnip-32
Iasnip-40
Iasnip-41
Iasnip-45
Iasnip-47
Iasnip-48
Iasnip-49
Iasnip-50
Iasnip-51
Iasnip-54
Iasnip-55
FMDEB-57/58
FMDEB-59/60
5
5 highly informative recombinants (R1 to R5) genotyped with 9 PCR-based markers + 9 SNPs
Iasnip-41
FMDEB-57/58
FMDEB-59/60
Iasnip-30
FMDEB-504/505
FMDEB-274/275
FMDEB-517/518
FMDEB-476/477
FMDEB-462/463
FMDEB-282/283
FMDEB-284/285
Iasnip-50
Iasnip-55
Iasnip-49
Iasnip-16
Iasnip-15
Iasnip-18
Iasnip-17
Fernández-Melero et al. Figure S1. Schematic representation of the different genotyping rounds of the fine mapping population with SNP and PCR-based markers. White flowers represent susceptible plants. Black flowers represent resistant plants.
