## Supplemental Figure S2 for "Fine mapping and genomic analyses reveal a tandem kinase-pseudokinase candidate underlying *Or_Deb2_*-mediated resistance to *Orobanche cumana* in sunflower"

### Slide 1
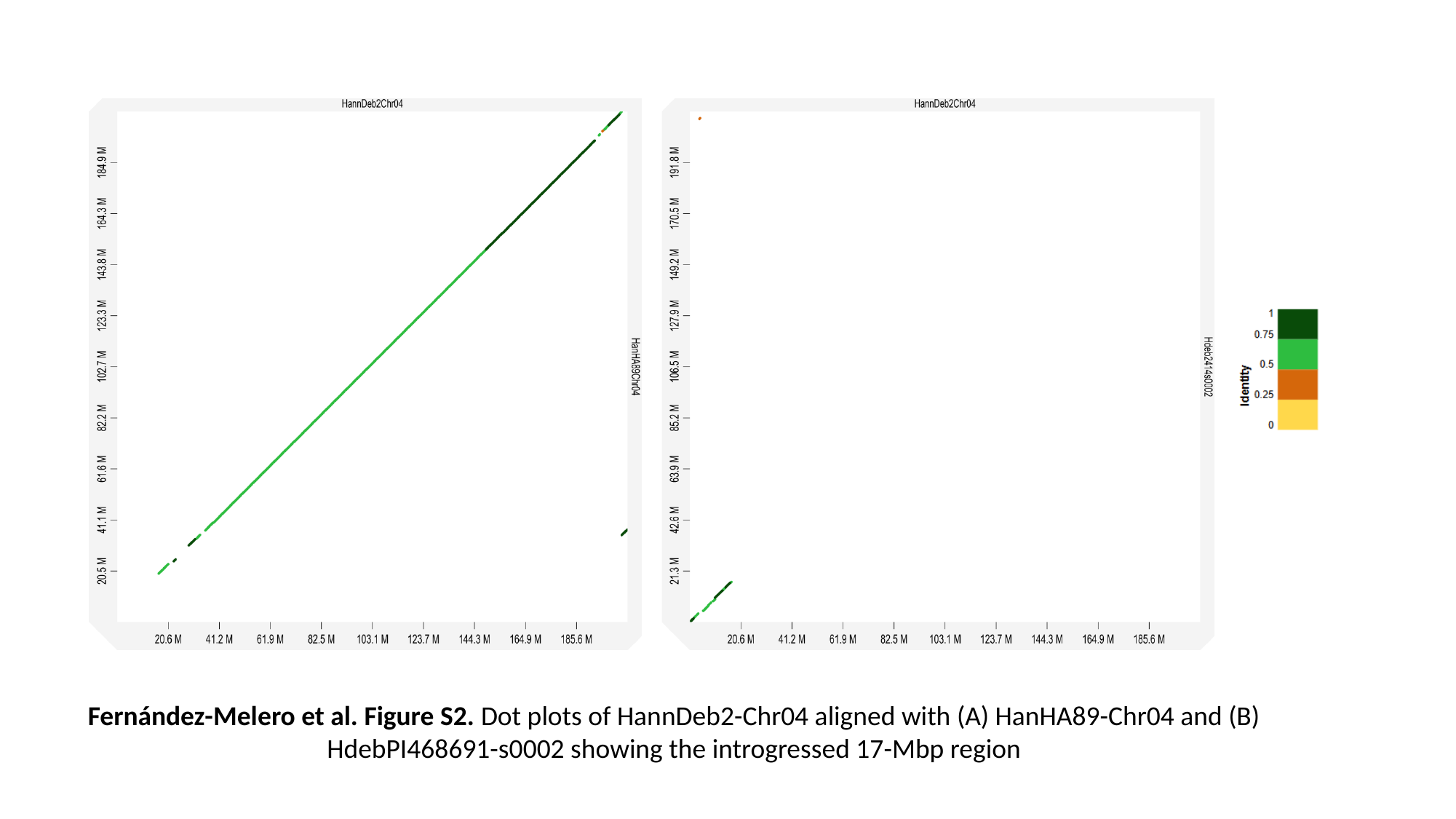

Fernández-Melero et al. Figure S2. Dot plots of HannDeb2-Chr04 aligned with (A) HanHA89-Chr04 and (B) HdebPI468691-s0002 showing the introgressed 17-Mbp region
