## Supplemental Figure S3 for "Fine mapping and genomic analyses reveal a tandem kinase-pseudokinase candidate underlying *Or_Deb2_*-mediated resistance to *Orobanche cumana* in sunflower"

### Slide 1
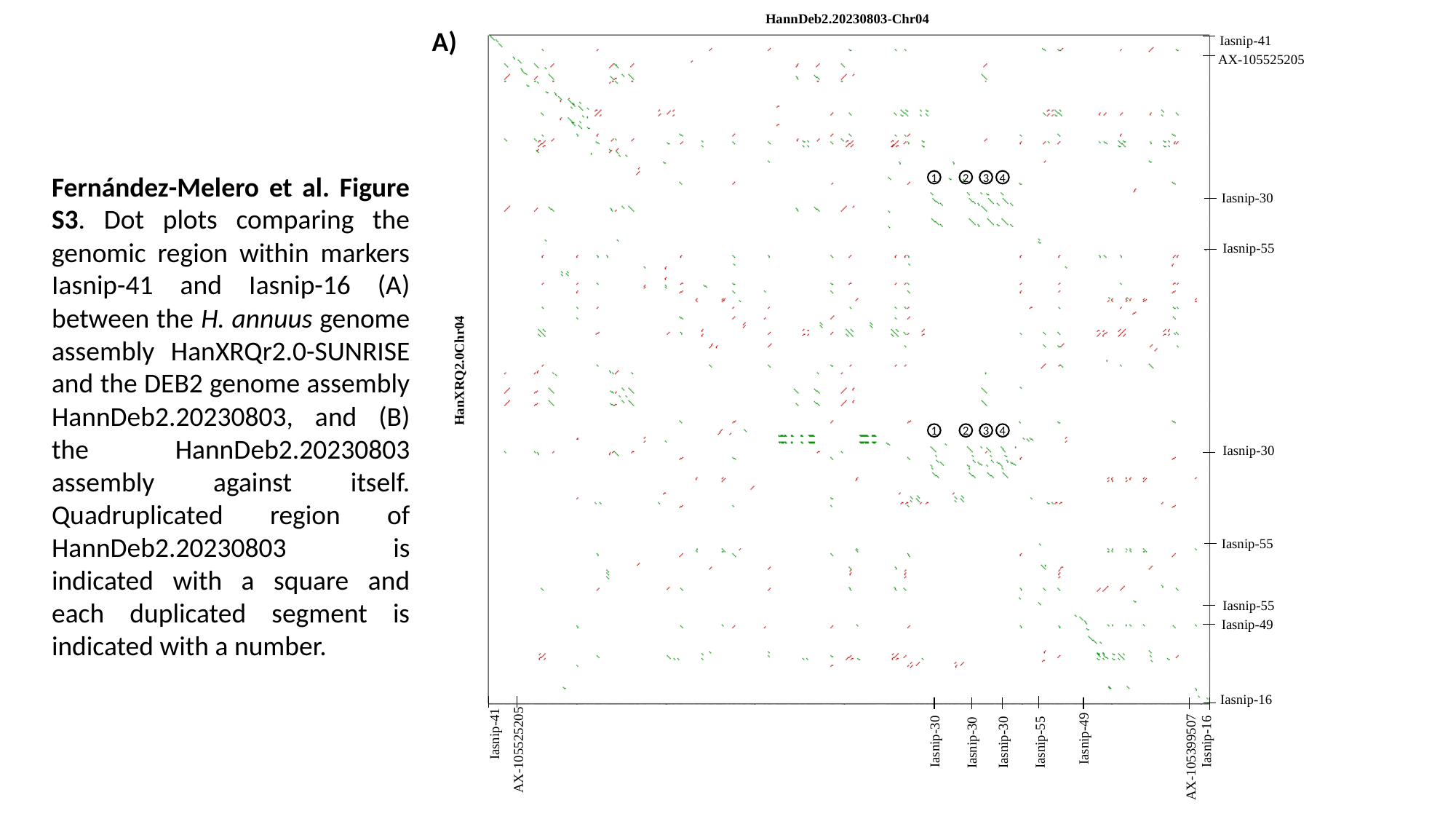

HannDeb2.20230803-Chr04
A)
Iasnip-41
AX-105525205
Fernández-Melero et al. Figure S3. Dot plots comparing the genomic region within markers Iasnip-41 and Iasnip-16 (A) between the H. annuus genome assembly HanXRQr2.0-SUNRISE and the DEB2 genome assembly HannDeb2.20230803, and (B) the HannDeb2.20230803 assembly against itself. Quadruplicated region of HannDeb2.20230803 is indicated with a square and each duplicated segment is indicated with a number.
Iasnip-30
Iasnip-55
HanXRQ2.0Chr04
Iasnip-30
Iasnip-55
Iasnip-55
Iasnip-49
Iasnip-16
Iasnip-41
Iasnip-49
Iasnip-16
Iasnip-30
Iasnip-55
Iasnip-30
Iasnip-30
AX-105525205
AX-105399507
1
2
3
4
1
2
3
4

### Slide 2
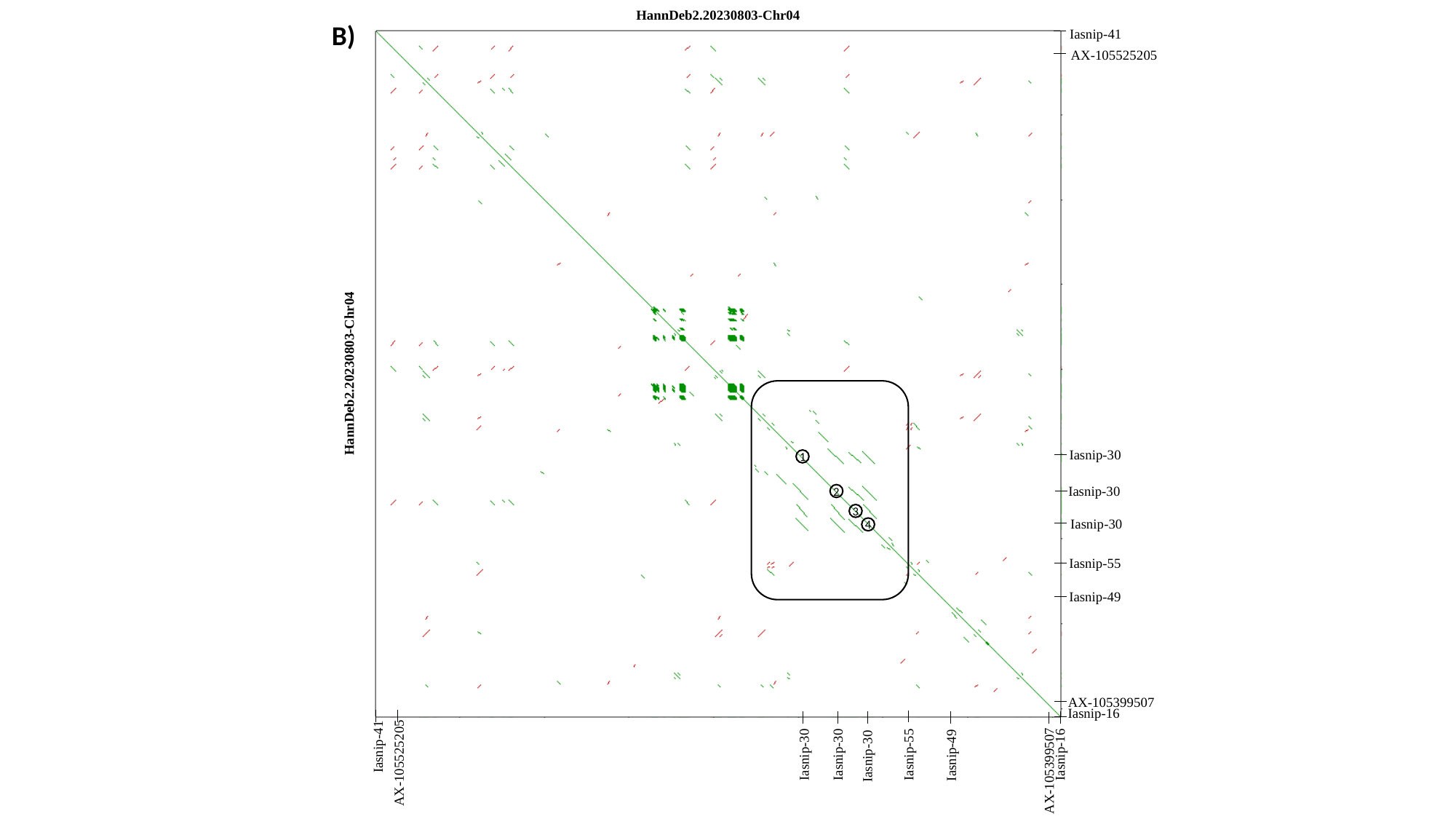

HannDeb2.20230803-Chr04
B)
Iasnip-41
AX-105525205
HannDeb2.20230803-Chr04
Iasnip-30
Iasnip-30
Iasnip-30
Iasnip-55
Iasnip-49
AX-105399507
Iasnip-16
Iasnip-41
Iasnip-55
Iasnip-30
Iasnip-16
Iasnip-30
Iasnip-49
Iasnip-30
AX-105525205
AX-105399507
1
2
3
4
