## Supplemental Figure S4 for "Fine mapping and genomic analyses reveal a tandem kinase-pseudokinase candidate underlying *Or_Deb2_*-mediated resistance to *Orobanche cumana* in sunflower"

### Slide 1
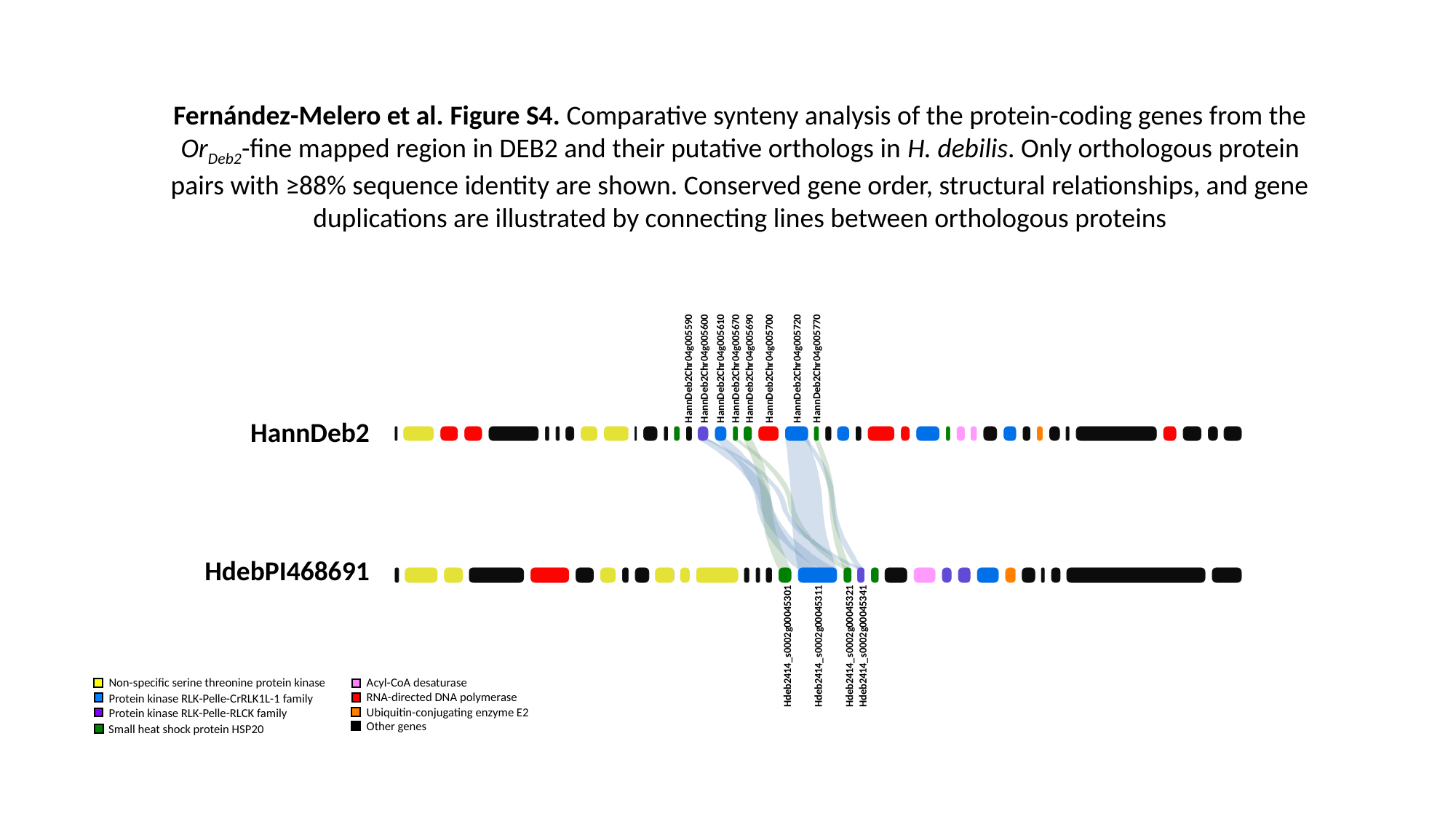

Fernández-Melero et al. Figure S4. Comparative synteny analysis of the protein-coding genes from the Or_Deb2_-fine mapped region in DEB2 and their putative orthologs in H. debilis. Only orthologous protein pairs with ≥88% sequence identity are shown. Conserved gene order, structural relationships, and gene duplications are illustrated by connecting lines between orthologous proteins
HannDeb2Chr04g005770
HannDeb2Chr04g005720
HannDeb2Chr04g005670
HannDeb2Chr04g005690
HannDeb2Chr04g005700
HannDeb2Chr04g005610
HannDeb2Chr04g005590
HannDeb2Chr04g005600
HannDeb2
HdebPI468691
Hdeb2414_s0002g00045341
Hdeb2414_s0002g00045311
Hdeb2414_s0002g00045321
Hdeb2414_s0002g00045301
Non-specific serine threonine protein kinase
Acyl-CoA desaturase
RNA-directed DNA polymerase
Protein kinase RLK-Pelle-CrRLK1L-1 family
Ubiquitin-conjugating enzyme E2
Protein kinase RLK-Pelle-RLCK family
Other genes
Small heat shock protein HSP20
