## Supplemental Figure S5 for "Fine mapping and genomic analyses reveal a tandem kinase-pseudokinase candidate underlying *Or_Deb2_*-mediated resistance to *Orobanche cumana* in sunflower"

### Slide 1
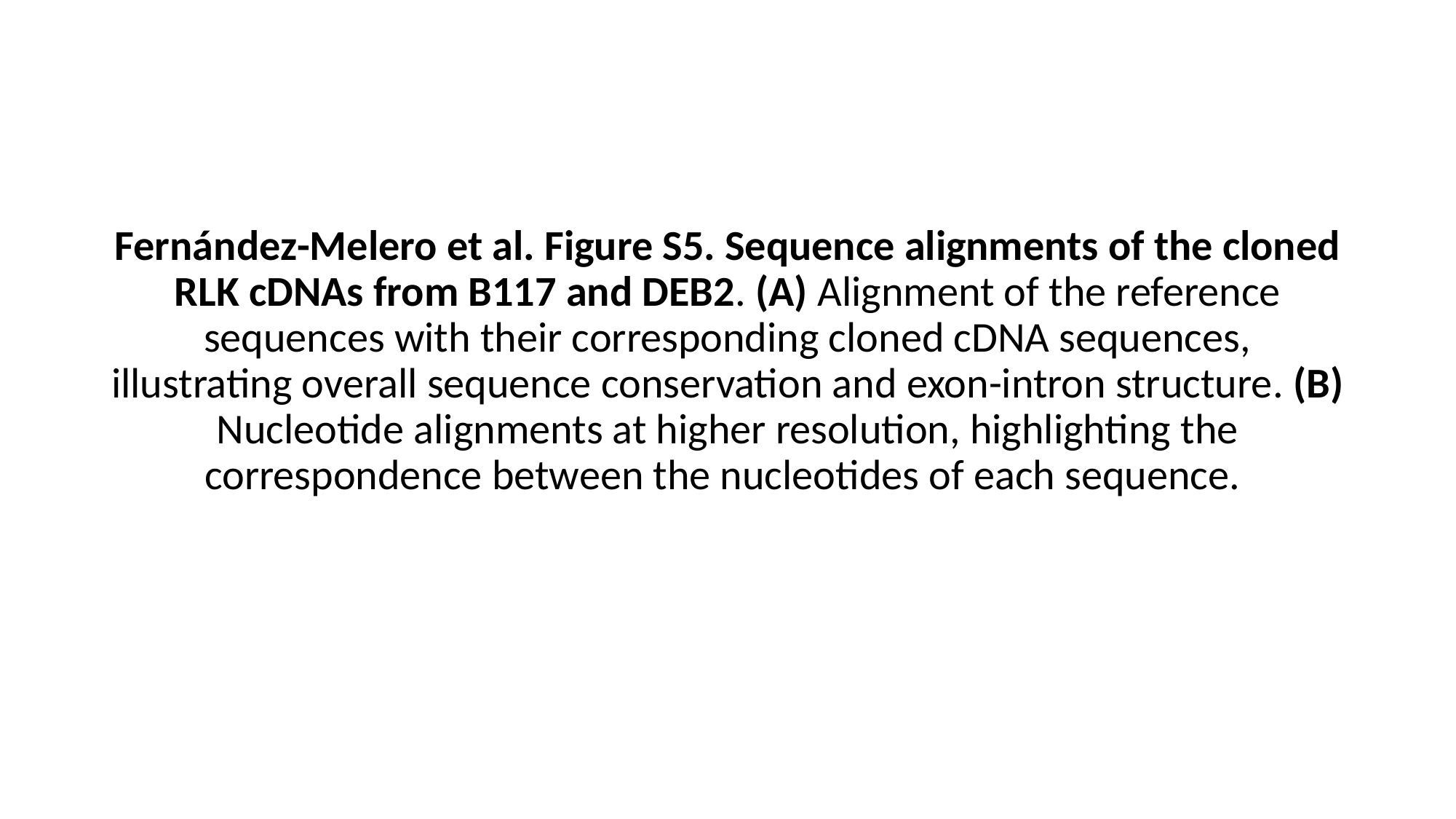

Fernández-Melero et al. Figure S5. Sequence alignments of the cloned RLK cDNAs from B117 and DEB2. (A) Alignment of the reference sequences with their corresponding cloned cDNA sequences, illustrating overall sequence conservation and exon-intron structure. (B) Nucleotide alignments at higher resolution, highlighting the correspondence between the nucleotides of each sequence.

### Slide 2
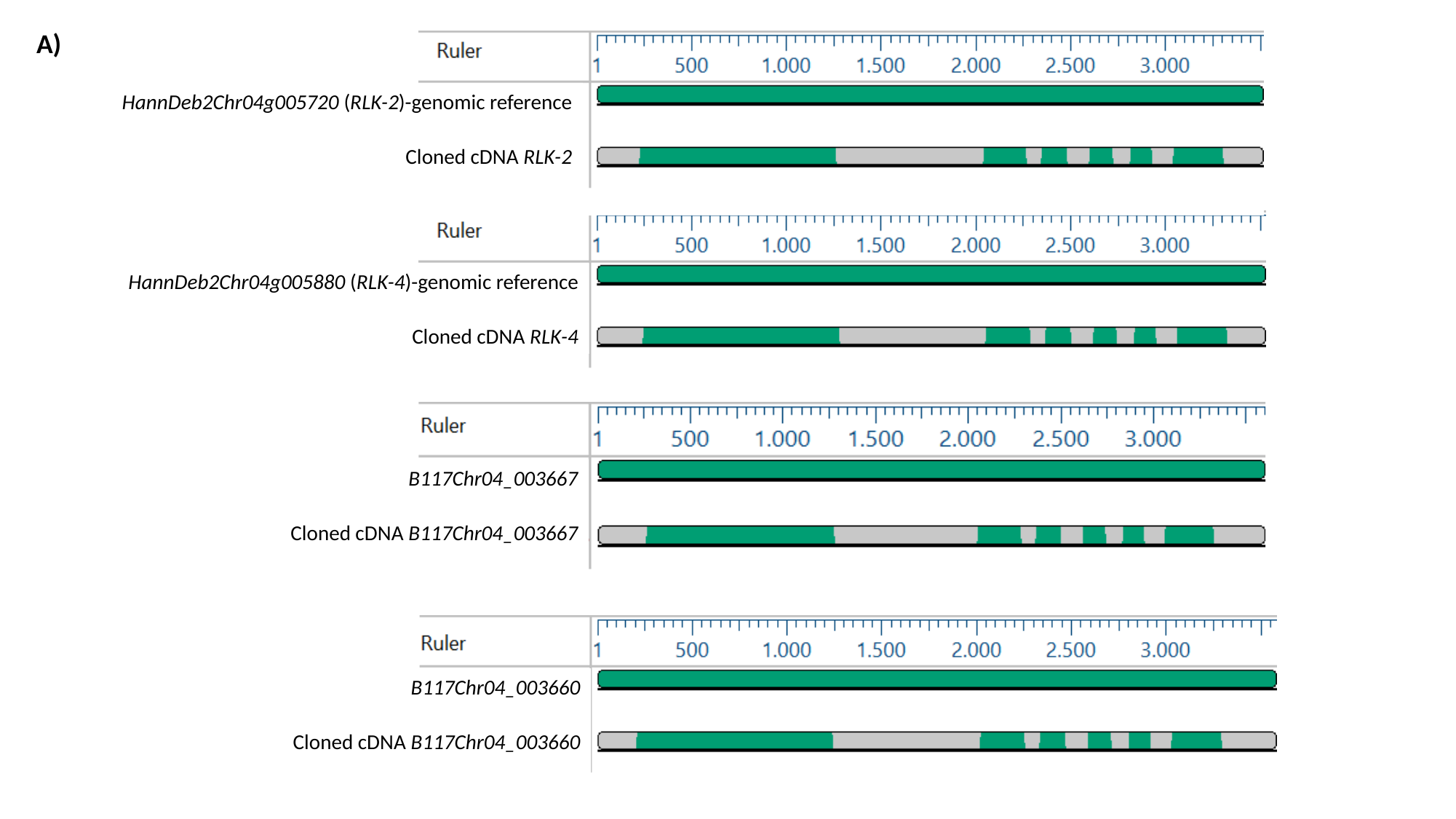

A)
HannDeb2Chr04g005720 (RLK-2)-genomic reference
Cloned cDNA RLK-2
HannDeb2Chr04g005880 (RLK-4)-genomic reference
Cloned cDNA RLK-4
B117Chr04_003667
Cloned cDNA B117Chr04_003667
B117Chr04_003660
Cloned cDNA B117Chr04_003660

### Slide 3
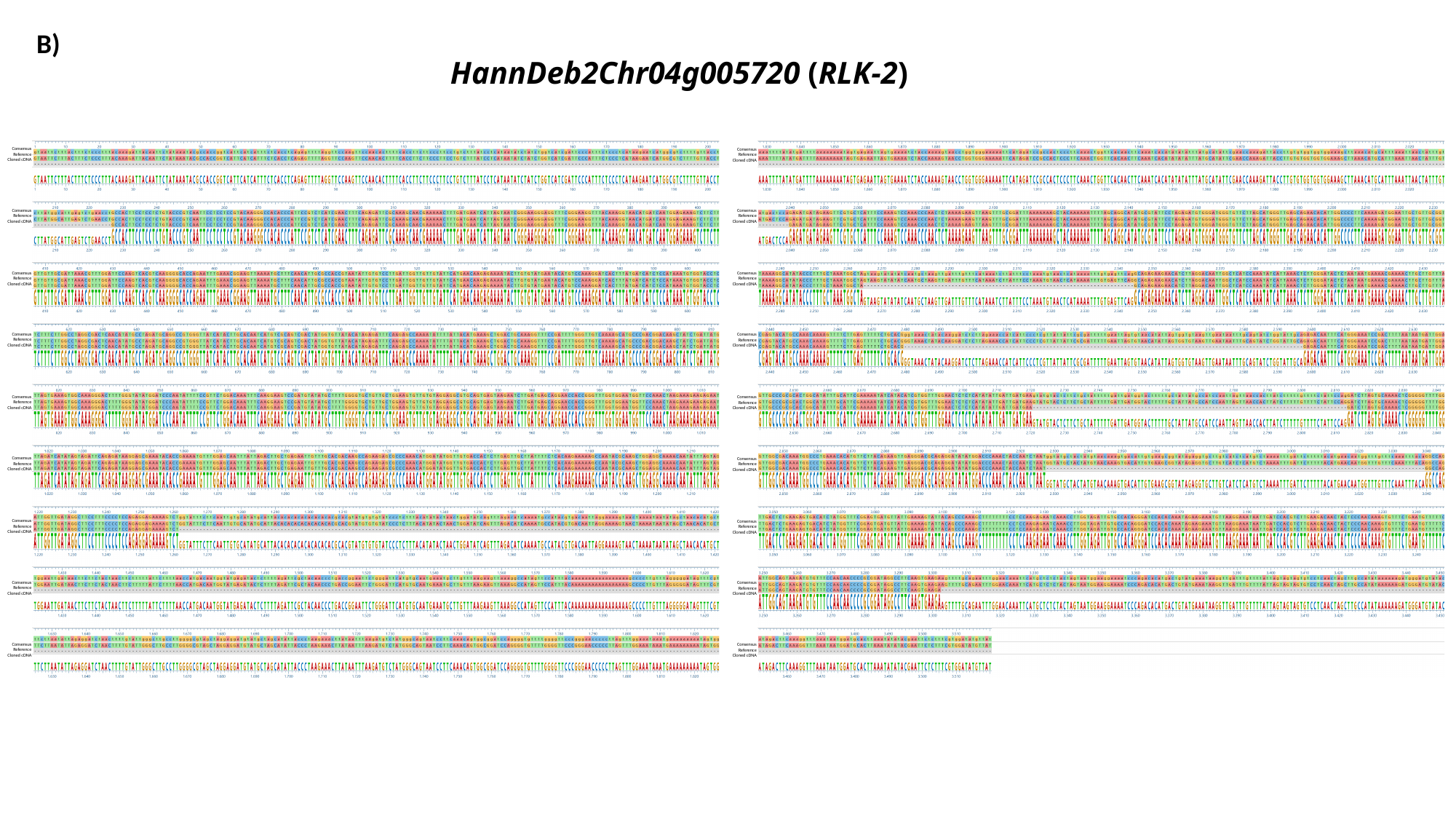

B)
#### HannDeb2Chr04g005720 (RLK-2)
Consensus
Reference
Cloned cDNA
Consensus
Reference
Cloned cDNA
Consensus
Reference
Cloned cDNA
Consensus
Reference
Cloned cDNA
Consensus
Reference
Cloned cDNA
Consensus
Reference
Cloned cDNA
Consensus
Reference
Cloned cDNA
Consensus
Reference
Cloned cDNA
Consensus
Reference
Cloned cDNA
Consensus
Reference
Cloned cDNA
Consensus
Reference
Cloned cDNA
Consensus
Reference
Cloned cDNA
Consensus
Reference
Cloned cDNA
Consensus
Reference
Cloned cDNA
Consensus
Reference
Cloned cDNA
Consensus
Reference
Cloned cDNA
Consensus
Reference
Cloned cDNA
Consensus
Reference
Cloned cDNA

### Slide 4
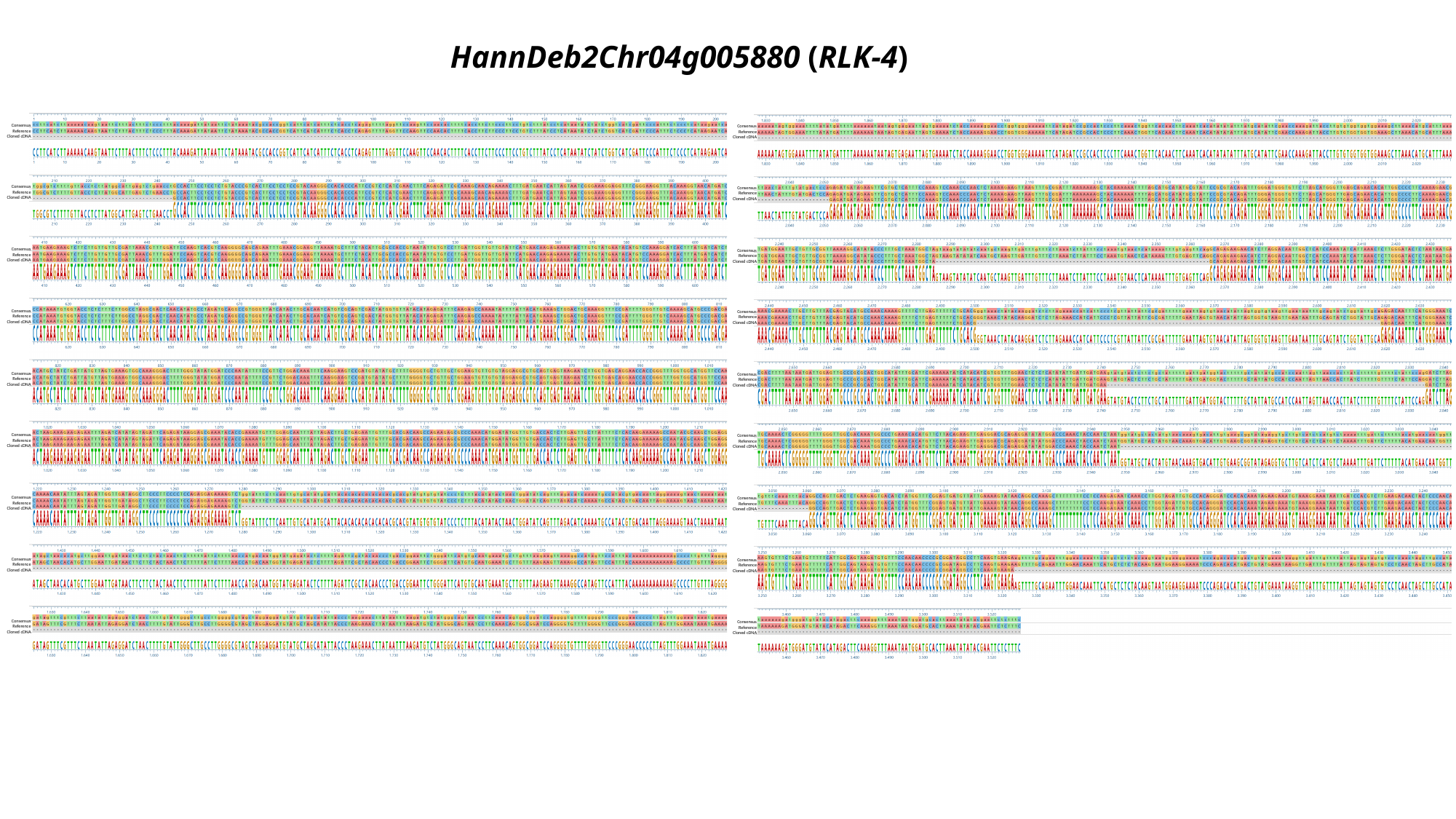

#### HannDeb2Chr04g005880 (RLK-4)
Consensus
Reference
Cloned cDNA
Consensus
Reference
Cloned cDNA
Consensus
Reference
Cloned cDNA
Consensus
Reference
Cloned cDNA
Consensus
Reference
Cloned cDNA
Consensus
Reference
Cloned cDNA
Consensus
Reference
Cloned cDNA
Consensus
Reference
Cloned cDNA
Consensus
Reference
Cloned cDNA
Consensus
Reference
Cloned cDNA
Consensus
Reference
Cloned cDNA
Consensus
Reference
Cloned cDNA
Consensus
Reference
Cloned cDNA
Consensus
Reference
Cloned cDNA
Consensus
Reference
Cloned cDNA
Consensus
Reference
Cloned cDNA
Consensus
Reference
Cloned cDNA
Consensus
Reference
Cloned cDNA

### Slide 5
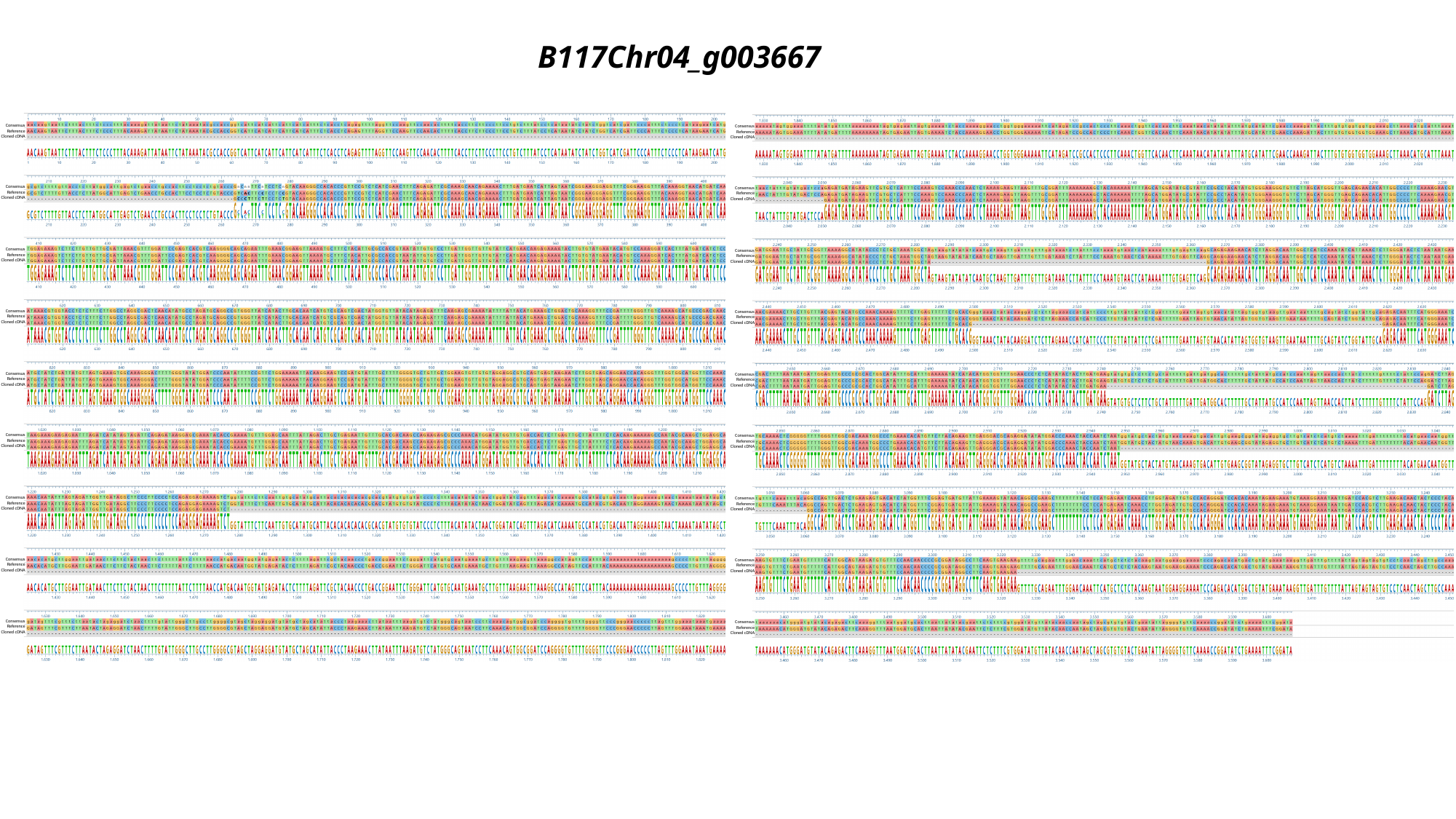

#### B117Chr04_g003667
Consensus
Reference
Cloned cDNA
Consensus
Reference
Cloned cDNA
Consensus
Reference
Cloned cDNA
Consensus
Reference
Cloned cDNA
Consensus
Reference
Cloned cDNA
Consensus
Reference
Cloned cDNA
Consensus
Reference
Cloned cDNA
Consensus
Reference
Cloned cDNA
Consensus
Reference
Cloned cDNA
Consensus
Reference
Cloned cDNA
Consensus
Reference
Cloned cDNA
Consensus
Reference
Cloned cDNA
Consensus
Reference
Cloned cDNA
Consensus
Reference
Cloned cDNA
Consensus
Reference
Cloned cDNA
Consensus
Reference
Cloned cDNA
Consensus
Reference
Cloned cDNA
Consensus
Reference
Cloned cDNA

### Slide 6
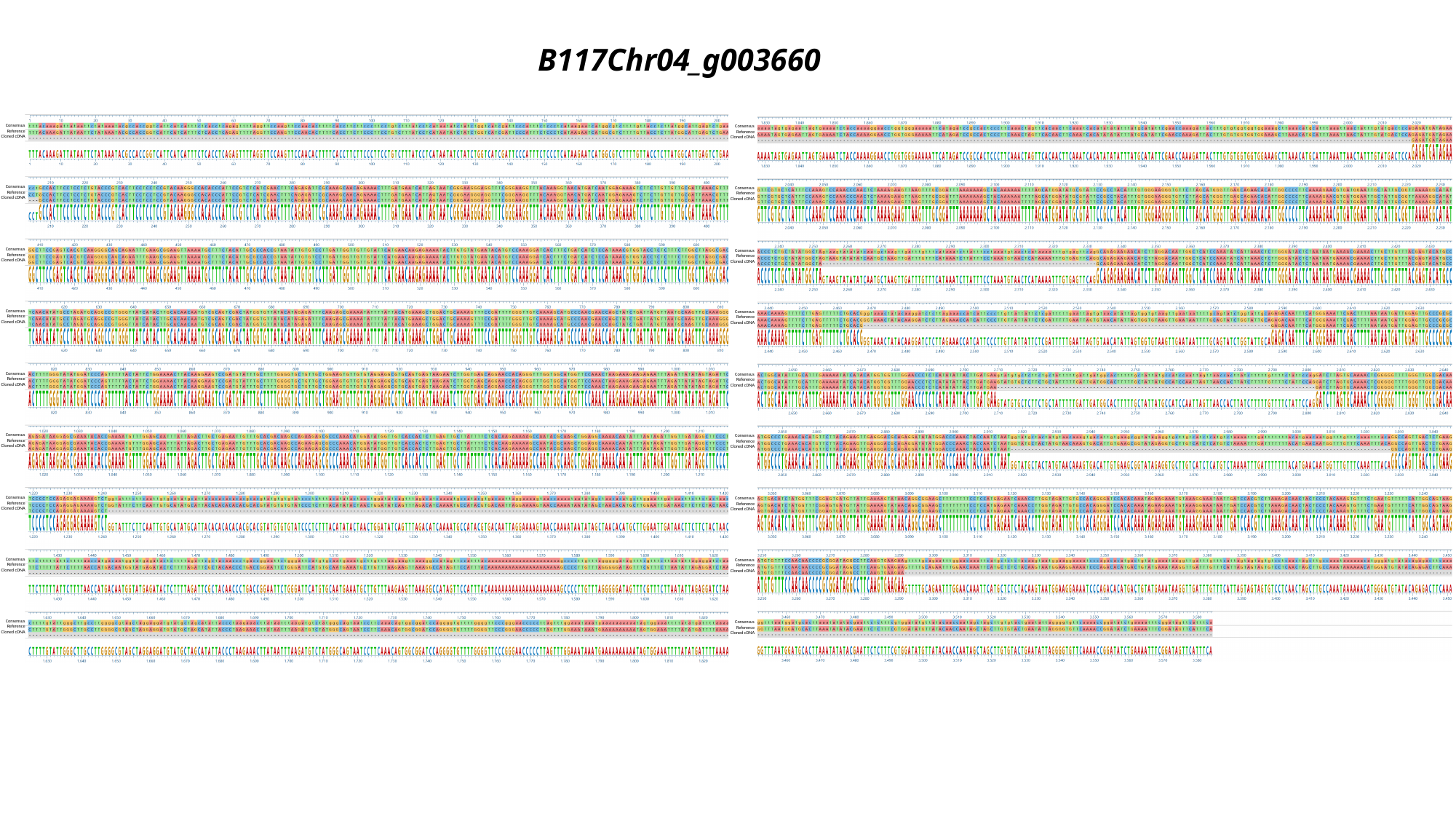

#### B117Chr04_g003660
Consensus
Reference
Cloned cDNA
Consensus
Reference
Cloned cDNA
Consensus
Reference
Cloned cDNA
Consensus
Reference
Cloned cDNA
Consensus
Reference
Cloned cDNA
Consensus
Reference
Cloned cDNA
Consensus
Reference
Cloned cDNA
Consensus
Reference
Cloned cDNA
Consensus
Reference
Cloned cDNA
Consensus
Reference
Cloned cDNA
Consensus
Reference
Cloned cDNA
Consensus
Reference
Cloned cDNA
Consensus
Reference
Cloned cDNA
Consensus
Reference
Cloned cDNA
Consensus
Reference
Cloned cDNA
Consensus
Reference
Cloned cDNA
Consensus
Reference
Cloned cDNA
Consensus
Reference
Cloned cDNA
