## Supplemental Figure S6 for "Fine mapping and genomic analyses reveal a tandem kinase-pseudokinase candidate underlying *Or_Deb2_*-mediated resistance to *Orobanche cumana* in sunflower"

### Slide 1
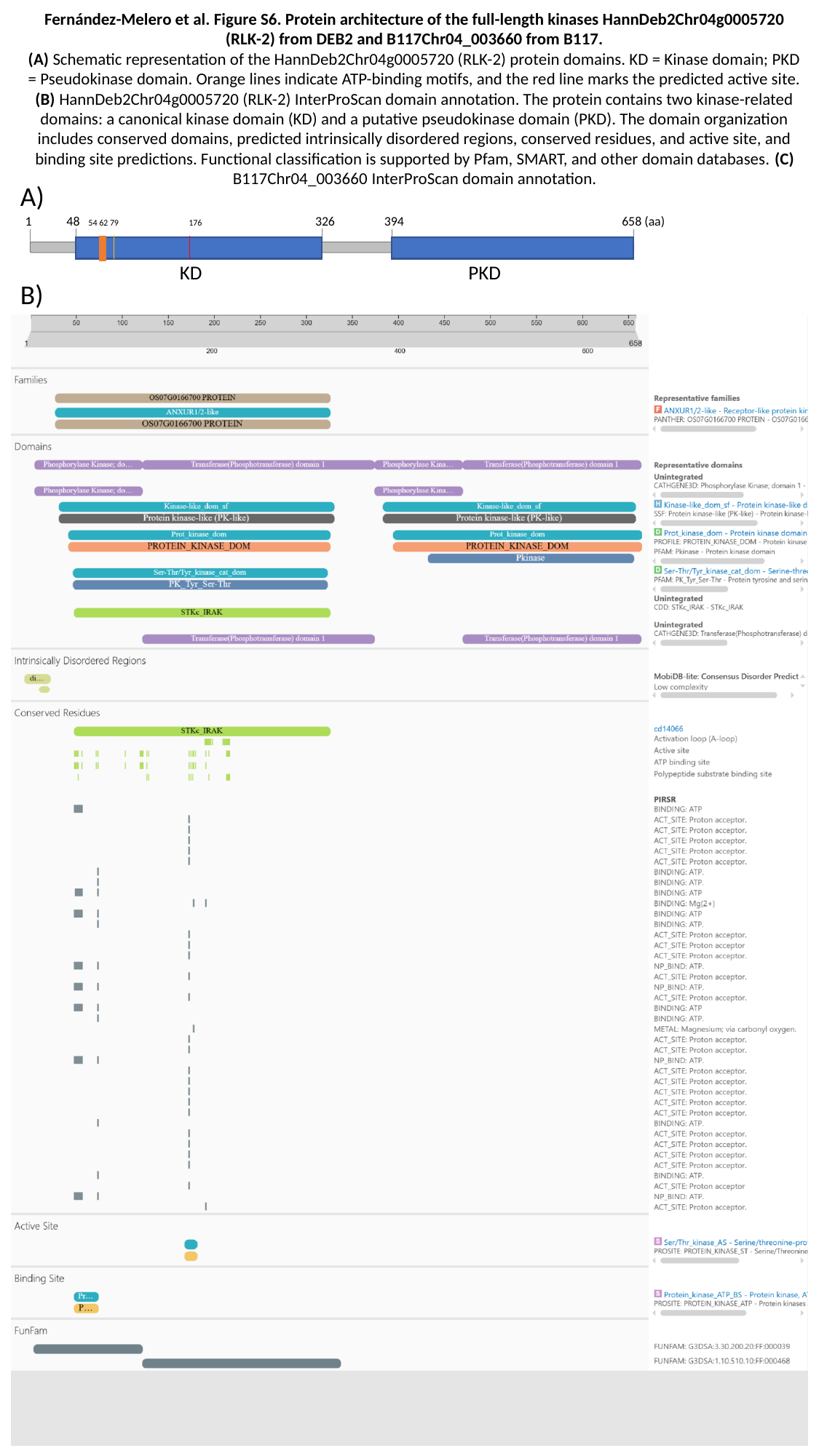

Fernández-Melero et al. Figure S6. Protein architecture of the full-length kinases HannDeb2Chr04g0005720 (RLK-2) from DEB2 and B117Chr04_003660 from B117.
(A) Schematic representation of the HannDeb2Chr04g0005720 (RLK-2) protein domains. KD = Kinase domain; PKD = Pseudokinase domain. Orange lines indicate ATP-binding motifs, and the red line marks the predicted active site.
(B) HannDeb2Chr04g0005720 (RLK-2) InterProScan domain annotation. The protein contains two kinase-related domains: a canonical kinase domain (KD) and a putative pseudokinase domain (PKD). The domain organization includes conserved domains, predicted intrinsically disordered regions, conserved residues, and active site, and binding site predictions. Functional classification is supported by Pfam, SMART, and other domain databases. (C) B117Chr04_003660 InterProScan domain annotation.
A)
B)
1 48 54 62 79 176 326 394 658 (aa)
KD PKD

### Slide 2
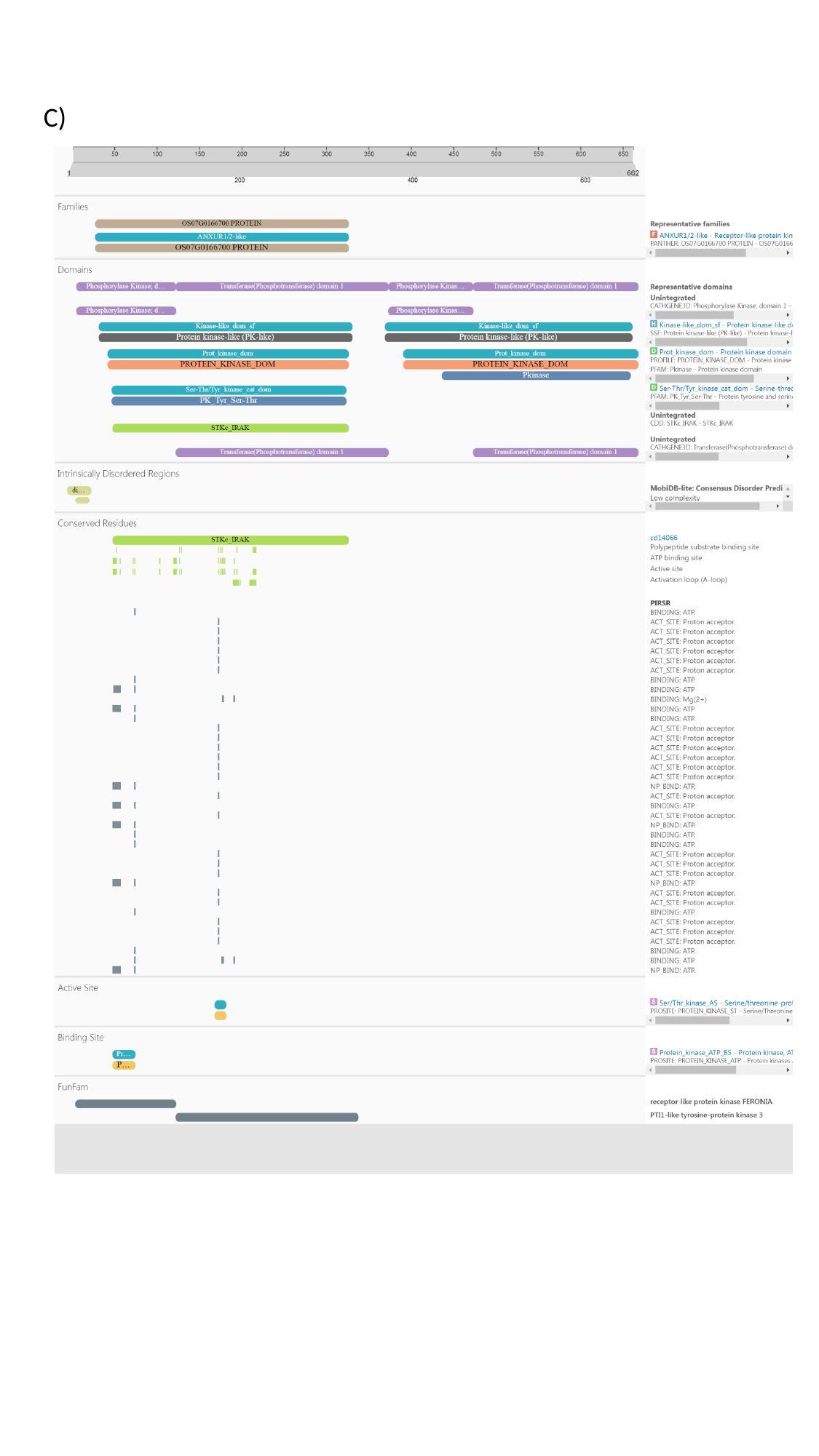

C)
