## Supplemental Figure S7 for "Fine mapping and genomic analyses reveal a tandem kinase-pseudokinase candidate underlying *Or_Deb2_*-mediated resistance to *Orobanche cumana* in sunflower"

### Slide 1
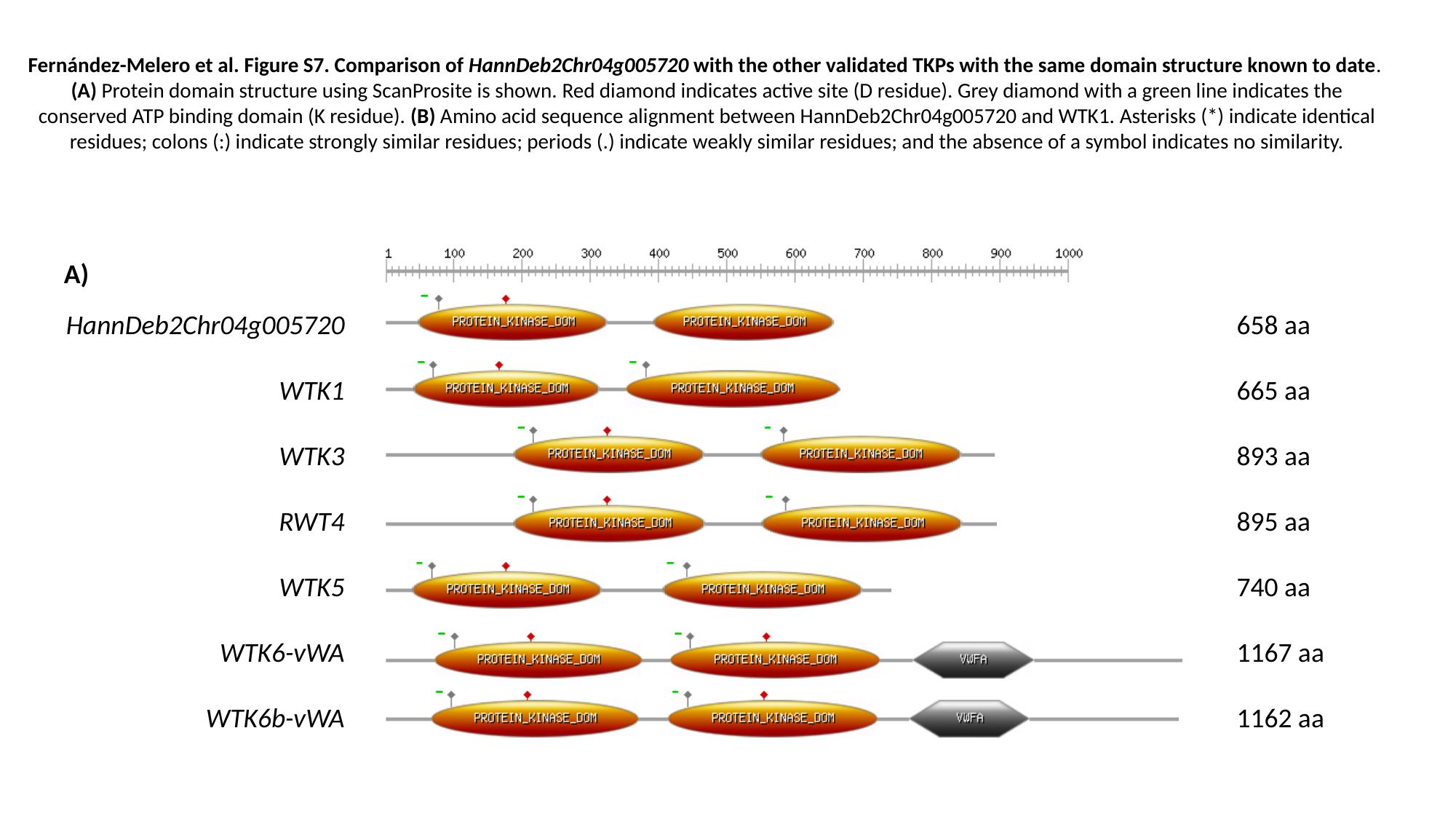

Fernández-Melero et al. Figure S7. Comparison of HannDeb2Chr04g005720 with the other validated TKPs with the same domain structure known to date.
(A) Protein domain structure using ScanProsite is shown. Red diamond indicates active site (D residue). Grey diamond with a green line indicates the conserved ATP binding domain (K residue). (B) Amino acid sequence alignment between HannDeb2Chr04g005720 and WTK1. Asterisks (*) indicate identical residues; colons (:) indicate strongly similar residues; periods (.) indicate weakly similar residues; and the absence of a symbol indicates no similarity.
A)
658 aa
665 aa
893 aa
895 aa
740 aa
1167 aa
1162 aa
HannDeb2Chr04g005720
WTK1
WTK3
RWT4
WTK5
WTK6-vWA
WTK6b-vWA

### Slide 2
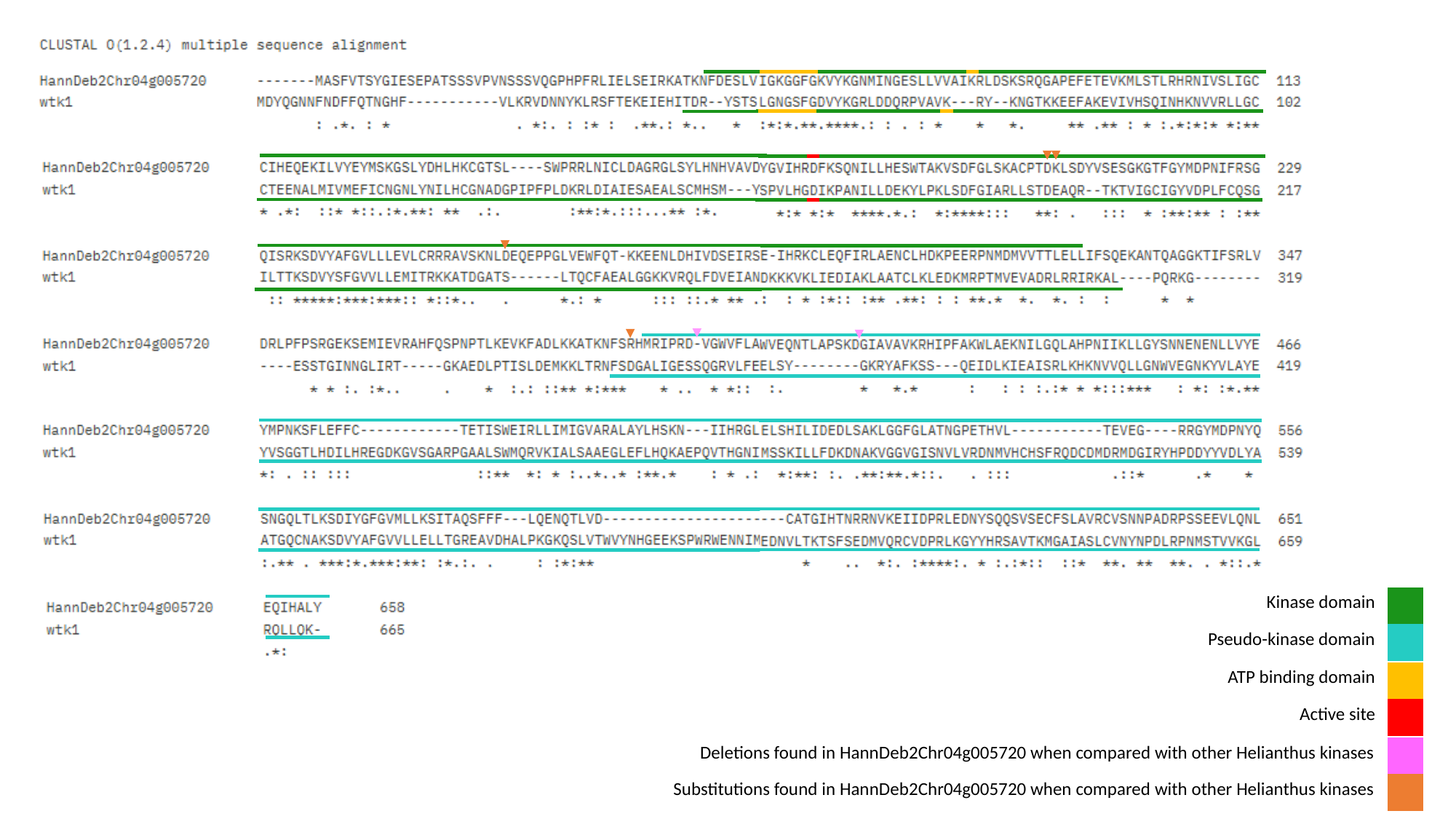

| Kinase domain |
| --- |
| Pseudo-kinase domain |
| |
| --- |
| ATP binding domain |
| --- |
| Active site |
| |
| --- |
| Deletions found in HannDeb2Chr04g005720 when compared with other Helianthus kinases |
| --- |
| Substitutions found in HannDeb2Chr04g005720 when compared with other Helianthus kinases |
| |
| --- |
