## Supplemental Figure S8 for "Fine mapping and genomic analyses reveal a tandem kinase-pseudokinase candidate underlying *Or_Deb2_*-mediated resistance to *Orobanche cumana* in sunflower"

### Slide 1
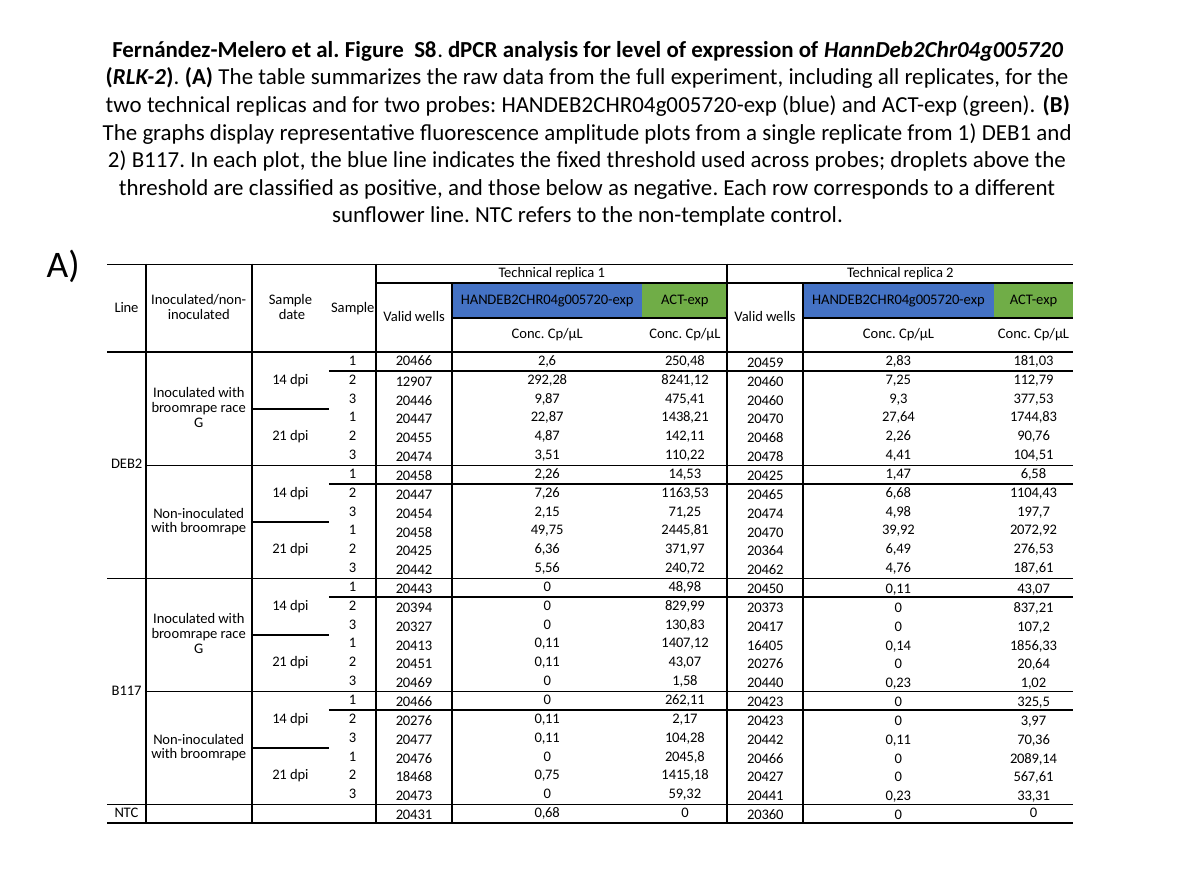

#### Fernández-Melero et al. Figure S8. dPCR analysis for level of expression of HannDeb2Chr04g005720 (RLK-2). (A) The table summarizes the raw data from the full experiment, including all replicates, for the two technical replicas and for two probes: HANDEB2CHR04g005720-exp (blue) and ACT-exp (green). (B) The graphs display representative fluorescence amplitude plots from a single replicate from 1) DEB1 and 2) B117. In each plot, the blue line indicates the fixed threshold used across probes; droplets above the threshold are classified as positive, and those below as negative. Each row corresponds to a different sunflower line. NTC refers to the non-template control.
A)
| Line | Inoculated/non-inoculated | Sample date | Sample | Technical replica 1 | | | Technical replica 2 | | |
| --- | --- | --- | --- | --- | --- | --- | --- | --- | --- |
| | | | | Valid wells | HANDEB2CHR04g005720-exp | ACT-exp | Valid wells | HANDEB2CHR04g005720-exp | ACT-exp |
| | | | | | Conc. Cp/μL | Conc. Cp/μL | | Conc. Cp/μL | Conc. Cp/μL |
| DEB2 | Inoculated with broomrape race G | 14 dpi | 1 | 20466 | 2,6 | 250,48 | 20459 | 2,83 | 181,03 |
| | | | 2 | 12907 | 292,28 | 8241,12 | 20460 | 7,25 | 112,79 |
| | | | 3 | 20446 | 9,87 | 475,41 | 20460 | 9,3 | 377,53 |
| | | 21 dpi | 1 | 20447 | 22,87 | 1438,21 | 20470 | 27,64 | 1744,83 |
| | | | 2 | 20455 | 4,87 | 142,11 | 20468 | 2,26 | 90,76 |
| | | | 3 | 20474 | 3,51 | 110,22 | 20478 | 4,41 | 104,51 |
| | Non-inoculated with broomrape | 14 dpi | 1 | 20458 | 2,26 | 14,53 | 20425 | 1,47 | 6,58 |
| | | | 2 | 20447 | 7,26 | 1163,53 | 20465 | 6,68 | 1104,43 |
| | | | 3 | 20454 | 2,15 | 71,25 | 20474 | 4,98 | 197,7 |
| | | 21 dpi | 1 | 20458 | 49,75 | 2445,81 | 20470 | 39,92 | 2072,92 |
| | | | 2 | 20425 | 6,36 | 371,97 | 20364 | 6,49 | 276,53 |
| | | | 3 | 20442 | 5,56 | 240,72 | 20462 | 4,76 | 187,61 |
| B117 | Inoculated with broomrape race G | 14 dpi | 1 | 20443 | 0 | 48,98 | 20450 | 0,11 | 43,07 |
| | | | 2 | 20394 | 0 | 829,99 | 20373 | 0 | 837,21 |
| | | | 3 | 20327 | 0 | 130,83 | 20417 | 0 | 107,2 |
| | | 21 dpi | 1 | 20413 | 0,11 | 1407,12 | 16405 | 0,14 | 1856,33 |
| | | | 2 | 20451 | 0,11 | 43,07 | 20276 | 0 | 20,64 |
| | | | 3 | 20469 | 0 | 1,58 | 20440 | 0,23 | 1,02 |
| | Non-inoculated with broomrape | 14 dpi | 1 | 20466 | 0 | 262,11 | 20423 | 0 | 325,5 |
| | | | 2 | 20276 | 0,11 | 2,17 | 20423 | 0 | 3,97 |
| | | | 3 | 20477 | 0,11 | 104,28 | 20442 | 0,11 | 70,36 |
| | | 21 dpi | 1 | 20476 | 0 | 2045,8 | 20466 | 0 | 2089,14 |
| | | | 2 | 18468 | 0,75 | 1415,18 | 20427 | 0 | 567,61 |
| | | | 3 | 20473 | 0 | 59,32 | 20441 | 0,23 | 33,31 |
| NTC | | | | 20431 | 0,68 | 0 | 20360 | 0 | 0 |

### Slide 2
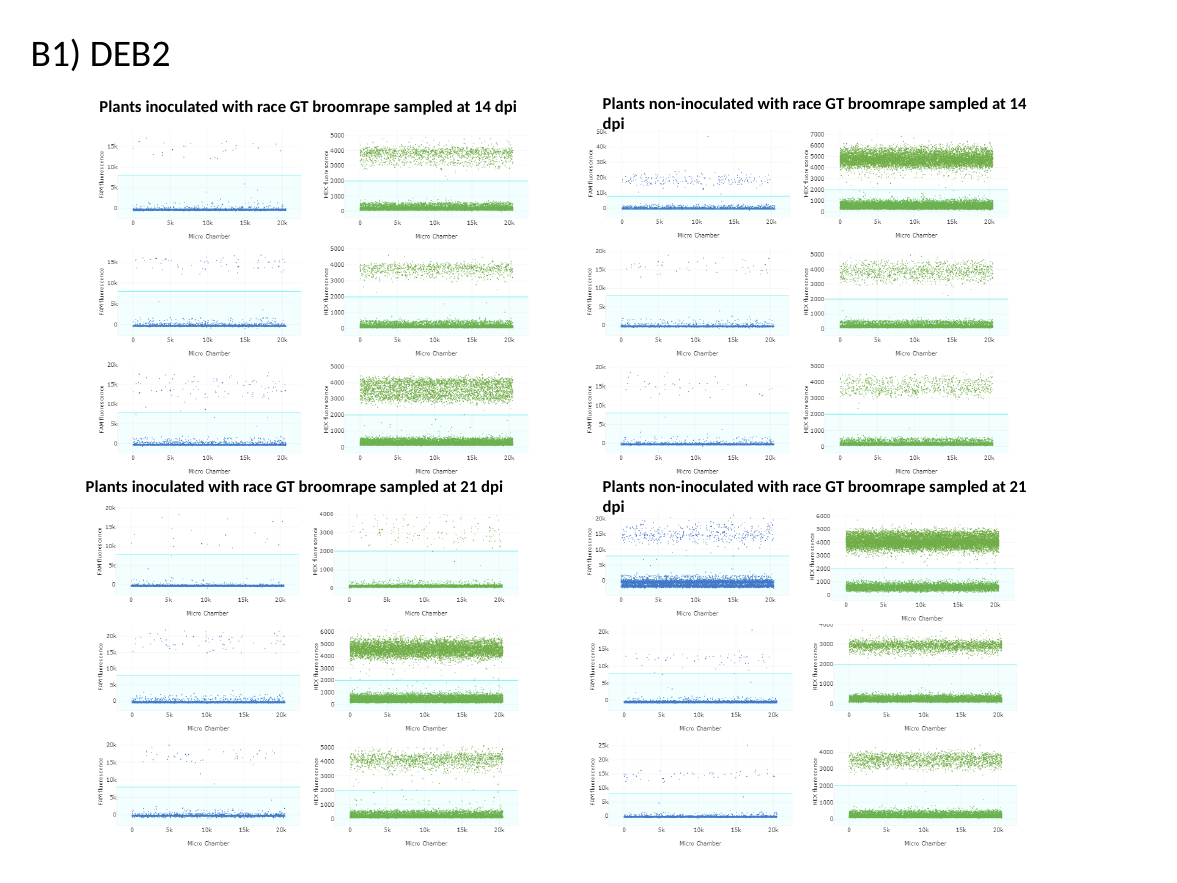

Plants non-inoculated with race GT broomrape sampled at 14 dpi
Plants inoculated with race GT broomrape sampled at 14 dpi
B1) DEB2
Plants non-inoculated with race GT broomrape sampled at 21 dpi
Plants inoculated with race GT broomrape sampled at 21 dpi

### Slide 3
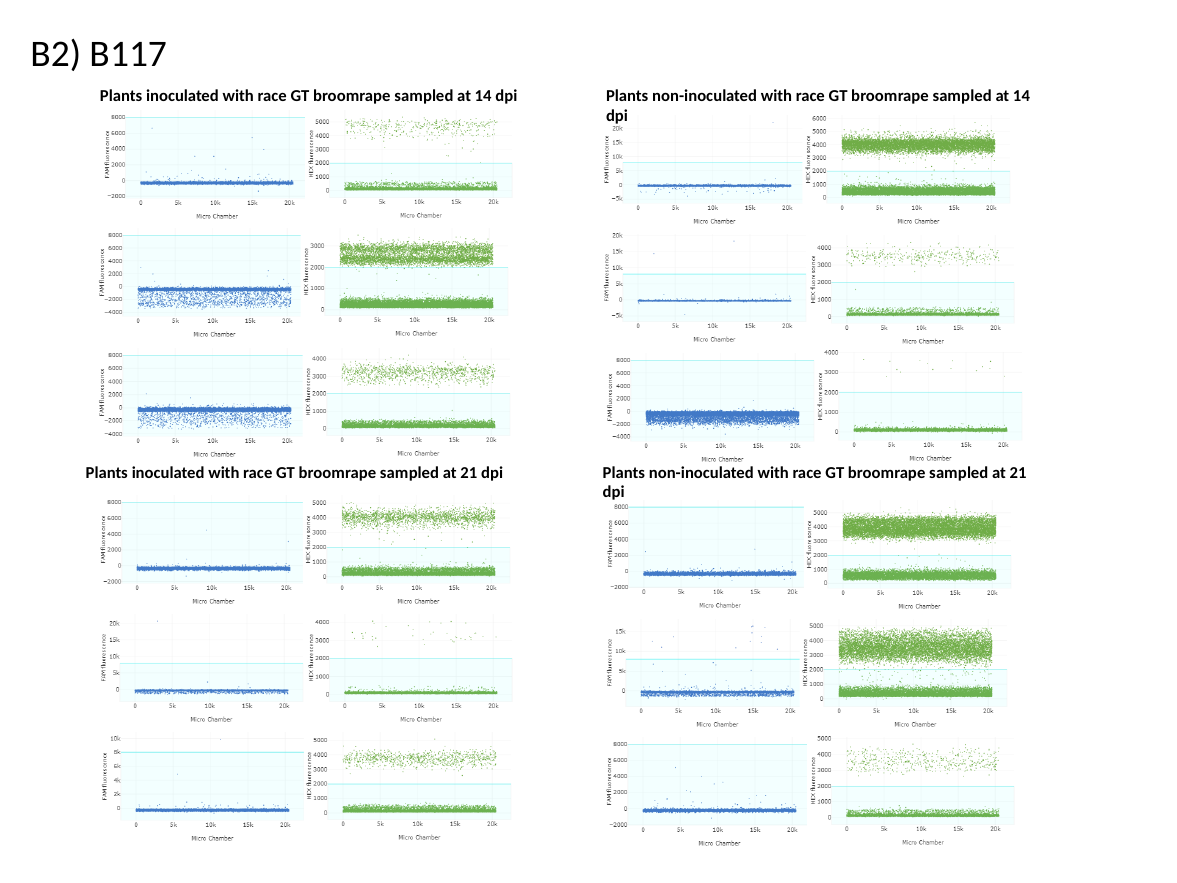

B2) B117
Plants inoculated with race GT broomrape sampled at 14 dpi
Plants non-inoculated with race GT broomrape sampled at 14 dpi
Plants non-inoculated with race GT broomrape sampled at 21 dpi
Plants inoculated with race GT broomrape sampled at 21 dpi
