## Supplemental Figure S9 for "Fine mapping and genomic analyses reveal a tandem kinase-pseudokinase candidate underlying *Or_Deb2_*-mediated resistance to *Orobanche cumana* in sunflower"

### Slide 1
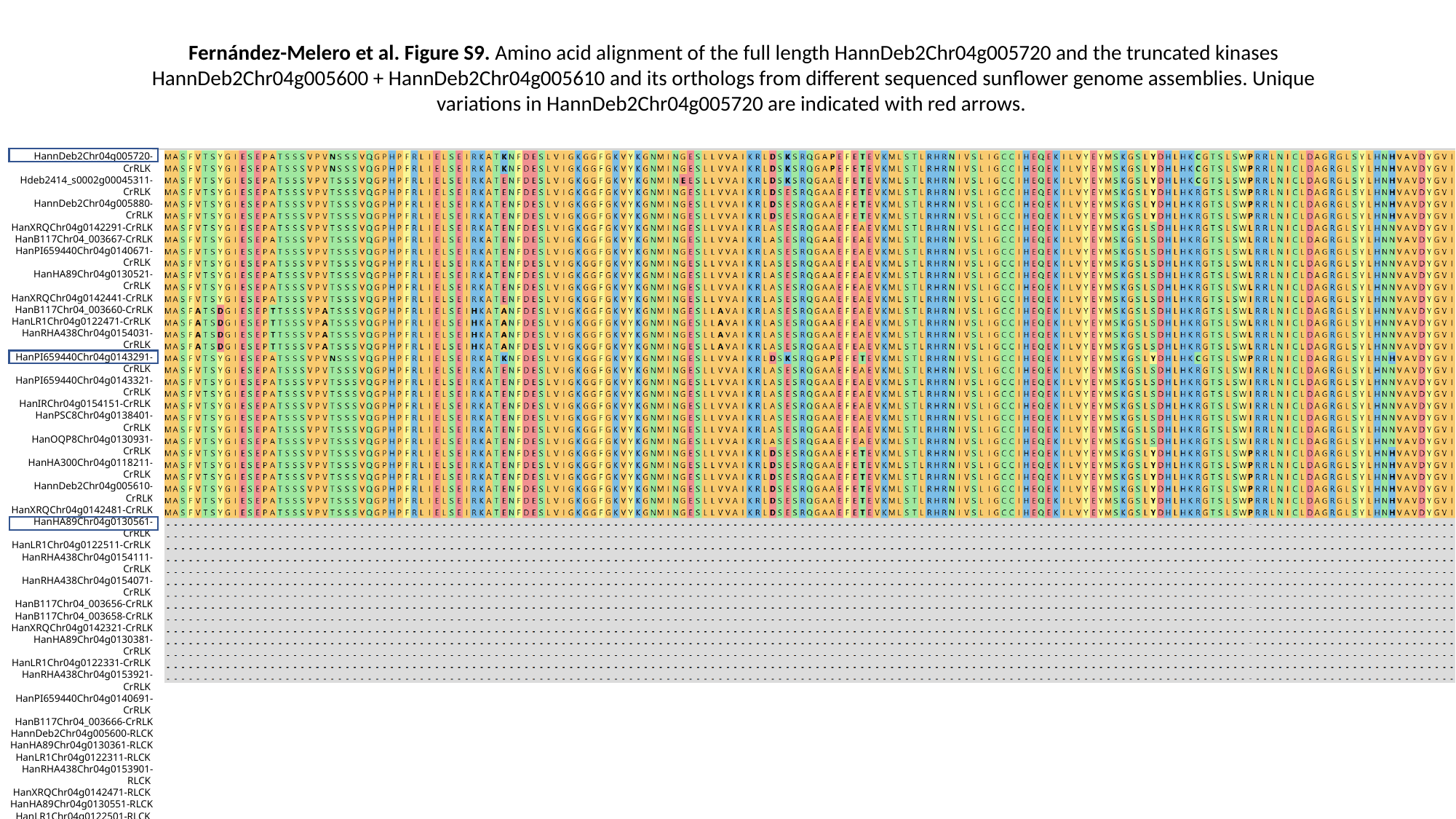

Fernández-Melero et al. Figure S9. Amino acid alignment of the full length HannDeb2Chr04g005720 and the truncated kinases HannDeb2Chr04g005600 + HannDeb2Chr04g005610 and its orthologs from different sequenced sunflower genome assemblies. Unique variations in HannDeb2Chr04g005720 are indicated with red arrows.
HannDeb2Chr04g005720-CrRLK
Hdeb2414_s0002g00045311-CrRLK
HannDeb2Chr04g005880-CrRLK
HanXRQChr04g0142291-CrRLK
HanB117Chr04_003667-CrRLK
HanPI659440Chr04g0140671-CrRLK
HanHA89Chr04g0130521-CrRLK
HanXRQChr04g0142441-CrRLK
HanB117Chr04_003660-CrRLK
HanLR1Chr04g0122471-CrRLK
HanRHA438Chr04g0154031-CrRLK
HanPI659440Chr04g0143291-CrRLK
HanPI659440Chr04g0143321-CrRLK
HanIRChr04g0154151-CrRLK
HanPSC8Chr04g0138401-CrRLK
HanOQP8Chr04g0130931-CrRLK
HanHA300Chr04g0118211-CrRLK
HannDeb2Chr04g005610-CrRLK
HanXRQChr04g0142481-CrRLK
HanHA89Chr04g0130561-CrRLK
HanLR1Chr04g0122511-CrRLK
HanRHA438Chr04g0154111-CrRLK
HanRHA438Chr04g0154071-CrRLK
HanB117Chr04_003656-CrRLK
HanB117Chr04_003658-CrRLK
HanXRQChr04g0142321-CrRLK
HanHA89Chr04g0130381-CrRLK
HanLR1Chr04g0122331-CrRLK
HanRHA438Chr04g0153921-CrRLK
HanPI659440Chr04g0140691-CrRLK
HanB117Chr04_003666-CrRLK
HannDeb2Chr04g005600-RLCK
HanHA89Chr04g0130361-RLCK
HanLR1Chr04g0122311-RLCK
HanRHA438Chr04g0153901-RLCK
HanXRQChr04g0142471-RLCK
HanHA89Chr04g0130551-RLCK
HanLR1Chr04g0122501-RLCK
HanRHA438Chr04g0154061-RLCK
HanRHA438Chr04g0154101-RLCK
HanB117Chr04_003657-RLCK
HanB117Chr04_003659-RLCK
Hdeb2414_s0002g00045341-RLCK
Hanom_Chr07g00585851-CrRLK
Hanom_Chr07g00585861-RLCK

### Slide 2
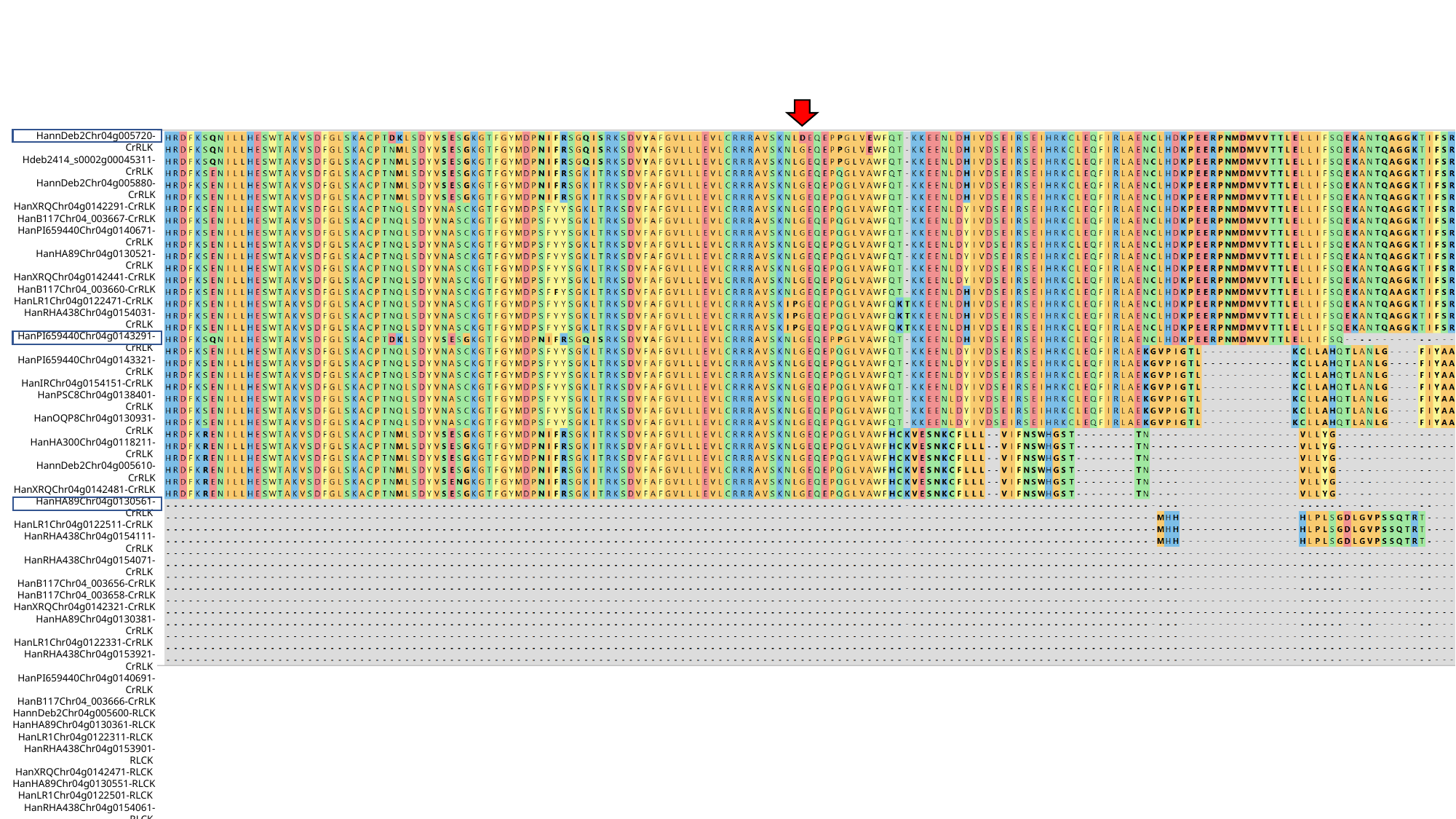

HannDeb2Chr04g005720-CrRLK
Hdeb2414_s0002g00045311-CrRLK
HannDeb2Chr04g005880-CrRLK
HanXRQChr04g0142291-CrRLK
HanB117Chr04_003667-CrRLK
HanPI659440Chr04g0140671-CrRLK
HanHA89Chr04g0130521-CrRLK
HanXRQChr04g0142441-CrRLK
HanB117Chr04_003660-CrRLK
HanLR1Chr04g0122471-CrRLK
HanRHA438Chr04g0154031-CrRLK
HanPI659440Chr04g0143291-CrRLK
HanPI659440Chr04g0143321-CrRLK
HanIRChr04g0154151-CrRLK
HanPSC8Chr04g0138401-CrRLK
HanOQP8Chr04g0130931-CrRLK
HanHA300Chr04g0118211-CrRLK
HannDeb2Chr04g005610-CrRLK
HanXRQChr04g0142481-CrRLK
HanHA89Chr04g0130561-CrRLK
HanLR1Chr04g0122511-CrRLK
HanRHA438Chr04g0154111-CrRLK
HanRHA438Chr04g0154071-CrRLK
HanB117Chr04_003656-CrRLK
HanB117Chr04_003658-CrRLK
HanXRQChr04g0142321-CrRLK
HanHA89Chr04g0130381-CrRLK
HanLR1Chr04g0122331-CrRLK
HanRHA438Chr04g0153921-CrRLK
HanPI659440Chr04g0140691-CrRLK
HanB117Chr04_003666-CrRLK
HannDeb2Chr04g005600-RLCK
HanHA89Chr04g0130361-RLCK
HanLR1Chr04g0122311-RLCK
HanRHA438Chr04g0153901-RLCK
HanXRQChr04g0142471-RLCK
HanHA89Chr04g0130551-RLCK
HanLR1Chr04g0122501-RLCK
HanRHA438Chr04g0154061-RLCK
HanRHA438Chr04g0154101-RLCK
HanB117Chr04_003657-RLCK
HanB117Chr04_003659-RLCK
Hdeb2414_s0002g00045341-RLCK
Hanom_Chr07g00585851-CrRLK
Hanom_Chr07g00585861-RLCK

### Slide 3
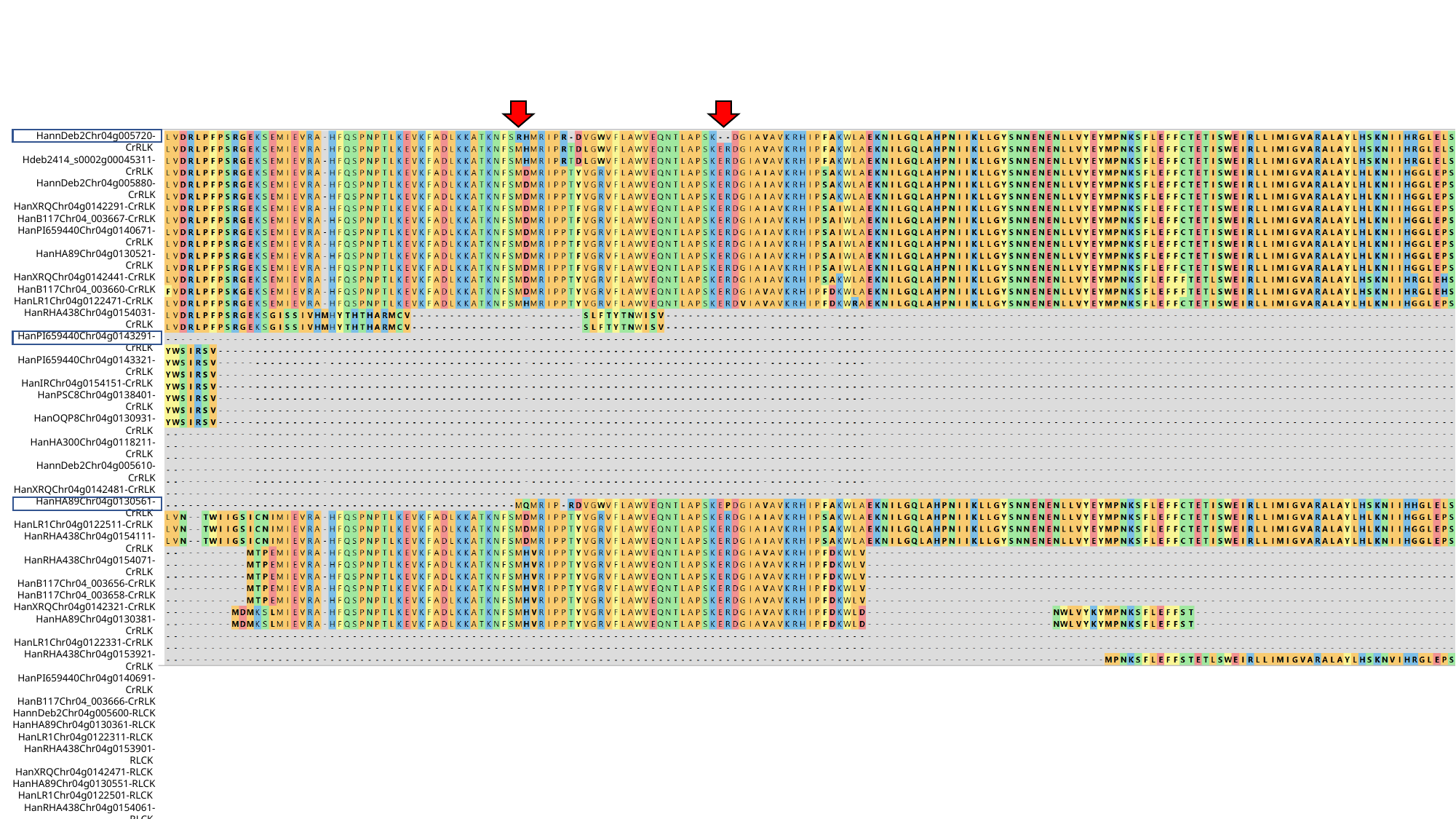

HannDeb2Chr04g005720-CrRLK
Hdeb2414_s0002g00045311-CrRLK
HannDeb2Chr04g005880-CrRLK
HanXRQChr04g0142291-CrRLK
HanB117Chr04_003667-CrRLK
HanPI659440Chr04g0140671-CrRLK
HanHA89Chr04g0130521-CrRLK
HanXRQChr04g0142441-CrRLK
HanB117Chr04_003660-CrRLK
HanLR1Chr04g0122471-CrRLK
HanRHA438Chr04g0154031-CrRLK
HanPI659440Chr04g0143291-CrRLK
HanPI659440Chr04g0143321-CrRLK
HanIRChr04g0154151-CrRLK
HanPSC8Chr04g0138401-CrRLK
HanOQP8Chr04g0130931-CrRLK
HanHA300Chr04g0118211-CrRLK
HannDeb2Chr04g005610-CrRLK
HanXRQChr04g0142481-CrRLK
HanHA89Chr04g0130561-CrRLK
HanLR1Chr04g0122511-CrRLK
HanRHA438Chr04g0154111-CrRLK
HanRHA438Chr04g0154071-CrRLK
HanB117Chr04_003656-CrRLK
HanB117Chr04_003658-CrRLK
HanXRQChr04g0142321-CrRLK
HanHA89Chr04g0130381-CrRLK
HanLR1Chr04g0122331-CrRLK
HanRHA438Chr04g0153921-CrRLK
HanPI659440Chr04g0140691-CrRLK
HanB117Chr04_003666-CrRLK
HannDeb2Chr04g005600-RLCK
HanHA89Chr04g0130361-RLCK
HanLR1Chr04g0122311-RLCK
HanRHA438Chr04g0153901-RLCK
HanXRQChr04g0142471-RLCK
HanHA89Chr04g0130551-RLCK
HanLR1Chr04g0122501-RLCK
HanRHA438Chr04g0154061-RLCK
HanRHA438Chr04g0154101-RLCK
HanB117Chr04_003657-RLCK
HanB117Chr04_003659-RLCK
Hdeb2414_s0002g00045341-RLCK
Hanom_Chr07g00585851-CrRLK
Hanom_Chr07g00585861-RLCK

### Slide 4
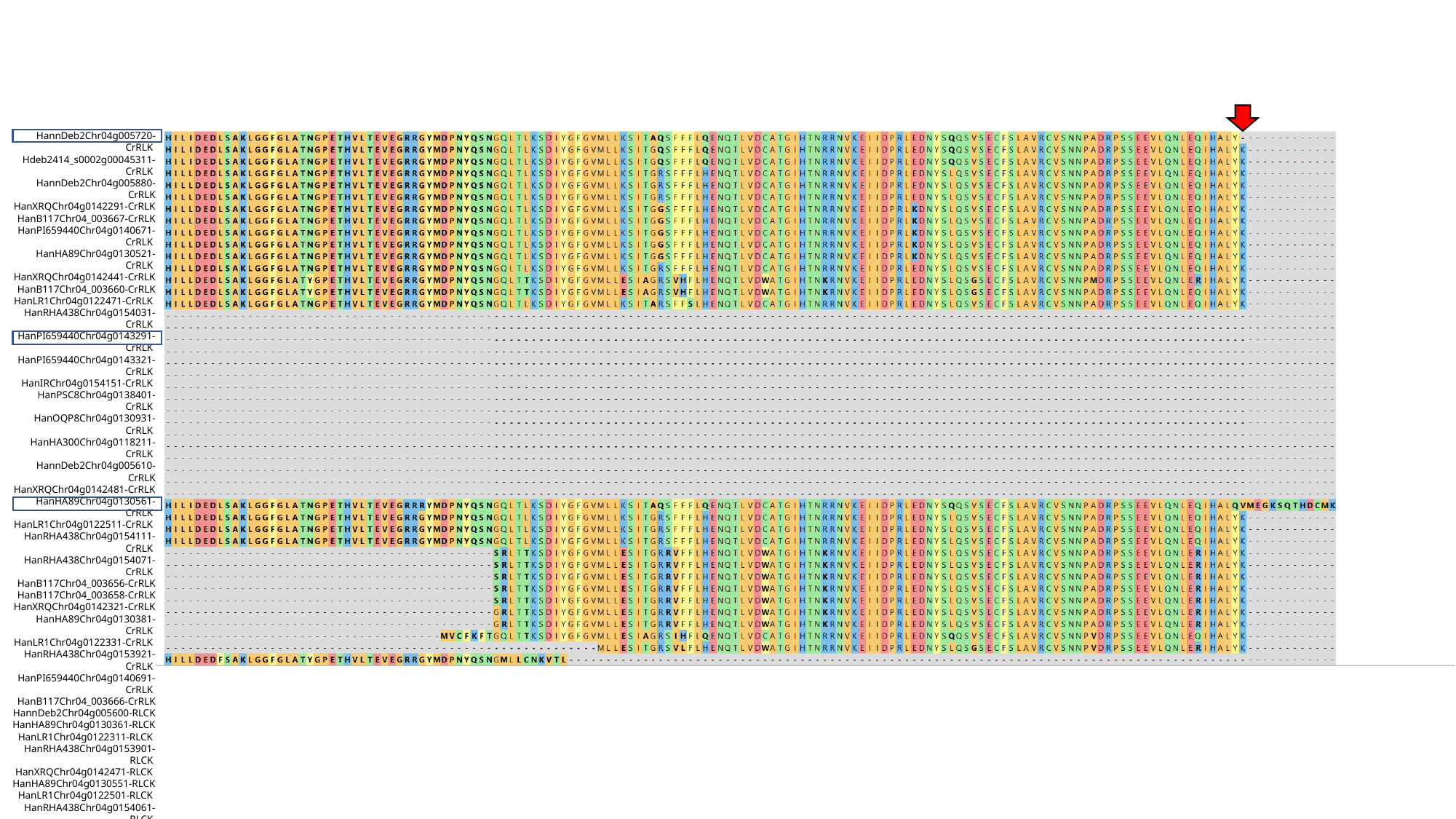

HannDeb2Chr04g005720-CrRLK
Hdeb2414_s0002g00045311-CrRLK
HannDeb2Chr04g005880-CrRLK
HanXRQChr04g0142291-CrRLK
HanB117Chr04_003667-CrRLK
HanPI659440Chr04g0140671-CrRLK
HanHA89Chr04g0130521-CrRLK
HanXRQChr04g0142441-CrRLK
HanB117Chr04_003660-CrRLK
HanLR1Chr04g0122471-CrRLK
HanRHA438Chr04g0154031-CrRLK
HanPI659440Chr04g0143291-CrRLK
HanPI659440Chr04g0143321-CrRLK
HanIRChr04g0154151-CrRLK
HanPSC8Chr04g0138401-CrRLK
HanOQP8Chr04g0130931-CrRLK
HanHA300Chr04g0118211-CrRLK
HannDeb2Chr04g005610-CrRLK
HanXRQChr04g0142481-CrRLK
HanHA89Chr04g0130561-CrRLK
HanLR1Chr04g0122511-CrRLK
HanRHA438Chr04g0154111-CrRLK
HanRHA438Chr04g0154071-CrRLK
HanB117Chr04_003656-CrRLK
HanB117Chr04_003658-CrRLK
HanXRQChr04g0142321-CrRLK
HanHA89Chr04g0130381-CrRLK
HanLR1Chr04g0122331-CrRLK
HanRHA438Chr04g0153921-CrRLK
HanPI659440Chr04g0140691-CrRLK
HanB117Chr04_003666-CrRLK
HannDeb2Chr04g005600-RLCK
HanHA89Chr04g0130361-RLCK
HanLR1Chr04g0122311-RLCK
HanRHA438Chr04g0153901-RLCK
HanXRQChr04g0142471-RLCK
HanHA89Chr04g0130551-RLCK
HanLR1Chr04g0122501-RLCK
HanRHA438Chr04g0154061-RLCK
HanRHA438Chr04g0154101-RLCK
HanB117Chr04_003657-RLCK
HanB117Chr04_003659-RLCK
Hdeb2414_s0002g00045341-RLCK
Hanom_Chr07g00585851-CrRLK
Hanom_Chr07g00585861-RLCK
